# A dual genetic constraint underlies the conservation of early brains in vertebrates

**DOI:** 10.1101/2025.10.29.684766

**Authors:** Rodrigo Senovilla-Ganzo, Christina Bekiari, Eneritz Rueda-Alaña, Tetsuya Yamada, Bastienne Zaremba, Ana María Aransay, Laura Escobar, Mats Nilsson, Marco Grillo, Henrik Kaessmann, Fernando García-Moreno

**Author notes:** CIRB, Collège de France, 11 Pl. Marcelin Berthelot, 75231 Paris France.

## Abstract

As the body plan, the embryonic brain bauplan reflects the shared features of vertebrate brains. Yet, disagreements among sparse histogenetic frameworks have undermined the bauplan’s power to trace homologies. Here, we generate and integrate five vertebrate single-cell multi-omic atlases of early embryonic brains, revealing a conserved cellular blueprint that defines equivalent progenitor domains across species. Our cellular neuromeric model provides an unified and unbiased framework for the vertebrate brain bauplan and revises the regionalisation of the prosencephalon, refining its molecular boundaries and developmental relationships. Furthermore, cross-species gene-network analyses expose regulatory complexity beyond classical neuromeric patterning, resolving networks into modules aligned with regional cell types or cell class (progenitor/neuron). Evolutionarily, two main developmental constraints emerge: brain bauplan genes, early essential for regional identity, and pleiotropic stemness gene modules, indispensable across all proliferating cells. In turn, later development displays tissue-specific and less essential modules, explaining its rapid divergence into species-specific features. Together, these findings reveal a dual evolutionary constraint—neuromeric identity and pleiotropic stemness—that underlies the conservation of early vertebrate brains. This duality explains how deeply conserved regulatory architectures coexist with evolutionary flexibility to develop into the immense diversity of vertebrate nervous systems.

**Graphical abstract:** 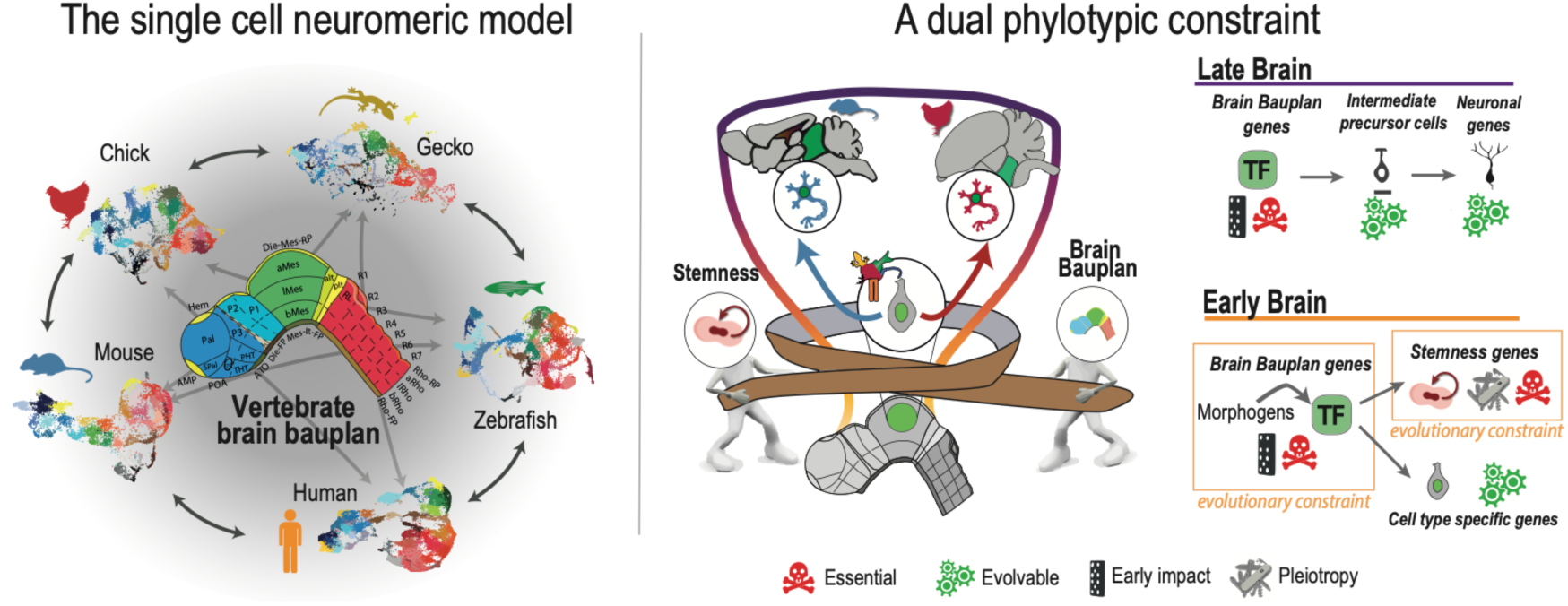

## Introduction

Just as the vertebrate body plan refers to the shared anatomical features across all vertebrate species^1–5^, the brain plan -or *bauplan-* is hypothesized to reflect a conserved organizational architecture of vertebrate brains^6,7^. As early as the 19^th^ century, von Baer^5^ and other early neuroanatomists^8,9^ had already illustrated how brain primordia are common vertebrate hallmarks, despite their later divergence into diverse adult structures and circuits^10,11^. These early morphological similarities laid the foundation for identifying evolutionary homologies between disparate adult structures (e.g., the mammalian superior colliculus vs. the avian optic tectum^12^) (**Fig. 1a**), as well as for tracing evolutionarily related adult regions, such as the telencephalon and hypothalamus, back to common developmental origins^8,13^. How these structures and homologies were preserved over hundreds of millions of years of evolution remains unresolved.

**Fig. 1.**
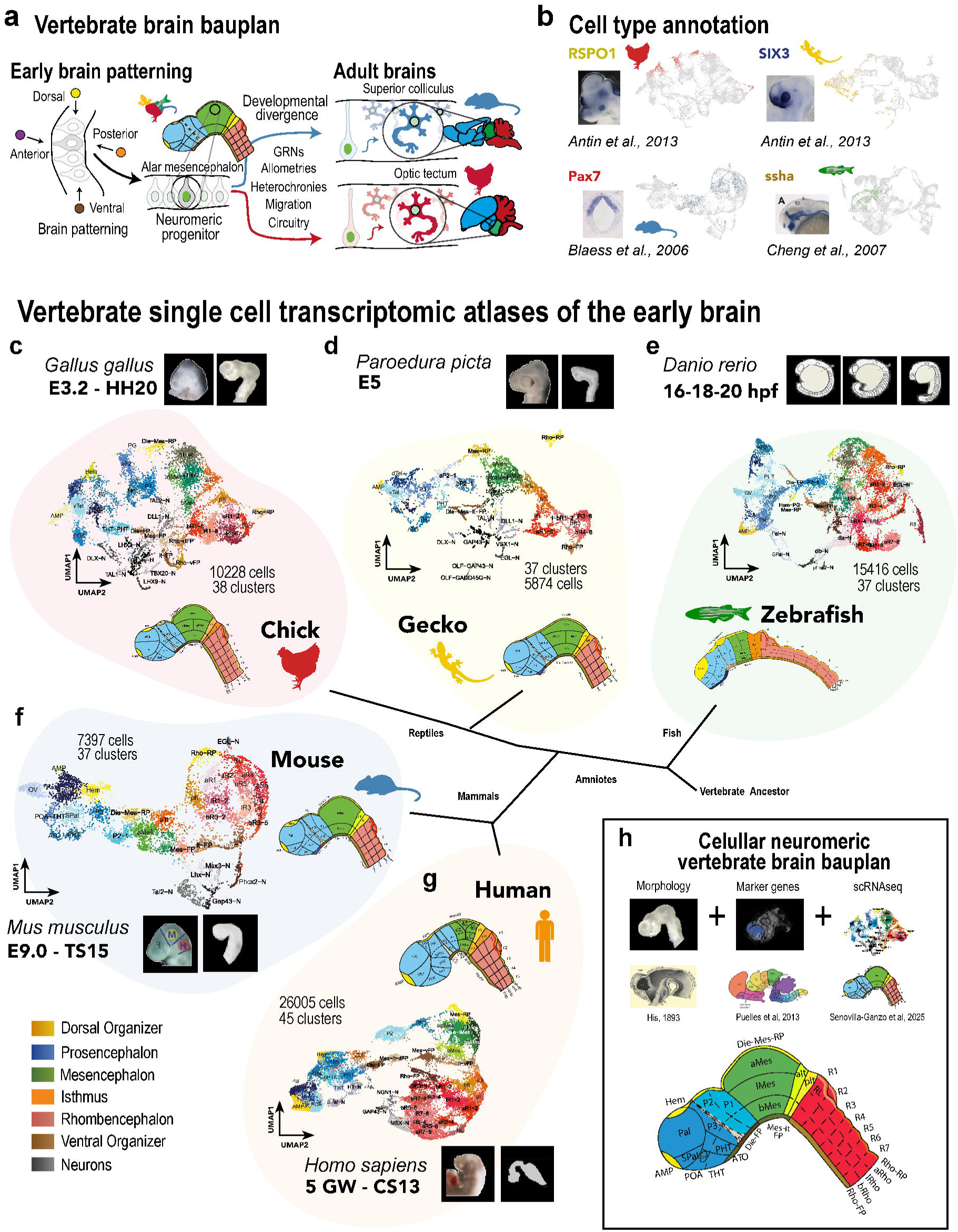
The cellular brain bauplan: Five vertebrate single cell transcriptomic atlases. **a**, The molecular patterning of the early brain into the brain bauplan and its later developmental and evolutionary divergence. **b**, Cell type example annotation based on available histological data on literature and databases^41,124–127^ (left) and single cell expression plots (right). **c-g**, Single cell atlases and neuroanatomical representation of the early brain bauplans from chick (**c**), gecko (**d**), zebrafish (**e**), mouse (**f**) and human (**g**). Each species’s head and its subsequent dissection is visualized on black squares close to its biological name. **h**, Cellular neuromeric foundations and its cell type neuroanatomical distribution. Color code is differentiated per brain region and species: blue (prosencephalon and mouse), green (mesencephalon and zebrafish), red (rhombencephalon and chick), gold (dorsal organizers and gecko), orange (isthmus and human), brown (ventral organizers) and grey tonalities (neurons). Continuous lines in neuroanatomical schemes indicate confidence over discontinuous lines.

With the combination of genetics and histological tools (e.g., in situ hybridization), an evolutionarily conserved array of morphogens and transcription factors (TFs) was identified as the basis of the vertebrate brain bauplan and its segmental units, the neuromeres^8,9^. This phenomenon extends beyond the brain: the entire body plan was shown to rely on deeply conserved patterning regulators, such as the *HOX* genes^14,15^. Building on von Baer’s laws⁵, these insights gave rise to the hourglass model^16,17^, which highlights the high conservation of mid-embryonic, or *phylotypic*, stages in contrast to the more variable early and late stages of development^18,19^. However, the limited power of available tools could not untangle the causes behind such conservation.

Nowadays, -*omic* approaches have not only demonstrated strong transcriptomic conservation during vertebrate mid-embryogenesis at the whole-embryo bulk level^20^, but also revealed extensive pleiotropy and developmental interdependence among organs^21,22^. These are features hypothesized to render these stages essential and constrain their evolutionary divergence^23,24^. However, as these omics comparisons are based on whole-embryo bulk data, they cannot specifically address cell-type composition or transcriptomic evolutionary conservation^25,26^. These technical constraints obscure the specific evolutionary contributions of distinct cell type signatures, as well as the underlying complexity of gene regulatory networks (GRNs) and developmental programs. This ultimately limits the interpretability of mean scores and masks the evolutionary reasons for such conservation ^27–29^.

In this study, we investigate the early brain at the cellular and multi-omic levels as a proxy for the brain bauplan cell-type architecture and to resolve the molecular mechanisms underlying the phylotypic evolutionary constraint. First, our single-cell RNA-seq (scRNA-seq) comparisons across five vertebrate early brains reveal that the neuromeric cellular organization of the brain bauplan is evolutionarily conserved, revealing high cell-type transcriptomic similarity beyond previous marker-gene-based comparisons. Complementarily, our single-nucleus ATAC-seq (snATAC-seq) distinguish a deeply conserved set of brain bauplan TFs driving neuromeric cell types and lineages, highly valuable to unravel homologies across vertebrate structures. Together with our gene co-expression network analysis, snATAC-seq also resolves the existence of a secondary set of TFs and gene modules that are non-brain-specific. These are highly conserved gene modules related to stemness and are expressed across all body tissues and stages, also constraining the evolutionary variability of the early brain landscape due to their high pleiotropy and essentiality. Later in development, the brain undergoes successive replacement of these essential pleiotropic modules by non-essential brain-specific modules -such as synapsis or ciliary machinery. This maturation switch might involve the release from phylotypic constraints, promoting developmental variability subject to natural selection.

Overall, the early brain represents a paradigmatic case of dual, intertwined evolutionary constraints, independently driven by brain bauplan-specific and essential pleiotropic stemness genes. This duality provides a valuable paradigm on how evolution acts differently on specific elements of brain developmental GRNs, and potentially across all organs within the body plan.

## Results

### The brain bauplan conservation at the cellular level

Starting at gastrulation, an additive series of morphogens shapes every organ axis and cell type lineage in vertebrates by inducing a cardinal, region-specific code of TFs (**Fig. 1a**). These tightly preserved ancestral identities allows evolutionary biologists to trace homologous identities despite later divergence in adulthood^30–32^. While bird wings and human arms are a paradigmatic example of homology^30,33,34^, brain regions still remain a contentious subject regarding the molecular evidence needed to sustain their evolutionary relationships^35–37^. Current homology proposals are based on histogenetic approaches that yield heterogeneous, targeted, and low throughput for evolutionary comparisons^37^. Moreover, they lack cellular resolution, failing to capture the combinatorial nature of patterning genes across single cells. These limitations have consistently acted as a double-edged sword, bolstering models centered on a handful of selected master genes while dismissing competing proposals on the same grounds. This lack of consensus and cross-cutting evidence has weakened the brain bauplan homologies in favour of intuitive alternatives such as *functional homology*, which are highly prone to reflect convergent phenomena^11,31,36,38,39^.

To overcome these limitations, here we employ single-cell transcriptomics to systematically and unbiasedly determine the cellular composition of the vertebrate brain bauplan. For this purpose, five cellular neuromeric atlases (scRNA-seq) from representative vertebrate taxa were generated (**Fig. 1c-g**): reptiles, chicken (**Fig. S01**) and gecko (**Fig. S02**) (in-house data); mammals, mouse^40^ (**Fig. S03**) and human^41^ (**Fig. S01**); and fish, zebrafish^42^ (**Fig. S05**). Comparable stages were selected based on three criteria: recent neural tube closure, initial neurogenesis (indicator of an established bauplan) and high whole-embryo transcriptomic similarities^22^. Besides morphological similarities, the five vertebrate species characterized here show a common vertebrate catalogue of cell types and a conserved marker-gene dictionary that reflects a highly evolutionary constrained brain bauplan (**Fig. 1b; Table S01**). Despite these clusters are the result of non-guided transcriptome-based clusterization, these cell types clearly depict the brain bauplan along each axis of the neural tube.

This spatial information is differently encoded by different cell classes beyond regional patterning identities: neuroepithelial stem cells and neurons. Among the former, organizers (e.g. mesencephalic roof plate) segregate morphogens that are received by other neuroepithelial progenitors. The combination of positional cues turns these neuroepithelial cells into region or neuromere specific progenitors (e.g. alar mesencephalon), which are identifiable by their TF code. These progenitors will generate neurons in time-controlled neurogenic waves during development to carry out specific neuronal roles^43^. In our early atlases, the first wave of early neurons can be already observed (e.g. TAL mesencephalic neurons).

Beyond their classification into major cell classes, the five single-cell atlases primarily reflect the underlying embryonic patterning networks. Along the anterior-posterior (A–P) axis **—** visible from left to right in the five UMAPs**—** cells are organized according to their origin in major brain vesicles: the prosencephalon (adult forebrain, blue shades) occupies the anterior end, followed by the mesencephalon (adult midbrain, green shades), and the rhombencephalon (adult hindbrain, red shades) in the posterior. In parallel, morphogenetic gradients along the dorsal–ventral (D–V) axis shape a top-to-bottom organization: dorsal territories (yellow shades; e.g., *RSPO1*, *RSPO3*) give rise to alar plate identities, ventral organizers (brown shades; e.g., *SHH*, *FOXA1*) induce basal identities, and an intermediate ‘liminal’ domain emerges between them, all of which are represented in the vertical distribution of cell types across the five vertebrate atlases.

The additive and combinatorial effects of multiple morphogenetic gradients become more evident at higher-resolution clustering. At this finer level, sharply defined domains -such as the anterior basal mesencephalon, a fourth liminal rhombomere, or the prosencephalic cortical hem- are remarkably preserved across the five vertebrate species. Nonetheless, we also observe differences in the relative abundance of cells per cluster, the presence of patterning genes, and how broad neuromeres are subdivided into more or fewer D–V or A–P domains. Some of these differences are consistent across evolutionary lineages and may reflect genuine adaptations like an increased mesencephalic region in reptiles^44^. However, the apparent absence or low expression of certain genes (e.g. *FOXG1* in mouse) or subdomains (POA in mouse) in some clusters or species could rather stem from technical limitations such as single-cell sparsity^45,46^. Overall, our cellular neuromeric model (**Fig. 1h**) offers a robust cross-species framework that captures the shared architecture of the vertebrate brain bauplan. Our cellular neuromeric framework supports and extends the histogenetic prosomeric model^6^ of the brain bauplan, validating its core principles while providing additional resolution. A core principle in both models is the D-V delimitation of each A-P segment into distinct alar and basal domains. While there has always been consensus on this D-V delimitation for the mesencephalon and rhombencephalon^47,48^; it remains contentious in the case of the secondary prosencephalon. According to classical columnar theory^49^ and other current alternatives^50^, the hypothalamus and telencephalon are evolutionarily unrelated as they have different developmental origins. In contrast, our model posits that both adult structures arise from the same A-P segment or neuromere (P4). In particular, our single-cell analysis identifies *EMX2* and *FEZF2* as key TFs defining the whole P4 segment: its alar domain gives rise to the telencephalon, while its basal portion develops into the posterior peduncular hypothalamus (PHT) (**Fig. S06**). Recognizing this alar–basal organization within each A– P segment is crucial not only for interpreting transcriptomic similarities within species, but also for understanding the shared developmental origin and evolutionary history of these structures since the bilaterian divergence^51–53^ (**Fig. S07**).

On the other hand, our model also challenges some assumptions of the prosomeric framework. Puelles and collaborators’ model often extends the diencephalon rostrally beyond the zona limitans intrathalamica (ZLI), including prosomere 3 (P3) or prethalamus. In contrast, our cellular neuromeric model defines the secondary prosencephalon (blue shades, **Fig. 1h**) by early SIX3 expression^54^: encompassing the telencephalon, optic field, hypothalamus (PHT and THT), and notably, P3. This would extend the secondary prosencephalon up to the ZLI, reclassifying P3 as part of it and not of the diencephalon. This revised organization is supported not only by our TF code, but by our transcriptomic similarities -for example, in zebrafish, P3 cells are more similar to pallial (telencephalic) cells than to diencephalic types (**Fig. S07**). This new framework not only better describes developmental patterning, but also opens new perspectives on the evolution and specialization of the secondary prosencephalon.

In the mesencephalon and rhombencephalon, fine-grained clustering does not always align with the segmental boundaries described in classical histological studies^47,55^. For example, clusters in the rhombencephalon do not consistently follow HOX gene-based segmentation and integrate cells from different A-P or D-V classical segments (e.g. basal rhombomere 1–2; bR1–2). As others have noted^56^, such discrepancies may arise from assigning equal weight to functional effector genes and cell-type– specific TFs in clustering algorithms. However, they may also reflect the serial homology between these segments, where D-V identities (alar, basal) share a more evolutionarily conserved signature across several A-P neuromeres^57–59^.

Our single-cell atlases, in conjunction with existing morphological and histogenetic data, provide a robust foundation for defining the shared cell-type composition of the vertebrate brain bauplan (**Fig. 1h**). Yet, evolutionary inferences cannot rely solely on clustering annotations; they require rigorous, unbiased transcriptomic comparisons to uncover true cross-species relationships. To this end, we applied a comprehensive suite of complementary computational approaches **—** including GSI Correlations^60^, Label Transfer^61,62^, SAMap^63,64^, integration using Seurat v4’s “rpca” method^11,62,65^ **—** detailed in the following section.

### The transcriptomic conservation of brain bauplan cell types

Computational tools provide a more systematic and unbiased alternative to classical single-gene comparisons for analyzing -omic data. Yet, no gold-standard has emerged among these rapidly evolving methods^66,67^. Here, we leveraged the complementary strengths of multiple approaches to robustly assess the evolutionary conservation of the brain bauplan and of the single-cell transcriptomes across five vertebrate species. Our first goal was to identify homologous cell types independently of known marker genes (**Fig. 2a,b**). We applied two complementary mapping strategies: Label Transfer^62,68^ (**Fig. 2a**; **Fig. S08**) -which relies on 1-to-1 orthology-, and SAMap^63^ (**Fig. 2b**; **Fig. S09**) -which uses gene pairs based on BLAST similarities^69^. This dual validation strengthens our predictions and overcomes a key limitation in zebrafish, where teleost-specific genome duplication leads to a high proportion of 1:many orthologues^70,71^.

**Fig. 2.**
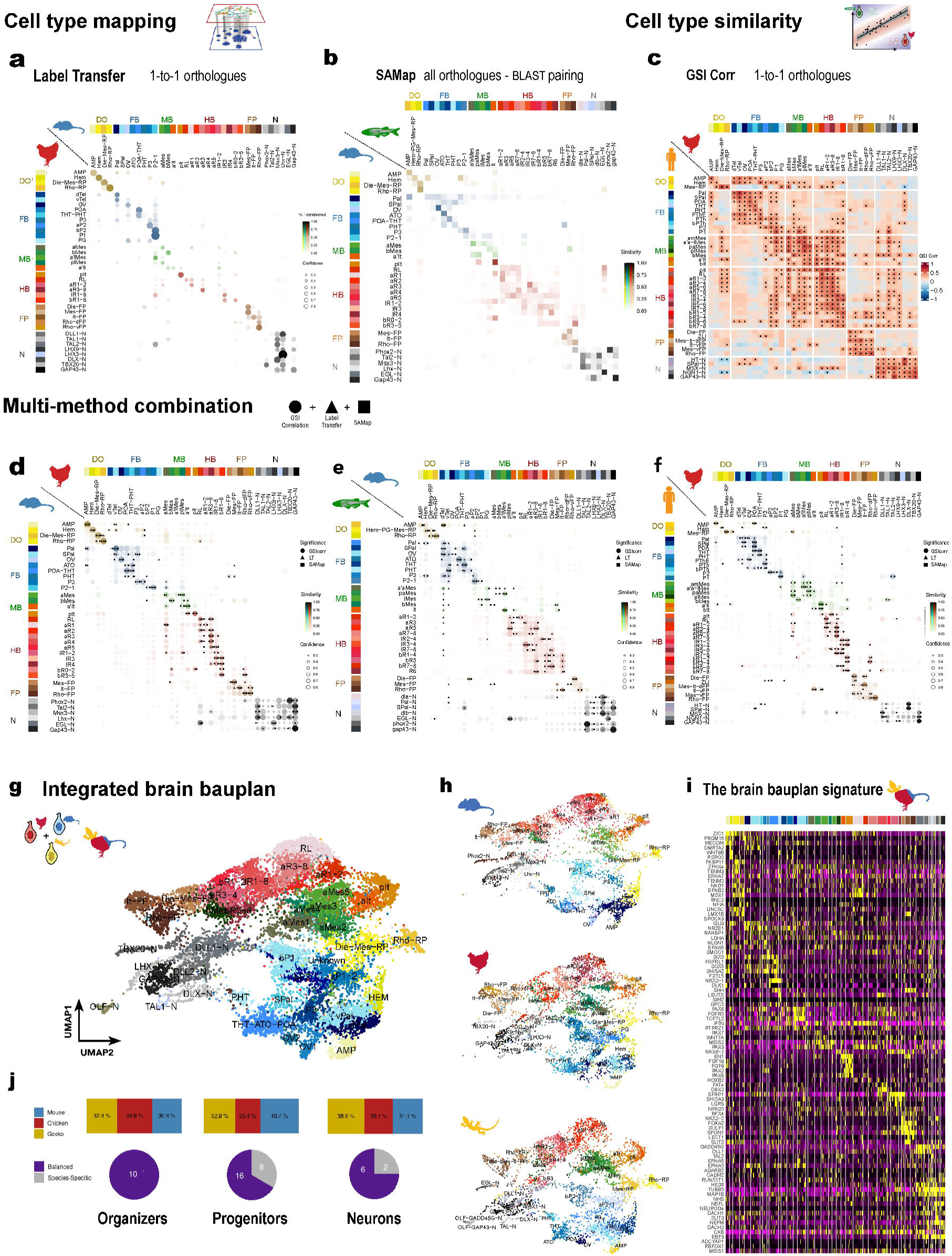
Inter-species computational cell type comparisons of the cellular brain bauplan. a-b, Cell type mapping: Label transfer (**a**, chick-mouse) and SAMap (**b**, mouse-zebrafish). **c**, Cell type similarity: GSI Correlations (chick-human). **d-f**, Multi-method combination (d, mouse-chick; e, zebrafish-mouse; f, human-chick). **g-j**, Amniote integration of early brains: **g**, Integrated cell types and UMAP plot. h, subset and original cell type retrieval. **i**, Brain bauplan signature of the integrated cell types by *spapros*. **j**, Cell species distribution per cell class and species specific cell types (>60%) within these classes. Color code in all plots matches with previous Fig. 1 code in each specific species, adapted for the integrated UMAP atlas. Circle size is associated to prediction confidence and color intensity to percentage transference (Label Transfer), similarity (SAMap) and correlation (GSI Corr).

In these comparisons, clusters in rows (query) and columns (reference) are ordered and color-coded to reflect the topological homologies hypothesized based on marker genes. In **Fig. 2a,b**; these brain bauplan homologies are transcriptomically validated by the unbiased assignment of most query cells to their hypothesized pair cell type in the other species. At first sight, the strong conservation of homologous cells is visible through the diagonal that emerges from dorsal organizers to neurons in both mapping methods. Nonetheless, when examining this overall conservation at the level of specific pairs, the preservation of brain bauplan identities becomes remarkable.

From the prosencephalic AMP and Hem to the roof plates of both diencephalon and mesencephalon, the dorsal organizer cells are most robustly mapped. Although these cells tend to combine into single clusters due to its similar and heterogeneous dorsal profile (e.g. zebrafish Hem-PG-Mes-RP), mapping approaches such as label transfer can distinguish these profiles with high confidence. Nonetheless, there are also exceptions to this paradigmatic cell type homology, as there are some cases where equivalent topological positions prevail (serial homology). For instance, the high similarities of the mouse Die-Mes-RP with the zebrafish Rho-RP, both roof plates from different A-P segments, but with a similar dorsal signature (Wnt signalling).

Among the most highly conserved cell types in the chicken-mouse label transfer, we also identify telencephalon, future pre-tectum and thalamus (P1-2), isthmus, rhombencephalic floor plate or mature neurons: depicting a high conservation of the whole brain bauplan. Although these “high conservations” could be slightly weeker in other pairwise comparisons (human-gecko), the existence of a systematic evolutionary conservation across all vertebrate pairwise comparisons rules out this likely single cell variability^46^. Overall, most neuromeres/cell types assessed confidently map to their homologs when compared across atlases, an indicator of their evolutionary conservation

Our second goal was to assess the transcriptomic conservation quantitatively, comparing the transcriptomic correlations across atlases (GSI Correlations^11,60,61^) (**Fig. 2c, Fig. S10**). As in the mapping plots, the highest correlation is always found between hypothesized homologous cell types. Correlation values reach up to ∼0.8 in many cases, including those cell types that previously exhibited more ambiguous mappings. These cell types can now be clearly paired into neuromeric divisions and subdivisions, such as the basal mesencephalon or alar rhombomeres 1 and 5.

In contrast to the mapping tools, correlations show a more complex transcriptomic conservation of single-cell transcriptomes, highlighting topological and cell class similarities. Firstly, cells within the same broad region show a higher correlation among themselves, but there are also similarities driven by D-V topological position rather than cell type. For instance, alar cells tend to resemble other alar cells regardless of the neuromere they belong to, and basal cells follow the same pattern. Conversely, cells within a given neuromere resemble each other even if they occupy different alar– basal domains. This likely serial homology is exemplified by the basal mesencephalon, which shows high correlation with basal prosomeres, especially the first two (bR1-2) in the human-chicken comparison. These global similarities can also be observed at the cell class level: progenitors and neurons display opposing transcriptomic signatures (blue, negative correlation, **Fig. 2c**). All neuroepithelial stem cells irrespective of their region share a positive correlation, an indicator of a common progenitor machinery that is evolutionarily conserved in all neuroepithelial types. The only exception are human comparisons, where progenitors transcriptomically resemble other species’ neurons, likely due to their more advanced developmental stage. Unfortunately, no earlier human data could be included in our comparisons.

Furthermore, these high correlations between interspecies homologs (e.g. AMP similarities) are even stronger when compared with other evolutionary cases in the literature. For instance, conserved adult GABAergic interneurons (0.5 maximum) or hippocampal derived cell types (0.25 maximum) across amniotes^60,61^, show much lower correlations than in the early embryonic brain (reaching up to 0.8, and more than 0.5 in most pair comparisons). These high correlations underline the strong preservation of the vertebrate brain bauplan.

To combine the strengths of these mapping and similarity methods, we created a summary plot based on previous publications^64^ (**Fig. 2c-e, Fig. S11**). This combined technique allows us to remove technical noise and highlight the shared trends within the three computational comparisons. For instance, those similarities that escape the predicted homologies and could be neglected as an artefact, such as bMes and TAL2-N; but the double significance (SAMap and GSI) prompted its further exploration to ultimately relate them as part of a common neurogenic cell lineage^72^.

The third objective of these computational comparisons was to assemble a cellular atlas of the vertebrate brain bauplan independently of clustering differences. For this purpose, all cells were merged regardless of origin at first. However, human and zebrafish were finally excluded due to their advanced maturation and the low feature number after 1-to-1 orthology filtering, respectively (**Fig. S12**). Furthermore, several integration/batch-removal methods were iteratively tested, but we primarily relied on the “rPCA”^62,68^ as the least coercive method for the integration of chick, mouse and gecko datasets (**Fig. 2e**). While other methods such as “Harmony”^73^ and “CCA” ^62,68^ merged the gecko-specific OLF-N cluster (possibly due to dissection differences) with other neuronal clusters, “rPCA” default parameters did not (**Fig. 2g, Fig. S13**). This biological robustness made us opt for this integration method to identify conserved and divergent cell types.

The integrated and newly reclustered atlas recapitulates the catalogue of cell types described in each species brain bauplan (**Fig. 1h**), even untangling dubious evolutionary scenarios. For instance, the cortical hem (Hem) is a dorsal organizer described in all individual atlases (**Fig. 1e-g**), but apparently absent in the gecko (**Fig. 1d**). However, we considered that gecko cortical hem cells had been clustered together with gecko dorsal telencephalon cells because of their shared antero-dorsal signature. Consistently with this explanation, our integrated atlas displayed gecko hem cells clustering with other species hem cells, resolving their cell type conservation over clustering/technical differences.

Overall, the conservation extent of the brain bauplan is illustrated by the integrated atlas, which shows the same cell distribution observed in individual atlases (**Fig. 1c-g**). Even when subsetting each species’ cells, cell organization reflected the original and remains undistorted when overlaid with other species’ cells (**Fig. 2h**). This became particularly evident for organizers such as the isthmus and the floor plates; however, it was less accurate when referring to progenitors, such as rhombencephalic neuromeres or prosomere 3. D-V position appears to be the main driving force in clustering, another indicator of the conservation across serial homologues and neurogenesis functional programs^55^.

These evolutionary trends are quantitatively reflected in the per-species cell proportions within the new clusters. All categories (progenitors, organizers, neurons) show a balanced proportion of cells from the three species globally, but regionalized progenitors show higher species-specific cell types (clusters in which one species shows >50% of total cells in the cluster) when looking at the new clusters (**Fig. 2f, Fig. S13d,e**). This variability might reflect the complexity of the early brain beyond the brain bauplan, where other *functional* gene networks related to growth or neurogenesis may play an important role.

Nonetheless, it seems that the gene network responsible for the induction of the vertebrate brain bauplan is highly conserved. We identify this blueprint by running *spapros*^74^ on this integrated dataset. The resultant brain bauplan signature of 100 master genes (**Fig. 2i**) perfectly distinguishes most cell types and is expressed across all compared species.

All in all, our interspecies single cell transcriptomic comparisons reveal a highly conserved brain bauplan signature across vertebrates, but they also hint at the existence of non-identity gene networks beneath the brain bauplan. To further explore the regulatory complexity of the early brain, new snATAC-seq atlases of our in-house species chick and mouse were added. In the next section, we will focus on these both species and their multi-omic atlases to untangle the GRNs evolution of the early brain.

### The genetic drivers of the early brain

Cell types or lineages are not merely associations of cells based on spurious gene patterns over single cell data; cell types are established by specific regulatory signatures^25^. Before the emergence of single cell technologies, early neuro-geneticists already defined the territories of the brain bauplan based on TF expression^75^. Nonetheless, such combinations of a few biasedly selected gene expression patterns did not utterly prove the existence and conservation of regionalized cell types.

On the other hand, -omic studies have unveiled the depth and modularity of GRNs^76,77^. Such molecular complexity may have diverged substantially underneath the brain bauplan conservation, but could it instead be even more conserved than its brain bauplan drivers? Up to date, published studies have approached the conservation of patterning and its molecular evolutionary causes (pleiotropy, lethality) from a rather bulk and whole-body perspective^21,22^. Here, we aim to characterize and compare the evolution of early brain GRNs focusing in mouse and chick.

TFs play an important role in cell type establishment, and are highly conserved at the cellular level in the early brain (**Fig. 3a**, left), more than their downstream targets (**Fig. 3a**, right). To identify the molecular drivers of the early brain and their brain bauplan identities, we extended the characterization of their cell types by complementary snATAC-seq atlases of 10773 in mouse and 28719 cells in chick (**Fig. 3b; Fig. S14–15**). Following label transfer from their scRNA-seq counterpart onto the RNA prediction matrix, cell types were confidently identified and both RNA/ATAC datasets were integrated by merging the same cell number per technology. These multiomic atlases allowed us to generate *pseudocells*^29^, which were essential to define the early brain GRNs (**Fig. 3b, bottom**).

**Fig. 3.**
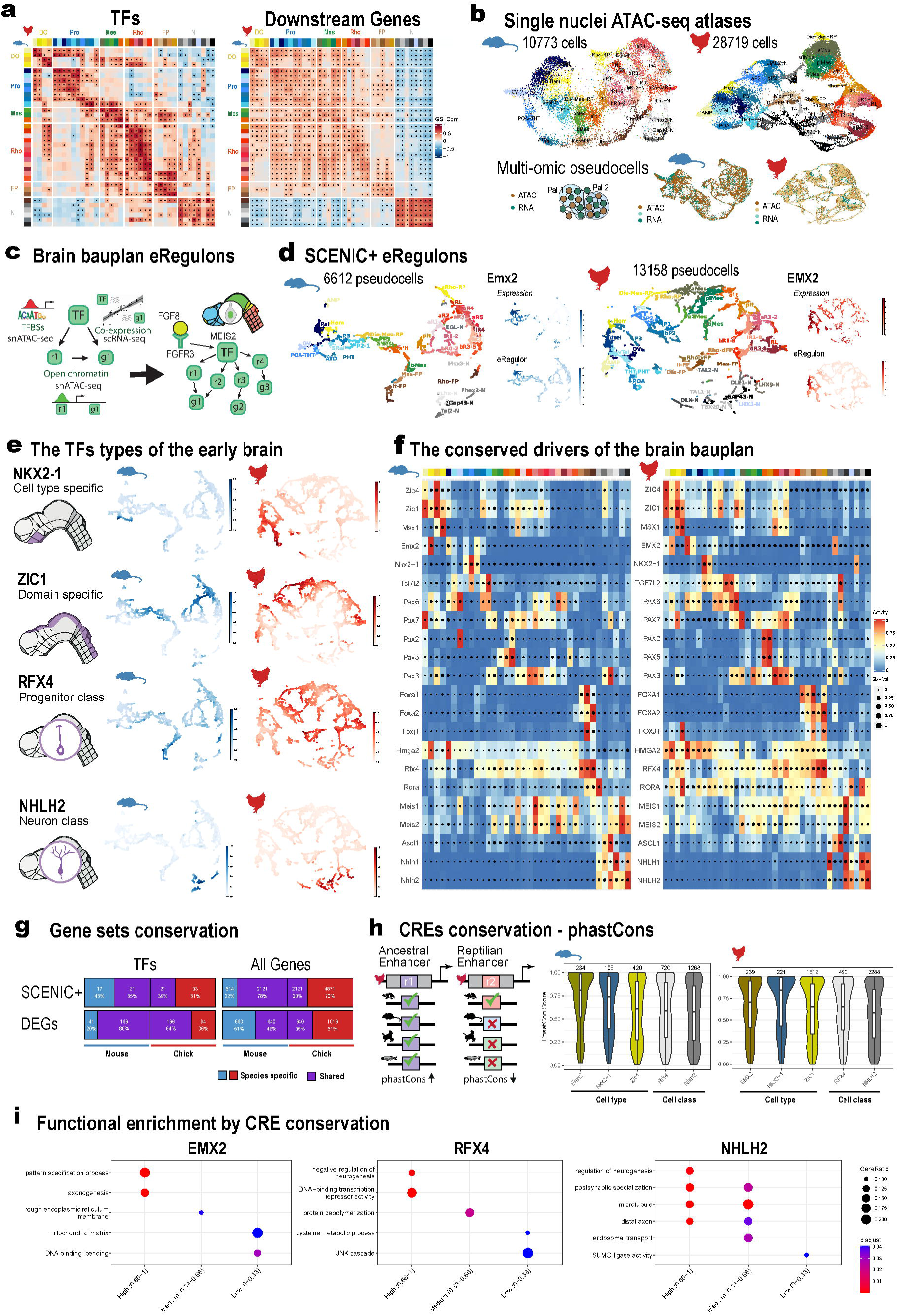
**Regulatory elements in the amniote early brain. a,b**, Chick against mouse GSI correlations filtered by only TFs (**a**) or non-TFs/morphogens (**b**). **b**, Single nuclei ATAC-seq atlases of mouse (10773 cells) and chick (28719 cells) (up). Cartoon representation of the pseudocells aggregation in the integrated RNA-ATAC atlases (bottom). **c**, SCENIC+ processing on multi-omic data to obtain GRNs and consequently TFs eRegulons. **d**, SCENIC+ eRegulons displayed as UMAP atlases of pseudocells. **e**, The TFs types of the early brain, representative activity on the neuroanatomical brain bauplan (left) and activity of its selected eRegulon in each amniote species (right). **f**, The conserved drivers of the brain bauplan. Simplified version of eRegulons for amniote shared TFs, irrespectively of detection method (region or gene). **g**, Gene sets conservation by cross-tabulation across SCENIC+ TFs and its downstream effector genes (first row); and all DEGs (FC > 0.25, pct = 0.25). Purple indicates shared percentage and number, while blue (mouse) and red (chick) are species specific. **h**, CREs conservation by phastCons scores of 100 bp windows across TFs eRegulons. Color code follows the cell type where TFs are mainly activated, and split between cell types and cell class. **i**, Functional enrichment by CRE conservation. Selection of most representative top10 functional terms (Y axis) for different phastCons bins (X axis) in EMX2, RFX4 and NHLH2 mouse eRegulons.

SCENIC+^78^ pipeline employed averaged multi-omic pseudocell data to model the relationships across TFs, genes and CREs into *eRegulons* –collections of genes and their associated regulatory regions that are all controlled by a single TF (**Fig. 3c**). Dimensional-reduction techniques allowed us to better visualize these eRegulons and their relative activity across *pseudocells* (**Fig. 3d,e**), and mean AUCell scores indicated their average importance per cell type (**Fig. 3f, Fig. S16–17**). In line with the high transcriptomic conservation of the brain bauplan cell-types, both species displayed highly similar patterns of eRegulon activity in both UMAPs and heatmaps (**Fig. 3d-f, Fig. S16-17**). However, the large number of pseudocells in the chick atlas yielded a broader and larger repertoire of eRegulons, which was reflected as a low conservation in cross-tabulation analysis versus mouse (**Fig. 3g**) ^46^.

In contrast to the conventional conception of cell-type-specific, these *cell-type* driver TFs are not confined only to a specific cell type (**Fig. 3e,f**). Some are rather region specific such as NKX2-1 – hypothalamus; and others show even broader within-brain pleiotropic expression (ZIC1, whole alar *domain*). These patterns suggest that unique cell type identities derive from the temporal and spatial combination of these within-brain pleiotropic TFs. For instance, EN1 controls the whole isthmus or midbrain-hindbrain boundary and FOXA2 controls all the floor plates (**Fig. 3e,f**); but FOXA2+/EN1+ defines exclusively the cell type isthmic floor plate. Furthermore, these *cell type* regional drivers are mostly expressed in both progenitors and their presumptive neuronal lineage, like TCF7L2^79^.

On the other hand, there are some TFs that lack deep regional specificity such as RFX4 and NHLH2, being expressed throughout most progenitors and neurons, respectively. For this reason, we have named this last TF type as *cell class TFs*, as drivers of progenitor or neuronal machinery; in contrast to *cell type TFs*, which specify regional domains. This regulatory complexity goes beyond the brain bauplan, highlighting the importance of cell class TFs for cellular function and developmental programs.

To further explore the GRNs below these conserved TFs, we explored the conservation of their downstream networks. Cross-tabular comparisons of effector genes could lead to false negatives, as substantial differences can be introduced by technical sequencing and annotation biases (**Fig. 3g**)^46^. Thus, we assessed the conservation of SCENIC+ network by a stable parameter: CREs’ phastCons scores^80^. This evolutionary parameter indicates the probability that a sequence has been syntenically preserved based on multiple species alignment. As already described by others^81,82^, the whole sequence of regulatory elements shows low conservation and cannot be lifted over genomes of such distant species (amniotes). However, here we show that 100 bp CRE windows do retain high sequence conservation in a manner similar to that observed in mammals (**Fig. 3h**)^29^.

However, the conservation of these CREs is highly heterogeneous (**Fig. 3h-i, Fig. S18**). Regarding eRegulon networks, the mean phastCons is higher for cell type TFs (e.g. EMX2, NKX2-1, ZIC1) than for cell class TFs (RFX4, NHLH2) (**Fig. 3h**). These differences would imply that cell type TFs-effector relationships are more constrained than more evolvable cell class networks. Furthermore, each eRegulon exhibits internal phastCons heterogeneity, which is linked to diverse functions. For instance, highly conserved CREs in EMX2 eRegulon genes are associated with *pattern specification* or *axonogenesis*, whereas the *mitochondrial matrix* shows weaker evolutionary preservation of EMX2 CREs (**Fig. 3i**). CREs conservation suggests a dependence on functional relevance, preserving brain bauplan patterning networks and loosening metabolism modules. These evolutionary constraints not only highlight the ancestral brain bauplan, but also the rapid evolution of TFs eRegulons and gene modules.

### The GRNs of the early brain

To further characterize the molecular complexity of the early brain, *hdWGCNA* was employed to perform gene co-expression network analysis of the mouse early brain (**Fig. 4a**) and later compare them with chick’s atlas (**Fig. 4f**). Based on their cellular expression patterns, hierarchical clustering of all genes led to early-brain representative gene modules. Among these modules, there are *class-specific* modules (**Fig. 4b**) and *cell type specific* modules (**Fig. 4c**), similarly to the previously presented shared genetic drivers.

**Fig. 4.**
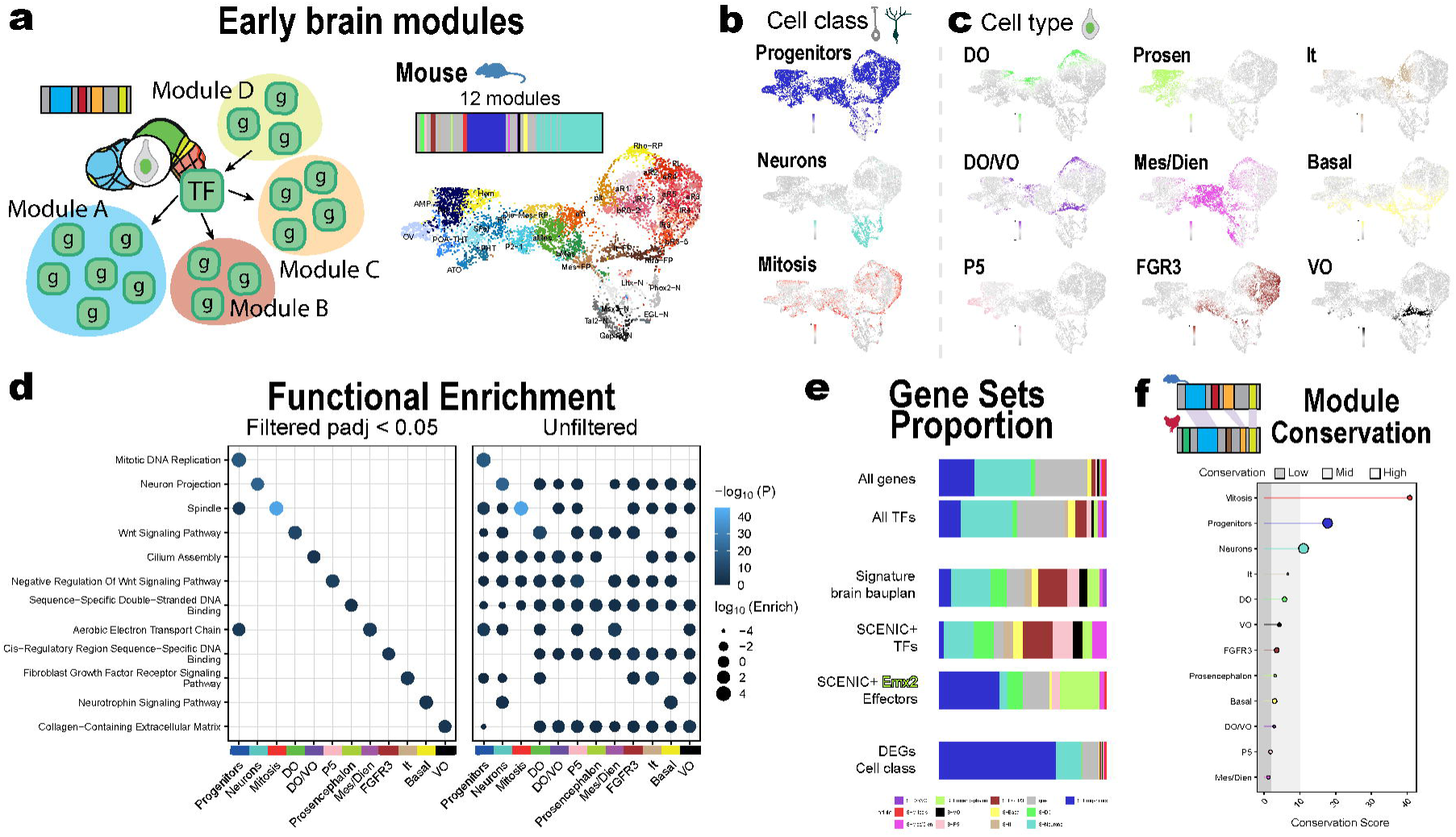
Gene modules of the early brain. a, Early brain modules of mouse atlas. Cartoon of a possible module detection within mouse early brain GRNs (left), hierarchical clustering of genes into modules (top right) and hdWGCNA single cell processed atlas (bottom right). **b-c**, Cell mean expression of gene modules across mouse single cell atlas split by cell class (**b**) and cell types (**c**). **d**, Functional enrichment of gene modules filtered by module-specific terms (left) and without (right). Color indicates significance and dot size enrichment. **e**, Proportion of modules within previously identified gene sets. **f**, Module conservation (Zsummary) between early brain mouse and chick atlases.

We termed all these modules according to their expression patterns, but also based on their associated functions (**Fig. 4d**). While class modules control broader functions (*proliferation* in *Progenitors*, *neuron projection* for *Neurons*), cell type specific modules are enriched for specific *morphogen pathways* (Wnt / FGF), but also cellular and molecular functions such as *collagen extracellular matrix* or *cilium assembly*. This enrichment filter searches functions mostly associated with only one module; but when this enrichment filter is removed, we can distinguish that most modules comprise gene sets belonging to all functional pathways. For instance, compared to others, *Prosencephalon* module is uniquely enriched for *Sequence-specific double-stranded DNA binding*, which are mainly brain bauplan genes. However, without module enrichment filter, *Prosencephalon* module also displays sets of *extracellular matrix, Wnt signaling and cilium assembly* genes, which appear in many other modules. Thus, all cell type modules are functionally similar regardless of their regional expression, but these gene signatures are distinctly unique for each specific region or cell type. Thus, prosencephalon cells will employ certain genes for their *extracellular matrix*, while basal plate cells will use others: leading to cell type differences.

This is further reflected in the composition of previous gene sets (**Fig. 3e**): On the one hand, brain bauplan (**Fig. 2i**) and SCENIC+ TFs (**Fig. 3d**) are mainly composed of cell type modules; and on the other, the neurons vs progenitors DEGs are enriched for class modules. Meanwhile, in the Emx2 eRegulon – an important prosencephalic TF (**Fig. 3f**)-most effector genes are part of the general *Progenitor* and *Prosencephalon* modules, an example of how progenitor functions depend on both broad cell class and cell type specific genes.

To investigate the evolutionary conservation of these gene modules, we transferred mouse modules into the chick atlas and assessed their preservation. The overall preservation (*Zsummary*) is higher for class modules (**Fig. 5f**), but it also biased by module size (cell class modules are larger). In contrast, module-size-independent metrics such as *medianRange* point out the also high conservation of cell type modules like *Isthmus* module (*It*, related to patterning). Nonetheless, cell class modules such as *Progenitor* and *Mitosis* also maintained their high conservation, but *Neuronal* module showed much lower medianRange preservation. Cell type modules such as *Prosencephalon* increased their preservation values in size independent metrics, but their conservation was substantially lower than stemness modules like *Progenitor* and *Mitosis* (**Fig. S19)**. This lower preservation could be explained by previously described heterogeneity (**Fig. 4d**): cell-type–specific modules are composed of conserved regulatory genes, but also enriched in functional cellular programs, that freely evolve (phastCons CREs, **Fig. 3i**): a spark of divergence (changes in size or timing) within conserved cell-type-lineages^10,28,83–87^.

**Fig. 5.**
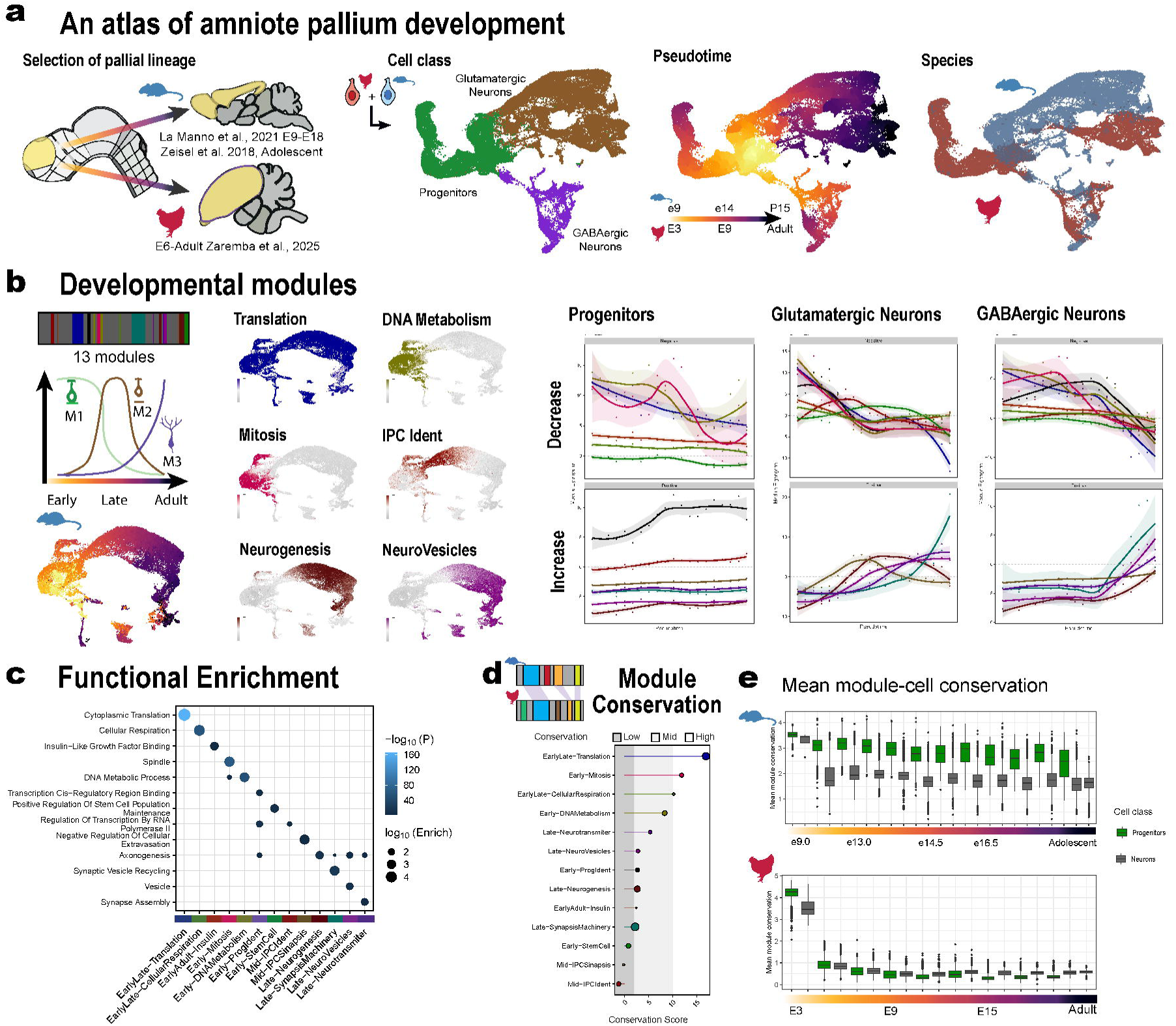
Gene modules of the amniote pallium development. a, Atlas of amniote pallium development. Schematic representation of developmental trajectories in mouse and chicken, and the associated employed datasets^40,64,88^ (left). Single cell integrated atlas of both amniote species coloured by cell class (green, neural progenitors; brown, glutamatergic lineage; purple, gabaergic lineage), pseudotime (monocle3^128^ translation of cell developmental trajectories) and species (blue, mouse; red, chick). b, hierarchical clustering of genes into modules (top left), hdWGCNA single cell processed atlas (bottom left), cell mean expression of gene modules across pallium development and tendencies on cell lineages (progenitors, glutamatergic and GABAergic lineages). **c**, Functional enrichment of gene modules filtered by module-specific terms. **d**, Module conservation (Zsummary) between developmental pallium mouse and chick atlases. **e**, Mean module-cell conservation in the pallium developmental cell classes of mouse and chick atlases.

Overall, the conservation (Z-summary and medianRange) of stemness modules (*Progenitor* and *Mitosis*) highlights them as an underexplored additional constraint in the early brain, which could have the same impact as the brain bauplan identity program. Yet, it remains unresolved whether early brain modules actively contribute to a phylotypic constraint, or whether cell-class modules merely represent a developmental constant. A global developmental perspective is thus essential to properly contextualize these findings.

### The GRNs of pallium development

Organogenesis has always been conceived as the most conserved stage of development, but previous longitudinal studies disregarded the molecular complexity of cell types and underlying gene networks^20–22,27^. To shed light on the existence of an hourglass within brain evolution, we further focused on the sequence of pallium development as a model for the brain. We extended the transcriptomic analysis to other developmental timepoints till adulthood/adolescence in mouse and chick pallium from available databases [La Manno et al. (2021)^40^, Zeisel et al. (2019)^88^, Zaremba et al. (2025)^64^ (**Fig. 5g, Fig. S20**)].

When integrating both species developmental trajectories, early stages cells — especially progenitors — are intermingled regardless of their original species. However, cells from later stages follow separate species-specific developmental trajectories to mature progenitors and neurons (**Fig. 5g, Fig. S20**).

To identify developmental modules and their evolutionary preservation, we first profile the developmental atlas of mouse brain with hdWGCNA. Hierarchical gene clustering leads to 13 modules that we named after their enriched functions (**Fig. 5i**). These modules show different developmental expression patterns (**Fig. 5h, Fig. S20**): some display a positive activity over developing time (**Fig. 5h** – top row), whereas others present a negative trend (**Fig. 5h** – bottom row). For instance, *Translation module* (dark blue lines) is expressed throughout development, but higher at early stages. In contrast, other modules are exclusive of specific stages or cell types such as *DNA Metabolisms* (early-stage progenitors), *IPC ident* (intermediate progenitors), *Neurogenesis* (late development, but no adult) and *Neurovesicles* (neurons of all stages). To assess if these different functions shape a different evolutionary conservation over development, we compared these mouse modules by transfer to the chick counterpart.

Module preservation assessment to the equivalent chick pallial modules showed different evolutionary scores, which clearly depended on their main developmental expression patterns. Early modules are overall more conserved (Zsummary) than intermediate (*IPCs*) or later (*Neurogenesis* / *Synapsis Machinery*) modules (**Fig. 5j** left). The higher conservation of early stages is better visualized when plotting the mean module Zsummary per cell and across developmental time (**Fig. 5j** – right panels). The highest conservation is reached at the phylotypic stages, and decreases proportionally to time. This decay is more evident for neurons, possibly as they lack highly conserved proliferation modules (*Mitosis* and *DNA Metabolism*). Therefore, the early pallium is not only highly evolutionarily conserved due to its brain bauplan drivers (**Fig. 4**), but also due to its underlying progenitor molecular machinery (cell class modules).

### The evolutionary constraints of brain development

The evolutionary mechanism behind the conservation of these early neural GRNs remains unresolved (**Fig. 6a**). Whereas the *pleiotropic hypothesis* claims that the phylotypic stage is essential due to pleiotropic genes expressed across multiple organs and stages^21,22,89^, other hypotheses claim that mid-embryogenesis essentiality derives from the function and impact of early genes, a domino-like effect (*developmental burden*)^90^. To further investigate these hypotheses, we calculated gene-level parameters on developmental single-cell transcriptomes from whole-brain and pallial atlases ^40,64,88^ . These parameters were *gene lethality* — viability upon gene loss *in vitro* (cell) or *in vivo* (embryo, developmental) — from IMPC classification^91^ (only mammals) and *pleiotropy* —breadth of gene expression across tissues and time points— from Cardoso-Moreira et al. (2019)^21^, both mouse and chick.

**Fig. 6.**
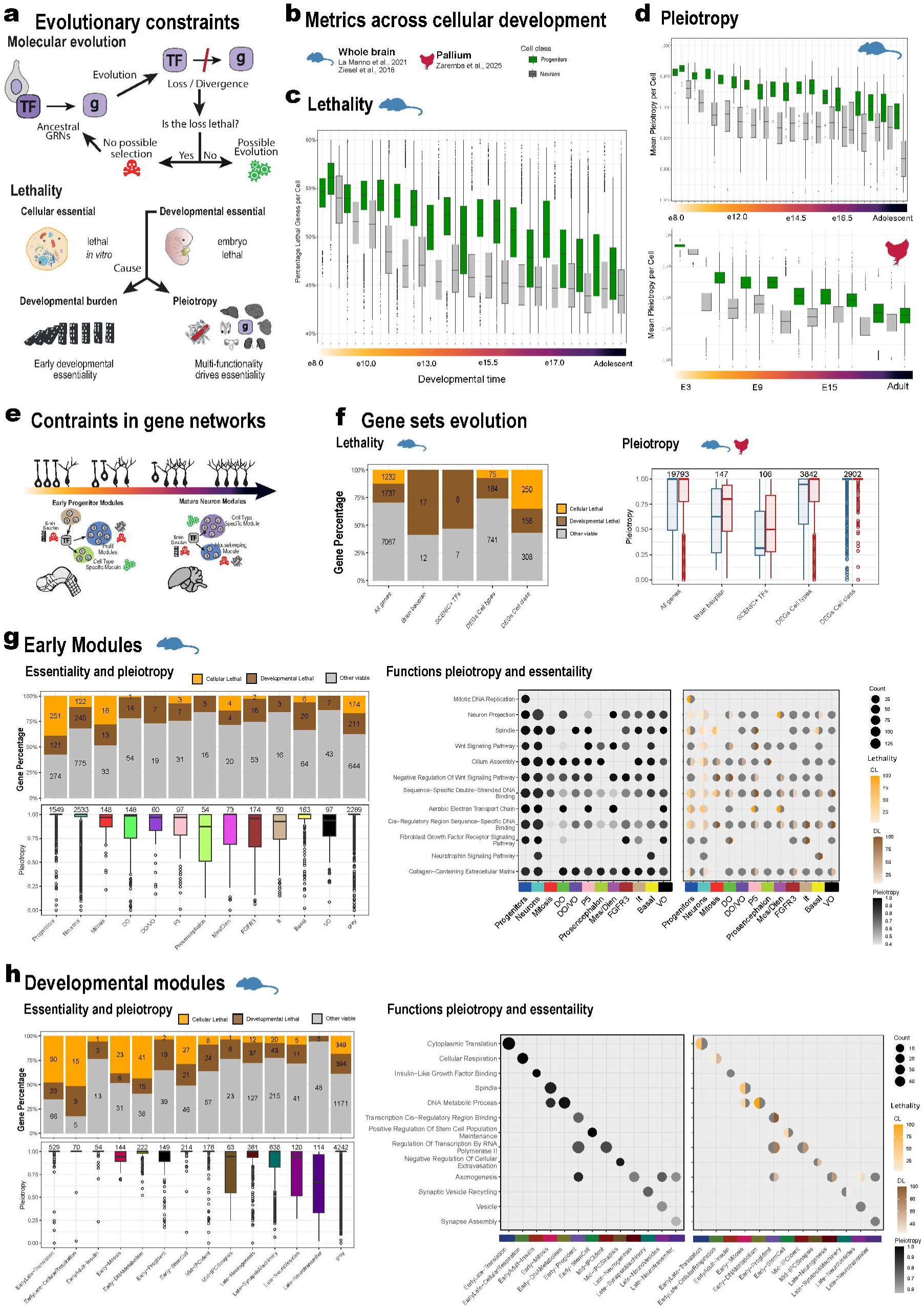
**Molecular mechanisms behind the phylotypic evolutionary constraint**. **a**, Cartoon of TF-gene relationship evolution based on essentiality. **b**, Metrics across cellular development: atlases and cell classes (green, progenitors; grey, neurons). **c**, Lethality per cell class and stages (mouse). **d**, Pleiotropy per cell class and stages (mouse, top; chick, bottom). **e**, Model for the evolutionary constraints in developmental gene networks (previously used color code). **f**, Gene parameters for lethality and pleiotropy. **g**, Essentiality and pleiotropy of early modules. Essentiality (left up) and pleiotropy (left bottom) of module set of genes. Functional pleiotropy (central) and essentiality (right) of the module specific functional gene sets. **h**, Lethality (left up) and pleiotropy (left bottom) of whole development pallial modules. Essentiality and pleiotropy of module set of genes. Functional pleiotropy (central) and essentiality (right) of the module specific functional gene sets.

Early brain developmental stages display higher percentages of essential genes, which decrease in later stages and is lower for neurons (**Fig. 6b,c**). Pleiotropy values follow the same trends in both amniote species: higher in early stages and lower for the neuronal class (**Fig. 6d**). Moreover, these plots correlate strongly with the mean module conservation in **Fig. 5j**, further suggesting that pleiotropic and essential genes of early timepoints drive the overall transcriptional conservation of the phylotypic brain. However, these mean values do not capture the complexity of GRNs and possible mechanistic differences across regulatory networks or functions, nor the relative contributions of different parts of the GRNs.

To further understand the different possible evolutionary constraints, we explored lethality and pleiotropy over the gene sets and modules previously described (**Fig. 2, 3, 5**). Regarding gene sets (**Fig. 6f**), the brain bauplan genes (*spapros*, **Fig. 2i**) are composed of highly essential developmental genes (>50%). However, these genes show very low pleiotropic scores compared to all genes’ distribution, which are even lower in the case of SCENIC*+* drivers. When we expand the list of cell type marker genes to *Seurat* DEGs, up to 200; there is an increase of cell type specific nonessential genes that might be related with less evolutionary constrained cell-type-modules, likely favoring inter-species differences (**Fig. 6f**). On the other hand, cell class DEGs are similarly enriched for essential genes; but their pleiotropy scores reach higher scores – 1 in > 75% of genes. This high pleiotropy indicates a broad expression across all organs and stages, a likely cause of evolutionary essentiality^22,24^.

This is further illustrated by specific early brain modules (**Fig. 4**), which associate these high levels of pleiotropy with stemness modules (**Fig. 6g**). For instance, *Progenitors* module displays a high percentage of essential cellular genes and the highest pleiotropic values, which are associated to functions like *mitotic DNA replication*. In contrast, other modules such as *Prosencephalon* show fewer developmental essential genes and low pleiotropic values, which are also functionally explained: *TFs binding* (essential) or *collagen extracellular matrix* (few essential genes). The existence of non-essential genes within cell-type-specific signatures makes them less evolutionarily constrained, likely easing its divergence compared to cell identity or stemness signatures.

Nonetheless, the early brain is characterized by its essential functions: proliferation and cell identity (**Fig. 5**). This evolutionary constraint is highlighted when contrasted with later development or adult stages (**Fig. 6h**). In our developmental modules, early signatures show higher essential and pleiotropic modules (*Translation* and *DNA Metabolism*), and associated essential functions such as *cellular respiration* or *spindle*. By contrast, later stages modules contain non-essential or tissue-specific genes, which are more likely to evolve and ultimately explain the divergence of the late and adult brain.

As a whole, the vertebrate brain bauplan is defined by a transcriptomic signature that is rather brain specific. Although some brain bauplan genes show a certain degree of bricolage^92^, its expression breadth is more limited than stemness machinery, which is expressed across all proliferative cells.

Thus, while the evolutionary conservation of the brain bauplan signature might rather derive from its early developmental impact in cell identity (developmental burden hypothesis^90^), stemness machinery is more likely to have been preserved for its essentiality in all proliferating cells (pleiotropy hypothesis^89^). Altogether, our findings suggest that both the elevated essentiality of the brain bauplan genes and the high pleiotropy of stemness genes act as phylotypic constraints.

## Discussion

Here we single-cell profiled five vertebrate early brains, covering from fish to mammals and reptiles. Through unbiased clustering and a neuroanatomical cell type–gene dictionary, we identify a conserved vertebrate cellular architecture underlying the brain bauplan, further validated by complementary computational transcriptomic comparisons. In addition, our GRNs analysis dissects the regulatory complexity of the early brain, as well as its functional modules across cells and in the whole developmental context. This GRN analysis reveals the additional conservation of stemness genes, especially relevant in early developmental stages.

Our single cell atlases, gene catalog and consensus cellular model for the brain bauplan not only reinforce the essential role of developmental patterning in evolutionary neuroscience, but also shed light on long-standing debates regarding the boundaries of evolutionary neural identities^35,37,50^. Our cellular neuromeric model supports the prosomeric model^6^, validating its hypothetical cell type boundaries based on sparse histological data. However, we also contradict some of its broader evolutionary relationships, considering P3 or prethalamus as developmentally and evolutionarily related to secondary prosencephalon. Following the prosomeric model^6^, our model provides evidence that confronts classical columnar theory^49^ and other current alternatives^50^, suggesting a shared evolutionary origin of the posterior hypothalamus and the telencephalon^51–53^. Our multi-omic identification of Tel-PHT common regulatory elements (e.g. EMX2) argues against a hypothalamus-diencephalon relationship, redirecting research toward the emergence of the hypothalamus-telencephalon^49,50,53,93,94^. Was the hypothalamus ancestral to the telencephalon?^51^ Was there an ancestral anterior neural center distinct from both the hypothalamus and telencephalon?^53^

Homologies should be established based on highly evolutionarily constrained elements^31^; and our combination of computational transcriptomic comparisons positions early-brain cell type transcriptomes as such. First, by mapping the highly conserved cell identities across our atlases; and then, by evaluating the high transcriptomic similarities across cell types. However, these comparisons reveal similarities between homologous or topologically equivalent identities within the neural tube, underlining a common domain molecular signature (alar, basal, mesencephalon). These broad transcriptomic similarities also extend to cell classes, showing high similarities between progenitors and neurons. This high evolutionary conservation was also noted in our multi-species integration, but these vertebrate integrated cell types already displayed evolutionary differences, with organizers being more conserved than progenitors. This could be explained by an early diversification of neuroepithelial cells, as we have further seen in their cell-type-specific signatures. However, the overall brain bauplan signature from all integrated cell types clearly reveals the vertebrate cellular model, originally depicted in the individual atlases.

The comparison of gene modules beyond marker genes is essential for understanding cellular function in evolutionary studies^26^. Our GRN analysis untangles previous bulk and aggregated longitudinal studies^21,22^, revealing how early brain stemness is a true evolutionary constraint across its whole development. Moreover, we also distinguish the mechanism behind this conservation, disentangling the high pleiotropy of early stages. These high pleiotropy values did not arise from patterning genes^89^ -which do show a certain degree of bricolage^92^, especially within the brain-but from the broad expression of stemness modules. An extremely conserved molecular machinery present in likely all proliferating cells, probably even more ancient and tightly preserved than the brain bauplan gene machinery^95^.

Given this complexity, the early brain is not a mere example of conservation, but a proxy to understand evolutionary constraints. Cell identities established during early patterning are maintained along their subsequent neurogenic lineages by patterning TFs^75,96^, but early downstream modules associated with stemness in progenitors are overtaken by brain-specific modules later in development. While stemness modules are extremely pleiotropic and evolutionarily conserved, later intermediate progenitor and neuron modules are more tissue-specific and consequently less evolutionarily constrained^24,97^. This developmental trend might be a general principle across organs and cell lineages, representing a paradigmatic rule in evolutionary developmental biology.

This evolutionary principle can help us to better define “cell types”, a subject still under debate in the era of single-cell genomics^25,98,99^. TFs are hypothesized to play a central role in cell type identity, but their action is far from being cell type specific. In the early brain, most TFs show a broad regional patterning action (e.g. ZIC1 on alar domains), and others control cell class functions such as progenitor stemness (e.g. RFX4) or neuronal maturation (e.g. NHLH2). It is only the combination of these “*within-brain-pleiotropic*” TFs that leads to specific cell types and lineages (e.g. MEIS2 with NKX6-results in bMes; and MEIS2 with ZIC1 becomes aMes). However, should cell class TFs such as RFX4 and NHLH2 have a role in defining cell types? Or do they just reflect a maturation state in the cell-type-specific maturation lineage or trajectory? This conflict is especially relevant in more advanced stages of patterning such as the pallium^100,101^, where cell identities are finer and our distinction between cell class and cell types TFs get blurrier (e.g. RORA)^60,64,65^. Despite this need for deeper characterization of regulatory elements and their combinatorial interaction for cell identity, here we demonstrate that evolutionary unconstrained features such as neuronal connectivity or cherry-picked expression patterns tend to be less evolutionarily informative^31,32^. In contrast, highly evolutionary constrained regulatory elements such as patterning TFs can serve as the Rosetta Stone for homology^75,102^.

On the other hand, even within the conserved phylotypic brain, there exists evolutionary heterogeneity ranging from highly conserved processes (*pattern specification*) to other more divergent functions (*mitochondrial machinery*). Only by untangling the complexity of GRNs can the different conservation degrees of gene modules be clearly identified regardless of developmental stages. This is especially relevant in adult processes such as hematopoietic fate determination, where cell identity program is likely to be as conserved as early brain bauplan^25^. In contrast, early mitochondrial networks are likely to have diverged as much as those in the adult brain^103^. After all, the perfect substrate for evolution is modular non-essential genes, no matter when these are expressed^24,97^. Later stages are enriched in these cell-type-specific genes, but the historically conceived conserved phylotypic stage might also encompass more evolvable features. Underneath the bauplan, the tinkering of its downstream unconstrained genes could be the spark for divergent proliferation (allometry) or/and maturation timing (heterochrony)^10,28,83–87^.

In conclusion, our cellular comparisons not only revealed an ancestral brain bauplan at transcriptomic and regulatory levels, but also unveiled a deeply conserved set of downstream pleiotropic modules related to stemness. A dual evolutionary constraint (cell identity and stemness) that underlies the conservation of early brains in vertebrates.

## Abbreviations

a: alar (plate)
a′: anterior
ACC: accessible regions matrix
AMP: anterior medial pole
ATAC: assay for transposase-accessible chromatin
ATO: acroterminal organizer
AUC: area under the curve
b: basal (plate)
BAM: Binary Alignment/Map
BED: Browser Extensible Data
CCA: canonical correlation analysis
CDS: coding sequence
CL: cellular essential
Corr: correlation
CRE: cis-regulatory element
CS: Carnegie stages
DAR: differentially accessible region
DEG: differentially expressed genes
Die: diencephalon
DL: developmental essential
E: embryonic day
EI: early immature
FB: forebrain
FBS: fetal bovine serum
FP: floor plate
GEM: gel bead-in-emulsion
GEX: gene expression matrix
GRN: gene regulatory network
GSEA: gene set enrichment analysis
GSI: gene specificity index
HB: hindbrain
HH: Hamburger-Hamilton stages
hpf: hours post-fertilization
HT: hypothalamus
IMPC: International Mouse Phenotyping Consortium
It: isthmus
l: liminal
LI: late immature
MB: midbrain
Mes: mesencephalon
N: neuron
NE: neuroepithelium
NGS: next-generation sequencing
NS: nucleosome signal
NSC: neural stem cell
OV: optic vesicle
P: prosomere
PCA: principal component analysis
PHT: peduncular hypothalamus
PhT: prethalamus
PhTE: prethalamic eminence
POA: preoptic area
Pro: prosencephalon
QC: quality control
R: rhombomere
R2G: region-to-gene relationship
RL: rhombic lip
Rho: rhombencephalon
RP: roof plate
RPCA: reciprocal principal component analysis
RSS: regulon specificity score
RT: reverse transcription
scRNA-seq: single-cell RNA-seq
Seq: high-throughput sequencing
SNN: shared nearest-neighbor
snATAC-seq: single-nucleus ATAC-seq
SPal: subpallium
TAI: transcriptome age index
Tel: telencephalon
TF: transcription factor
TF2G: TF-to-gene relationship
TF2R: TF-to-region relationship
TFBS: transcription factor binding site
THT: terminalis hypothalamus
TSS: transcription start site
UMAP: Uniform Manifold Approximation and Projection
ZLI: zona limitans intrathalamica
ρ: Spearman’s rank correlation coefficient.

## Methods

### Animals

Data shown in this study derives from animal experimentation conducted by authors and available datasets from published studies (see Data availability section), as shown in Table S1.

In-house animal experiments were approved by a local ethical review committee and conducted in accordance with personal and project licenses in compliance with the current normative standards of the European Union (Directive 2010/63/EU) and the Spanish Government (Royal Decrees 1201/2005 and 53/2013, Law 32/107).

**Chick:** Fertilized chick eggs (*Gallus gallus*) were purchased from Granja Santa Isabel and incubated at 37.5 °C in humidified atmosphere until required developmental stage. The day when eggs were incubated was considered embryonic day (E)0.

**Mouse:** Adult C57BL/6 mice (*Mus musculus*) were obtained from a mice breeding colony at Achucarro Basque Center for Neuroscience (Spain). They were housed in a 12/12-hour light/dark cycle (8 AM, lights on) and provided with *ad libitum* food and water. T The day when the vaginal plug was detected was referred to as embryonic day 0.5 (E0.5) -to accommodate to La Manno et al (2021) dating.

**Gecko:** Fertilized ground Madagascar gecko eggs (*Paroedura picta*) were obtained from a breeding colony at Achucarro Basque Center for Neuroscience. They were housed in a 12/12-hour light/dark cycle (8 AM, lights on, 27°C/ 8 PM light off, 22°C) and provided with *ad libitum* food (live crickets) and water. Gecko eggs were incubated at 28 °C in a low humidified atmosphere until required developmental stage. The day when eggs were incubated was considered E0.

Selection of comparative stages across vertebrate species (human, mouse, chick, gecko and zebrafish).

To select a specific timepoint that captured the brain bauplan, a representative stage of this early brain was selected based on two main criteria:

**1.** Neurulation must have been completed at the brain end (anterior neural pore closure). Being every broad segment of the adult brain supposed to be already patterned.
**2.** Existence of few neurons. Neurogenesis is itself an indicator of the early bauplan establishment as progenitors are patterned before dividing into neurons. Avoiding neurogenic stages allows us to better study early patterning and segmentation in neural stem cells. Later developmental timepoints will rather depict more progenitor’s sub-regionalization (e.g. pallium subdivisions) and neuron maturation.

The selected stages are the following: mouse (E9.5), chick (E3.2, equivalent to stage HH19), gecko (E5), human (CS13) and zebrafish (16/18/20 hours post-fertilization (hpf)) ^20,22,104^. The selected human timepoint is more advanced than the other species, not complying to the neuron abundance criterion. Ideally, the first stage after neural tube closure would be CS12 (4 weeks after conception), but there are no available datasets of such an early stage –Xu et al. (2023)^105^ lacks enough quality cells to conduct a comparable atlas. In the zebrafish, several stages have been grouped to expand cell profiling, as all fulfill the criteria and individual stages did not have comparable cell numbers to other species.

On the other hand, developmental program comparisons were only possible regarding the pallium – annotated as dorsal telencephalon by La Manno et al. (2021) - as there was no more available chick data besides Zaremba et al. (2025)^64^. Thus, mouse whole brain developmental data from La Manno et al. (2021)^40^ was subsetted for dorsal telencephalon. Since dissections prior to E12 did not distinguish between dorsal and ventral telencephalon, only samples from E13 onwards were retained. A comparable gap in early developmental stages was also present in the chick datasets, which lacked samples between E3 (this manuscript) and E6 (Zaremba et al., 2025).

### Brain dissections

There was a common dissection procedure for brain isolation that had subtle variations for every species. For mouse embryos, the pregnant mother was euthanized, the uterine horns were exposed by laparotomy and embryos were extracted out of the yolk sac. For chick and gecko embryos, embryos were extracted from egg yolk and vascularized amnion with tweezers. Brains were placed in a Petri dish with ice-cold HBSS to maintain cellular viability. All embryos were micro-dissected under a stereomicroscope to remove mesenchymal and endodermal tissue (including optic vesicle) surrounding the brain neuroepithelium. Thus, the brains here characterized were established from the anterior-most forebrain region to the otic vesicle level at the hindbrain.

Single cell RNA sequencing (scRNAseq) Cell preparation:

Every dissected brain was visually segmented into prosencephalon, mesencephalon and rhombencephalon; by morphological dissection of the three neural tube enlargements or vesicles down to the otic vesicle. Then, three pieces were placed into a petri dish where they were enzymatically disaggregated in a species-specific manner, as previously described^106^. Chick brains undergone an enzymatic disaggregation of 37°C and 15 minutes; for the gecko, brains enzymatic disaggregation was conducted at 27°C for 20 minutes to convey their body temperature and maintain equal enzymatic activity.

After the incubation, tissue was manually homogenized in the Petri dish by carefully pipetting with descending-volume pipettes (P1000, P200, P2). Tissue clogs were removed by filtering the cell suspension through a 40 µm nylon strainer to a 15 ml Falcon or conical tube containing inactivating enzyme solution (4 ml of 25% fetal bovine serum (FBS) in HBSS). To dilute biological variability, each Falcon tube included the filtered cell suspension from two to three pooled brain embryos, at least.

Dissociated cells were then centrifuged at 200g for 10 min at room temperature. The solution was diluted with HBSS until obtaining a concentration of 1200 cells per microliter, as recommended by the single cell capture equipment manufacturer (*10X Genomics*). To reach that concentration, it was serially diluted, and concentration was calculated using a TC20 Automated Cell Counter (*BioRad*). To do TC20 measurements of viability and number, cell suspension was mixed 1:1 with trypan blue (*BioRad*).

Up to 24,000 cells per reaction were loaded into the Chromium 10X (*10X Genomics*) (Droplet-based method) aiming a targeted cell recovery of 12,000 cells (with estimated 50% efficiency). As described before, single cell library was prepared using “Chromium Next GEM Chip G Single Cell Kit”, 16 rxns PN-1000127, “Chromium Next GEM Single Cell 3ʹ Kit v3.1*”*, 4 rxns PN-

1000269 and *“*Dual Index Kit TT Set A*”*, 96 rxns PN-1000215, following *“*Chromium Next GEM Single Cell 3*’* Reagent Kits v3.1(Dual Index) User Guide*”* (Document number CG000315). Following 10X Genomics library kit, the PCR product was purified and quantified. The obtained libraries were sequenced on a Novaseq 6000 (*Illumina*) for an approximated 50.000 reads per cell.

### Gecko *-Paroedura picta –* genome

Alignment against the current chromosome-level genome of Madagascar ground gecko (*Paroedura picta*)^107^ yielded extremely low read-mapping percentages. To improve this yield, a new annotation was generated by the *GeMoMa*^108^ pipeline. Apart from *in silico* predictions, this pipeline combined all the transcriptomic evidence existent in public repositories (*NCBI BioProject PRJDB4004*): *DRR047248*, *DRR047249*, *DRR047250*, *DRR047251*, *DRR047252*, *DRR047253* and *DRR047254*; and also the proteome of closed related species that had high quality annotated genomes in *NCBI*

Genomes:

- *Zootoca vivipara* (Common lizard) (UG_Zviv)
- *Lacerta agilis* (Sand lizard) (rLacAgi)
- *Sceloporus undulatus* (Fence lizard) (SceUnd)
- *Sphaerodactylus townsendi* (Townsend’s least gecko) (MPM_Stown)
- *Pogona vitticeps* (Central bearded dragon) (PogVit)
- *Pseudonaja textilis* (Eastern brown snake) (EBS)
- *Hemicordylus capensis* (Graceful crag lizard)
- *Podarcis raffonei* (Aeolian wall lizard) (rPodRaf)
- *Python molurus* (Indian python) (PytMol)
- *Protobothrops mucrosquamatus* (Brown-spotted pit viper) (ProtoMuc)
- *Thamnophis sirtalis* (Common garter snake) (ThamSirt)
- *Pantherophis guttatus* (Corn snake) (PanGut)
- *Gekko japonicus* (Schlegel’s Japanese gecko) (JapGek)
- *Varanus komodoensis* (Komodo dragon) (ASM)
- *Thamnophis elegans* (Western terrestrial garter snake) (ThaEle)

The annotation output was highly satisfactory, as our annotation reaches equivalent levels of read alignment to the transcriptome to the current chick genome (please see *Table S2*).

The details of the pipeline employed are available on the *GitHub* site: https://github.com/rodrisenovilla/Senovilla-Ganzo2025.

### Single cell transcriptomics pre-processing

For both species, 10X *CellRanger* v7.1.0 was employed for alignment and demultiplexing of FASTQ files to obtain feature-barcode matrices. The genomes used as reference for each species alignment were: mm10-2020-A for mouse, galGal6 (Ensembl 99) and also bGalGal1 (Ensembl 112) for ATAC/SCENIC+ pipeline, and our custom annotation for the gecko genome. The quality control statistics related to this alignment steps and others are summarized in Supplementary Table S3. Afterwards, data matrices were imported to *R* (v4.1.0), where *Seurat* (v4.1.0)^109^ was employed to further analysis, as describe in their vignettes (https://satijalab.org/seurat/).

The output matrix of both in-house samples and published datasets were independently input to *Seurat* and processed following a common pipeline (*Figure 35*). The cell-cycle phase was determined by CellCycleScore() function, using “RRM2”, “PCNA”, “SLBP”, “WDR76”, “MCM5” as S-phase genes and “CENPF”, “TPX2”, “HMGB2”, “UBE2C”, “BUB1B”, “TOP2A”, “CENPE”, “TACC3”, “BUB1”, “AURKA”, “CDC20” as G2M genes ^110^, and the mitochondrial percentage of each cell was calculated with PercentageFeatureSet(pattern= “^MT-”). As an initial filter, poor-quality cells and doublets were first filtered considering the number of detected genes (e.g. < 8000 and > 1500) and mitochondrial percentage (> 5%). However, manual identification of poor-quality clusters was performed by adapting these threesholds.

Samples of equal temporal stages (and different in the case of zebrafish) were merged into one Seurat object, were subject to normalize and scale, and regress-out cell-cycle, mitochondrial percentage with the *SCTransform*() v1 method. Additionally, replicates were integrated to remove the potential batch effect associated with the merge of independent experiments with *FindIntegrationAnchors*(*reduction= “rpca”)* and *IntegrateData*() function. This procedure was also applied to zebrafish timepoints in order to obtain a more homogeneous integration of their cell types irrespective of maturation trajectories. From here, the “integrated” dataset is employed for downstream processing and visualization, gene expression levels were displayed from the “SCT” normalized assay, and SCENIC and SAMap comparative analysis employed raw counts (assay “RNA”).

### Cluster identification

To group cells by transcriptome similarity, expression data was linearly reduced into principal components (*RunPCA*, default parameters). A shared nearest neighbor (SNN) graph was calculated from the first principal components with *FindNeighbors*, default parameters. Based on this graph, the Louvain algorithm allows to detect communities or clusters with multi-level tuning of the resolution parameter (*FindClusters*, resolution (0.05, 0.1, 0.2, 0.3, 0.4, 0.5, 0.6, 0.8, 1.0, 1.2, 1.8, 2.4)). This resolution varied from lower values, to identify general cell classes (e.g. “neural stem cells”); to higher values, to identify cell types (e.g. “dorsal telencephalic neural stem cells”). This resolution parameter was optimized through iteration to obtain equivalent number of clusters or cell types despite cell number differences between species. Finally, cells were represented into a two-dimensional space by the non-linear dimensional reduction technique *Uniform Manifold Approximation and Projection* (*RunUMAP*) and also *t-Distributed Stochastic Neighbor Embedding (tSNE)*.

To identify the neurobiological cell identities of each cluster, differential expression analysis was carried out among clusters (*FindAllMarkers*, *min.pct = 0.25, logfc.threshold = 0.25*). *In situ* hybrization databases (Allen Brain Institute, Geisha Arizona, Mouse Genome Informatics, ZFIN) and literature articles were used to assign cell type identities based on the resulting marker genes. Additionally, our own spatial transcriptomic resource double confirmed the spatial location of cell types.

In all species’ datasets, we firstly filtered out cells not interesting for the neuroepithelium regionalization, such as erythrocytes (expressing *HBB*, *HBA* genes) and mesenchyme (*LUM*, *ALX1*)). As indicated previously, a first identification was conducted at low resolution, where cells were grouped in cell classes; and an annotation at high resolution, where only progenitors and neurons were clusterized and annotated into cell types. The gene dictionary employed for this task is available in the External Database S1.

### Comparative transcriptomic analysis

For the transcriptomic comparisons, four computational methods were carried out: Cell type mapping: ***Label Transfer*** ^111^ and ***SAMap*** ^63^; transcriptomic similarity: ***Gene Specificity Index* (*GSI*) Correlation** ^112^; and consensus cell type atlas: ***Seurat Integration*** (“RPCA” V4, Stuart et al., 2019). Additionally, quantitative pairwise comparisons were combined into a multi-method *bassi* plot^64^. This analysis is automatized in a “*conf.R”* file that contains all parameters and functions needed to generate the plots displayed in this article.

### Orthology and module filtering

All datasets underwent 1-to-1-orthology filtering as the state-of-the-art gene equivalency for the rest methods, except for the *SAMap* tool. The identification of paired 1-to-1-orthologues was carried out by *Ensembl* interface in R, *biomaRt* (v2.48.3). This tool downloaded gene identifiers in different notations (*external_gene_name, ensembl_gene_id_symbol*) and their respective paired orthologues (e.g. *with_mmusculus_homologue*, filtered by 1-to-1-orthology). To provide valid orthologues for gecko comparisons, *TransDecoder* ^113^ was used to predict CDS and protein sequences of all genes. Then, *OrthoFinder* ^114^(default parameters) compared gecko proteome (longest peptide for each gene) against the proteome of the other six species (longest peptide for each gene) that have been employed for inter-species transcriptomic comparisons:

- *Paroedura picta* (Madagascar ground gecko) (GeMoMa / TransDecoder)
- *Gallus gallus* (Chick) (Ensembl bGalGal1)
- *Mus musculus* (Mouse) (Ensembl GRCm39)
- *Homo sapiens* (Human) (Ensembl GRCh38.p14)
- *Danio rerio* (Zebrafish) (Ensembl GRCz11)
- *Lampetra fluviatilis* (Lamprey) (NCBI v7.0)

In case of additionally comparing gene ontologies, as TFs, the gene list matching with TFs GOs (*’GO:0006366’, ’GO:0000981’, ’GO:0003700’, ’GO:0006383’, ’GO:0000995’, ’GO:0001228’, ’GO:0001227’*) was obtained from *biomaRt* and datasets are additionally subset.

In both cases, raw counts and metadata were extracted from each *Seurat* object, 1-to-1-orthologues or gene modules were subset and transform into a common nomenclature if needed (by default, chick *external_gene_name*).

### GSI correlation

As firstly described by (Tosches et al., (2018b), the *Gene Specificity Index* (*GSI*) correlation starts from the raw count datasets, from which differential gene expression analysis was carried out to obtain a maximum of 400 markers per cell cluster. From these differentially expressed genes, only those in common for both species were selected. Finally, the raw counts datasets were filtered by this gene set and *gene specificity indexes* (*z-score*) are calculated. These indexes (𝑠*_g,c_*) were estimated as:

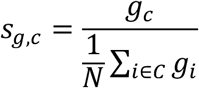

where 𝑔*_c_* is the average gene expression for a cluster or cell type, while 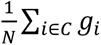, in the denominator, is the average expression for a gene across all cell types. From this matrix of indexes gene and cluster specific, a correlation matrix was obtained by the *Spearman’s* coefficient (ρ, *Rho*). To obtain the significance of these values, gene expression was shuffled 1000 times across cell types and the correlation was calculated. These values were used to calculate the significance, as the p-value was calculated as the fraction of the absolute Spearmann correlation that were greater than or equal to the absolute value of the non-shuffled data. Those with less than 5 percent (0.05) of aleatory greater correlations, out of the 1000 iterations, were marked as significant with a dot. The used visualization package for these matrices and their colour annotation is *ComplexHeatmap* ^115^. The scale color was fixed at -1-to-1 range for every comparison and plot presented, allowing direct comparison between graphs and species.

### Label Transfer

The Label Transfer tool ^111^ is integrated in *Seurat* utilities and analyses the similarity degree of cells based on their whole transcriptome. This method considers one of the datasets as the reference atlas, and the other as the query. After pre-processing the 1-to-1-orthologues datasets (*SCTransform, PCA, SNN, UMAP*…), cells from the query were mathematically projected into reference *PCA* embedding and are predicted cell type identities based on the chosen reference by *FindTransferAnchors* and *TransferData* functions. Although the transference was performed in both directions, we prioritized the larger dataset as reference for visualization.

With this weighted matrix, the algorithm predicted a new reference cell type out of each query cell. To visualize these predictions, we summarized into a matrix the percentage of cells of a query cell type predicted into certain reference cell type. The visualization package for these matrices is GSI code for *ComplexHeatmap* ^115^, adapting an extra layer to express the transference confidence related to each prediction as dot size. The scale color was fixed at maximum in 1 for every plot presented, allowing direct comparison between graphs and species.

### SAMap

The SAMap ^63^ method does not rely on 1-to-1-orthology between species, but stablishes the equivalencies between genes based on BLAST ^69^ protein similarity graph and their gene expression (map_genes.sh from https://github.com/atarashansky/SAMap). This tool overcomes the exiting limitation in clades with extra genome duplications, on which there are really few 1-to-1 orthologues. As this pipeline runs in *Python*, a function was created to automatize 1-to-1 comparisons and export the integrated object of *SAMap* in *h5ad* format, the coefficients of similarity between cell types and metadata of compared species. Afterwards, results were imported back into *R* for equivalent visualization with *Seurat* integrations. Moreover, the similarity coefficient provided by *SAMap* were plotted by *ComplexHeatmap* ^115^ in a similar fashion to the other cell type mapping tool, *LabelTransfer*. The scale color was fixed at maximum in 1 for every plot presented, allowing direct comparison between graphs and species.

### Multi-Method Combination

As described before^64^, pairwise comparisons can provide different results due to its intrinsic procedures. To obtain a more holistic picture of evolutionary comparisons, the *bassi* plot aims to combine them all (GSI Corr, Label Transfer and SAMap) into a single plot. For this purpose, the outputs of these individual methods are computed and plotted as the mean across methods. In the case of GSI Corr, the value used for the mean is the original correlation minus the median of the cell type’s row. Confidence is represented by circle size, being this proportional to the standard deviation of the mean across three methods. Finally, a significance label is inserted inside the circle if the pairwise method for a specific cell type is among the 95^th^ percentile of its row values. This significance label is specific of each method: square (GSI Correlation), circle (label transfer) and triangle (SAMap).

### Multi-Species Integration

For this approach, each species dataset was integrated as a replicate, but then equal number (smallest dataset number) of cells was randomly selected. After 1-to-1 ortholog filtering and gene name conversion, datasets were integrated into a joint Seurat object by *SCTransform* ^116^. The mathematical method for batch effect removal was the *Reciprocal Principal Components Analysis* (*RPCA*), conceived to be more conservative and faster than *Canonical Correlation Analysis* (*CCA*). All parameters were set by default, except for the following ones in: *FindIntegrationAnchors (reduction= “rpca”, normalization.method = “SCT”, dims = 1:50)*, and equivalently to the rest of the pipeline.

After the re-annotation of clusters in this embedding, we assessed the conservation of a specific cell type regarding the proportion of cells of each species. Moreover, the raw expression object was further analysed by extracting a specific signature for these integrated cell types.

#### Direct RNA In situ sequencing (ISS)

### Probe Selection

To select informative genes and select probe panels for chicken embryonic brains, we used clustered non-integrated data from (insert here the relevant sources and embryonic stages).

Padlock probes against the selected genes were designed using the Oligo Designer Toolsuite (https://oligo-designer-toolsuite.readthedocs.io/en/latest/) following the SCRINSHOT Probe design workflow (REF https://pubmed.ncbi.nlm.nih.gov/33216742/). The resulting probes were then converted into ISS probes using a custom script to replace the padlock backbone with gene-specific barcode sequences and common primer binding sequences, while maintaining the SCRINSHOT ligation chemistry.

Padlock probes were ordered as phosphorylated DNA Ultramers oPools Integrated DNA Technologies (IDT), with a synthesis scale of 50 pM, resuspended in IDTE buffer at a 1 uM/probe concentration. The sequence of all padlock probes (PLPs) is available in Datafile S1.

### Library Preperation

A detailed step-by-step protocol for the version of directRNA-ISS used is provided trough the following link: https://www.protocols.io/view/home-made-direct-rna-detection-kqdg39w7zg25/v1?version_warning=no

Tissue samples (stored at -80°C were allowed to reach RT, washed with PBS, fixed with 3,7% formaldehyde for 30 min and washed with PBS. The tissues were permabilised by incubating them in 1% SDS for 2 min, washed in PBS and incubated in 70% methanol on ice for 60 min, followed by a last PBS wash. The tissues were then progressively dehydrated using one wash each of 2 min in 70% ethanol and 100% ethanol. SecureSeal™ Hybridization Chambers were applied around the tissue sections. The simples were then washed with PBS-tween 0,1%. Next, phosphorylated padlock probes were hybridized at a final concentration of 10 nM/PLP in hybridization buffer (2X SSC, 10% formamide,) overnight at 37°C. Excess probes were washed with 2 washes in wash buffer (10% formamide, 2X SSC) and a ligation was performed at 37 °C for 2 hours with 0.25 U/uL of SplintR ligase in a home-made buffer whose composition is indicated in the link above. Rolling circle amplification (RCA) was then performed with phi29 polymerase overnight at 30°C. L-probes were hybridized at a concentration of 100 nM for 1 hour at room temperature in L-probe hybridization buffer (4X SSC, 40% formamide), followed by the hybridization of the detection oligos (100 nM) and DAPI for 1 hour at room temperature in the same hybridization buffer. Sections were washed with PBS and mounted with SlowFade Gold Antifade Mountant.

### Image acquisition

The mounted slides were then imaged using a Leica DMI6000 epifluorescence microscope equipped with an external LED light source (Lumencor® SPECTRA X light engine) and an automatic multi-slide stage (LMT200-HS). The camera used was a sCMOS camera (Leica DFC9000 GTC). All images were acquired using a 40X water objective (HCX PL APO 40X/1.10 W CORR).

Multispectral images were captured with microscope equipped with filter cubes for 6 dye separation and an external filter wheel (DFT51011).

Image acquisition was performed by outlining regions of interest (ROIs) that were saved and reused for cycling imaging. The image of the ROIs was captured with 10% overlap where the Z-stacks of the images had a total thickness of 12μm and 0.5 µm distance between them to cover the depth of the tissue.

### Image analysis

The raw images and the metadata from the microscope were fed into an in-house preprocessing software module (https://github.com/Moldia/ISS2025/tree/main/ISS_preprocessing). The module performs the following steps: first the images are deconvolved using a RedLionFish based pipeline (https://github.com/rosalindfranklininstitute/RedLionfish) that also maximum-projects all the z-stacks belonging to individual tiles and channels. The projected images are then simultaneously stitched and aligned across imaging cycles, using the ASHLAR algorithm (21). The aligned stitched images are sliced into smaller tiles, to allow a computationally efficient decoding.

The resliced images were then fed into an in-house decoding software module (https://github.com/Moldia/ISS2025/tree/main/ISS_decoding), which converts them into SpaceTX format (REF Starfish), and pipes them into the Starfish Python library for decoding of image-based spatial transcriptomics datasets (https://github.com/spacetx/starfish). In this decoding module, the images are normalized across the channels and the imaging cycles, the ISS spots are identified and decoded, and quality metrics are computed.

All the downstream analysis was performed using the code in this repository (https://github.com/Moldia/ISS2025/tree/main/ISS_postprocessing). The decoded gene expression spots were segmented using the Baysor algorithm (https://kharchenkolab.github.io/Baysor/dev/segmentation/), that generated a segmentation polygon .json file. This file was then translated into a segmentation mask (.npz file) that was applied onto the ISS data. This segmentation mask and the decoded spots table, as well as the reference scRNAseq dataset, were passed as inputs to a Python implementation of Probabilistic Cell Typing (25), as described in the section below.

ISS data analysis:

In order to integrate the ISS data with the pre-existing single-cell RNAseq datasets, and to map in-situ the scRNAseq clusters, Probabilistic Cell Typing (PCIseq, (25)) was used. A cluster-by-gene average expression matrix inferred from the clustered scRNAseq data was fed, together with a segmentation mask and the complete ISS dataset, to the PCIseq algorithm, following the implementation at https://github.com/Moldia/ISS2025/blob/main/ISS_postprocessing/Notebooks/PCIseq.ipynb. After the assignment, low probability cells were eliminated (p<0.x) and the remaining cells were plotted, cluster by cluster, over a DAPI image to analyze their anatomical distribution. A total of xxx cells were sequenced and analyzed.

### Single nuclei ATAC sequencing (snATACseq) Nuclei isolation

Mouse and chick brains were dissected following the same experimental procedure detailed before, but nuclei extraction was performed by osmotic shock. For this purpose, brains were placed in a hypotonic medium - and mechanical disaggregation - micro-pestle (10 times) and micropipette up-and-downs (10 P100, 10 P100+P2 tip). Then, 10 minutes incubation in ice led to complete membrane disruption. Nuclei were isolated by centrifugation (500g for 5 min), and resuspended in the aimed final concentration (5000 cells per µl in nuclei buffer - recommended by *10X Genomics*). Nuclei concentration and chromatin quality was evaluated manually in a Neubauer chamber after mixing with 1:1 trypan blue.

The wet lab part of 10X Genomics snATAC pipeline was performed following their protocols. Briefly, nuclei suspensions were incubated in a Transposition Mix (Tn5 Transposase). Then, 24,000 nuclei were loaded per reaction into the Chromium Controller 10X (10X Genomics) (Droplet-based method) for a nuclei recovery of 12,000 nuclei/ Single nuclei library was prepared using “Chromium Next GEM Chip H Single Cell Kit”, 16 rxns PN-1000162 or and “Chromium Next GEM Single Cell ATAC Kit v2”, 4 rxns PN-1000406 and “Single Index Kit N, Set A” 96 rxns PN-1000212, following “Chromium Next GEM Single Cell ATAC Reagent Kits v2” (Document number CG000496).

Similarly to scRNA-seq, barcoded fragments were generated for each bead-cell (Gel Beads-in-emulsion (GEMs)). For this step, oligonucleotides containing (i) an Illumina P5 sequence, (ii) a 16 nt 10x Barcode and (iii) a Read 1 (Read 1N) sequence were released in each GEM and mixed with DNA fragments and Master Mix. After incubation and amplification, the GEMs were broken and pooled. Next, P7 and a sample index were added during library construction via PCR.

To assess the quality of libraries, DNA concentration was calculated using Qubit dsDNA HS DNA Kit (Thermo Fisher Scientific, Cat. # Q32854) and fragment length was visualized on an Agilent 2100 Bioanalyzer using Agilent High Sensitivity DNA kit (Agilent Technologies, Cat. # 5067-4626). The obtained libraries were sequenced on a NovaSeq 6000 for an approximated 50.000 reads per cell.

### Single nuclei epigenomic pre-processing

For alignment and demultiplexing, *cellranger-atac-2.1.0* and mm10-2020-A and bGalGal1 (Ensembl 112) genomes. The alignment results and quality control are displayed in the Table S03. When multiple replicates, peaks were merged into consensus as indicated in *Signac*^68^ vignettes, https://stuartlab.org/signac/articles/overview. Briefly, *bed* peaks were imported as *GenomicRanges*, and then combined and filtered. The single cell metadata was associated with peaks and fragments objects created, then feature matrix created and annotations combined into a chromatin assay of the individual replicates that will be merged into the final merged *Signac* object (*Signac* v1.13).

A quality control (QC) assessment is performed to exclude low-quality nuclei. This step included filtering by *NucleosomeSignal*()  < 4), *TSSEnrichment*() (>2) – snATAC-seq specific - and number of feature counts (*nCount_ATAC* < 150000 & *nCount_ATAC* > 2750) - directly comparable to scRNA-seq QC metrics. Additionally, DoubletFinder pipeline was also included to filter possible doublets (0.075% likelyhood). Normalization and dimension reduction was carried ou by *FindTopFeatures*(), *RunTFIDF*(), *RunSVD*(), *RunUMAP*(*reduction = “lsi”, dims = 2:50*), *FindClusters(algorithm = 3, resolution = c(0, 0.5, 1, 2)*.

Open promotor regions upstream a gene were considered a proxy for gene expression. RNA expression was predicted with *GeneActivity*() function and stored as “*RNA*” assay. Normalization (*SCTransform*), *FindVariableFeatures*() and dimension reduction (*RunPCA*) was also performed in this RNA assay and transfer labels (*LabelTransfer*) with scRNA-seq equivalent experiment (original dataset, “*integrated*” assay). Based on these predicted labels, several cell filtering rounds were performed to finally obtain an equivalent scATACseq object to the scRNAseq atlas of mouse and chick, respectively.

### Pseudocells – Multiomic atlas

As a previous step to gene regulatory network inference, scRNAseq and snATACseq (“RNA” prediction) datasets were integrated by *FindIntegrationAnchors(method = “cca”) and IntegrateData.* Both objects were subsampled if needed to obtain the same cell/nuclei number in both datasets. Then, clusters of 10 cells/nuclei in a 5:5 proportion was established based on UMAP cell embeddings and the *nn2* function of *RANN* in *R*. A cell/nuclei could be included in more than one pseudocell to ease the ATAC and RNA proportion.

### SCENIC+ - eRegulon calculation

The calculation of eRegulons –gene networks associated with specific TFs– was performed following *SCENIC*+ pipeline^78^ created by Aerts’ lab (KU Leuven). Employed notebooks, github stored, are adapted from their tutorials.

Initially, snATAC was processed with *pyCisTopic* default parameters to process snATACseq to obtain accessibility regions and topic models – equivalently to *Signac*. The optimal number selected was 25 for the mouse, 45 for the chick. Next, candidate enhancer regions are inferred by binarization of region-topic probabilities; and then, calculating *differentially accessible regions* (*DARs*) per cell type. *DARs* are calculated based on the imputed region accessibility (*cistopic_imputed*), but we will keep a non-imputed region accessibility matrix stored for later processing.

For motif enrichment calculation, the *pyCisTarget* wrapper was employed on previously obtained topics. Precomputed motif scores for genomic regions were downloaded from Aerts’ repositories for mouse, and kindly provided by Dr. Nikolai Hecker from Aerts’ laboratory for chick by computing our snATAC-seq peaks with their adult snATAC-seq chick dataset^82^. Motif-topic assignment led to *cistromes*, set of target regions for a specific binding site or motif linked to a TF.

RNA and ATAC profiles were aggregated within each pseudocell, and this multi-omic dataset was inputted to SCENIC+ pipeline. Default gene and region search space for mouse (v114, stable), custom annotation for *bGalGal1* for the chick (Ensembl bGalGal1, custom chromosomes, not updated to bGalGal1 for UCSC). Then, relationships between elements in GRNs are inferred: TF, region and gene with default parameters. Statistical inference of these relationships resulted into eRegulons, which are stored as eRegulon metadata and stored here as External Database S2.

### Evolutionary comparisons of GRNs

#### Cross-tabula comparisons

Previously identified eRegulons and differentially expressed genes from both species were compared using set operations (intersection and diffSet functions) implemented in the provided R code ().

#### LiftOver of cisRegulatory element (CRE)

Regulatory regions obtained in SCENIC+ eRegulons were converted into chick genomic regions (bGalGal1) as Granges and lifting them to mouse genome (mm10) using *liftOver*^117^ (GCF_016699485.2ToMm10.over.chain). Regions with only one corresponding region (1-to-1 orthologue regions) showed inferior lengths to 100 bp, not achieving original enhancer-length 700-800 bp on mean. Similarly occurred to differential accessible regions calculated by *Signac*.

#### PhastCons of cisRegulatory element (CRE)

Regulatory regions obtained in SCENIC+ eRegulons were exported into *BED* files in *R* through *rtracklayer* (v1.60.1) and windowed into 100 bp frames with *bedtools* ^118^ . *phastCons* genome scores were imported from UCSC repository for mm10 and galGal6, as bGalGal1 is not available – liftOver of bGalGal1 regions was performed, followed by 1-to-1 orthologue regions filtered. Then, regulatory regions BEDs were parsed over phastCons scores by *bigwigAverageOverBed*. Regulatory regions metadata (associated gene or TF) was further employed for eRegulons evolutionary comparisons and functional enrichment analysis by *clusterProfiler* ^119^ / compareClusters() function.

#### Co-expression networks – hdWGCNA

Co-expression network was performed as described in hdWGCNA vignettes^120,121^. Both early brain and pallial developmental raw scRNAseq datasets were processed with Seurat, but by *FindVariableFeatures(), NormalizationData()*, *ScaleData()* and *RunHarmony()* functions. Then, pseudocells were generated and signed weighted networks were constructed using soft-thresholding to approximate scale-free topology. Gene co-expression modules were identified from the topological overlap matrix by hierarchical clustering and dynamic tree cutting, and summarized by their eigengenes.

Functional enrichment of gene modules is performed within the hdWGCNA framework by enrichR package. Resultant enrichment tests provide functional terms associated to each gene module, but also a significance value regarding its enrichment among modules. Therefore, functional terms that are not module-specific can be filtered by adjusted p-values < 0.05. Nonetheless, each gene can only belong to a specific module, so even non-module specific functional terms are carried out by module-specific gene sets.

To assess evolutionary conservation, we performed cross-species transference and module preservation using the *modulePreservation* function in hdWGCNA. Modules with Zsummary > 10 were considered highly preserved, 2–10 moderately preserved, and <2 non-preserved; but other parameters such as *medianRange* and *Z.connectivity* were assessed to avoid module size bias^122^.

### Gene metrics

For calculating these metrics, we used available data that compile the different parameter scores for every gene: supplementary table 3 and 9 Cardoso-Moreira et al. (2019)^21^ and International Mouse Phenotyping Consortium ^91,123^. The code employed for both analysis is available on the GitHub site and its raw version was provided by Dr. Xuefei Yuan from Prof. Henrik Kaessmann’s laboratory.

Pleiotropy, defined as the breadth of gene expression across tissues and time points, was assigned to each expressed gene according to Cardoso-Moreira et al, (2019) / Ensembl 69 ^21^. In our calculations, we employed their coefficient scores available in their supplementary data to estimate the pleiotropy of certain cell type or gene set.

The impact of gene loss defined as essentiality or lethality was calculated based on Essential Genes Data from the International Mouse Phenotyping Consortium (IMPC)^91,123^. Regarding the Full Spectrum of Intolerance to Loss-of-function (FUSIL) categorisation, we considered Cellular Essential (CL) and Developmental Essential (DL) as lethal; and other subviable and viable as general non-essential.

The code employed for mean pleiotropy and lethality per cell is available on the GitHub site and its raw version was provided by Dr. Xuefei Yuan from Prof. Henrik Kaessmann’s laboratory. Module pleiotropy and lethality was calculated similarly, dropping off genes without pleiotropy or lethality value.

### Data availability

The mouse scRNA-seq data was downloaded from http://mousebrain.org/development/downloads.html (Linnarsson’s laboratory) in *loom* format (dev_all.loom), subset for E9.5 for the early brain and for dorsal telencephalon for developmental comparisons. Filtered *loom* object was transformed to a count matrix by *loomR* into *Seurat*/*R.* Likewise, the scRNA-seq data from Braun et al (2024) was downloaded from https://github.com/linnarsson-lab/developing-human-brain in *loom* and CS13 stage was subset. Developmental pallium scRNA-seq data from Zaremba et al. (2025) was downloaded from heiDATA repository (https://heidata.uni-heidelberg.de/dataset.xhtml?persistentId=doi:10.11588/DATA/BX6REK). scRNAseq zebrafish data from Raj et al., (2020) was downloaded as indicated in https://github.com/brlauuu/zf_brain and 16/18/20 hpf stages were subset. All data and code will be publicly available upon publication (https://github.com/rodrisenovilla/Senovilla-Ganzo2025).

## Supporting information

Supplemental Table 1. Gene catalog

Supplemental Table 2. Eregulon metadata

## Acknowledgements

We thank current and past members of the Garcia-Moreno group, members involved in Kaessmann group and those many researchers who have discussed the data with our team over these years: especially M. Cardoso-Moreira (The Crick Institute, UK), J.L Ferran (Universidad de Murcia, Spain) and N. Irie (RIKEN, Japan). We gratefully acknowledge support from E. Vazquez (Universidad de Zaragoza, Spain) and L.J. Chueca (BC3, Spain) with genome annotation, and N. Hecker (VIB-KU, Belgium) with SCENIC+, and X. Yuan (ZMBH, Germany) for cell gene parameter code.

## Funding sources

This work was funded by a Ikerbasque Research Fellowship to F.G.-M.; Spanish Ministry grants MICINN PGC2018-096173-A-I00 and MICINN PID2021-125156NB-I00 to F.G.-M; Basque Government grants PIBA 2020_1_0057 and PIBA_2022_1_0027 to F.G.-M.; and a Fundacion Tatiana predoctoral fellowship to R. S.-G. CIC bioGUNE support was provided by the Department of Industry, Tourism and Trade of the Government of the Autonomous Community of the Basque Country (Emaitek and Elkartek Research Programs 2015-2023, KK-2020/00008), the Innovation Technology Department of the Bizkaia County, and by the Spanish Ministry of Science and Innovation grant MCIN/AEI/10.13039/501100011033 (PID2020-118464RB I00 and Severo Ochoa Excellence Accreditation CEX2021-001136-S) to A.M.A.

## Author contributions

R.S-G and F.G.-M. conceived the project, interpreted the data, designed and performed experiments and data analysis, and wrote the manuscript with input from all authors. C.B. performed embryo work and experiments. A.M.A., B.Z. and R.S-G. performed scRNA-seq and snATAC-seq. C.B., B.Z., T.Y.,M.G. and R.S.-G. performed bioinformatic analysis. C.B. and M.G., performed and analyzed the *in situ* sequencing experiments. M.N., H.K. and F.G.-M. provided reagents.

## Fig. Supplementary

**Figure S01.**
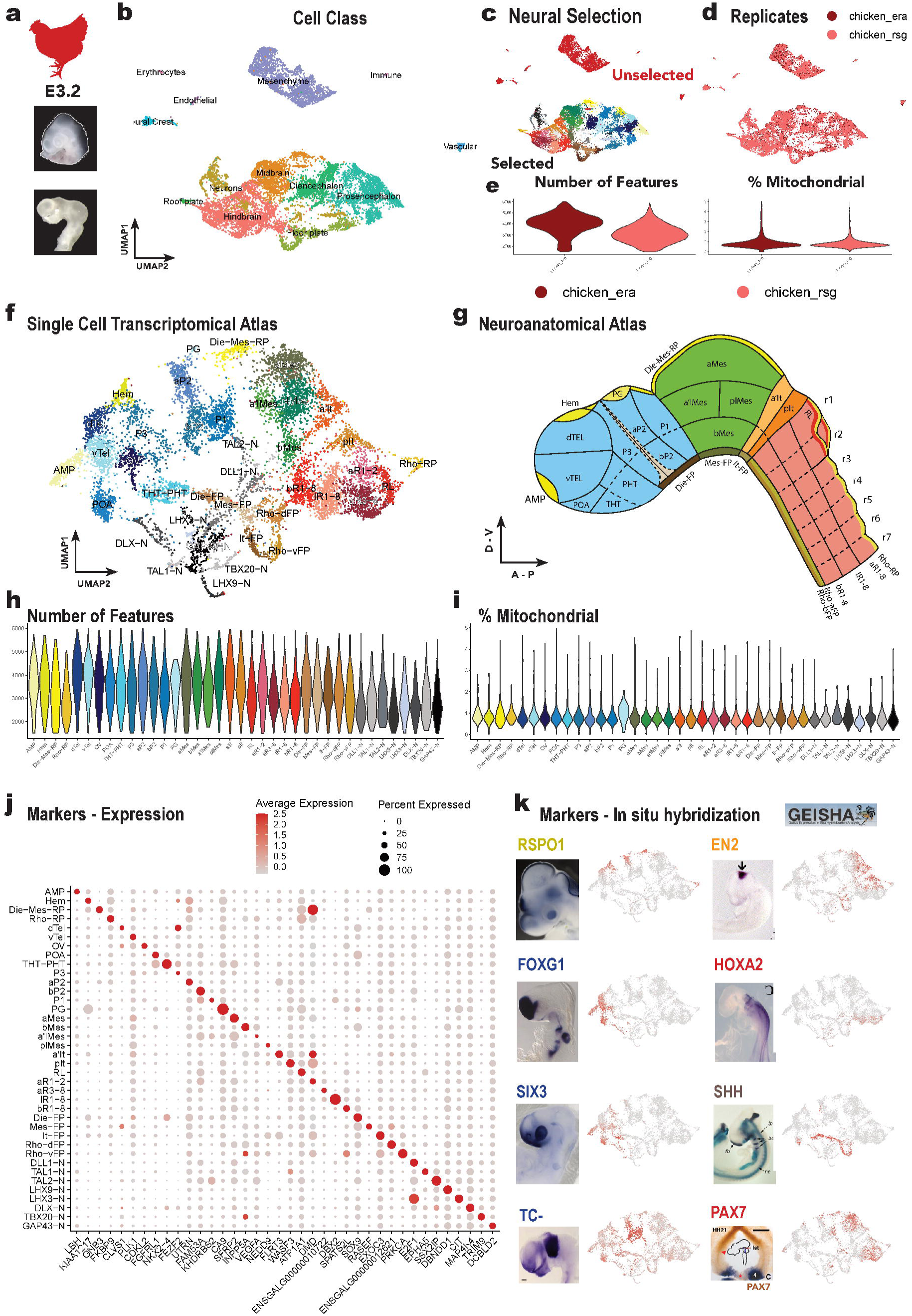
Chick - Single cell atlas quality control and processing. **a**, Head morphology before and after brain dissection (*in house*) of E3.2 chick. **b**, Unfiltered datasets composed by all neural and non-neural cell grouped by cell classes. **c**, Neural selection coloured by their final cell type annotation, unselected cells in blue. **d**, Technical replicates plotted over the all cells UMAP. **e**, Quality control variables (Feature number and mitochondrial percentage) plotted by replicate. **f-g**, Final single cell transcriptomic atlas and respective neuroanatomical atlas filtered by neural cells. **h-i**, Quality control variables (**h**, feature number; and **i**, mitochondrial percentage) plotted by final cell type. **j**, Marker genes obtained from FindAllMarkers() function (Top1 per cell type). k, Examples of *in situ* hybridization assays in chick of several marker genes from GEISHA database^124^ to annotate each cell type and the corresponding FeaturePlot().

**Figure S02.**
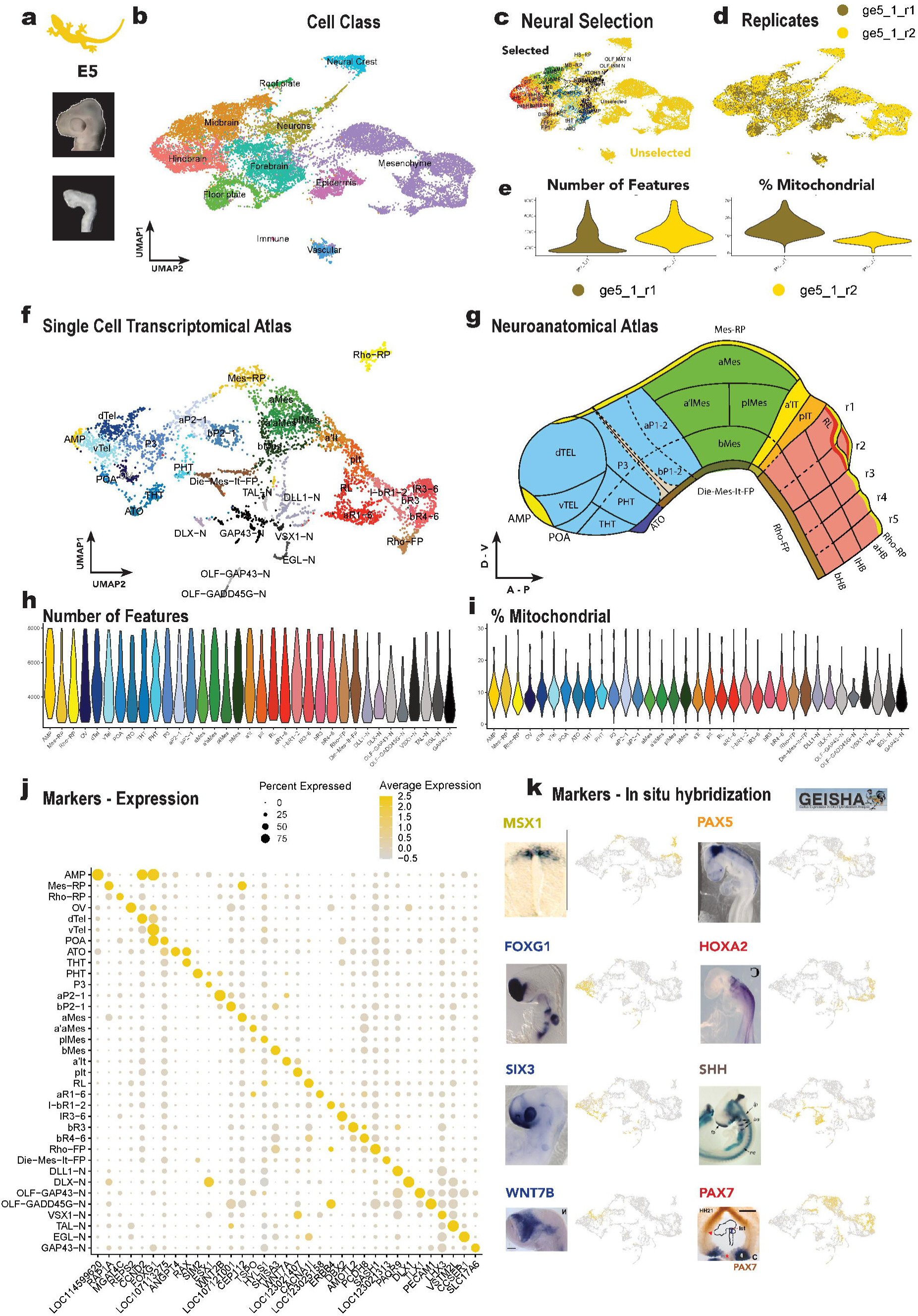
**Gecko - Single cell atlas quality control and processing**. **a**, Head morphology before and after brain dissection (*in house*) of E5 gecko. **b**, Unfiltered datasets composed by all neural and non-neural cell grouped by cell classes. **c**, Neural selection coloured by their final cell type annotation, unselected cells in blue. **d**, Technical replicates plotted over the all cells UMAP. **e**, Quality control variables (Feature number and mitochondrial percentage) plotted by replicate. **f-g**, Final single cell transcriptomic atlas and respective neuroanatomical atlas filtered by neural cells. **h-i**, Quality control variables (**h**, feature number; and **i**, mitochondrial percentage) plotted by final cell type. **j**, Marker genes obtained from FindAllMarkers() function (Top1 per cell type). **k**, Examples of *in situ* hybridization assays in chick of several marker genes from GEISHA database^124^ to annotate each cell type and the corresponding FeaturePlot().

**Figure S03.**
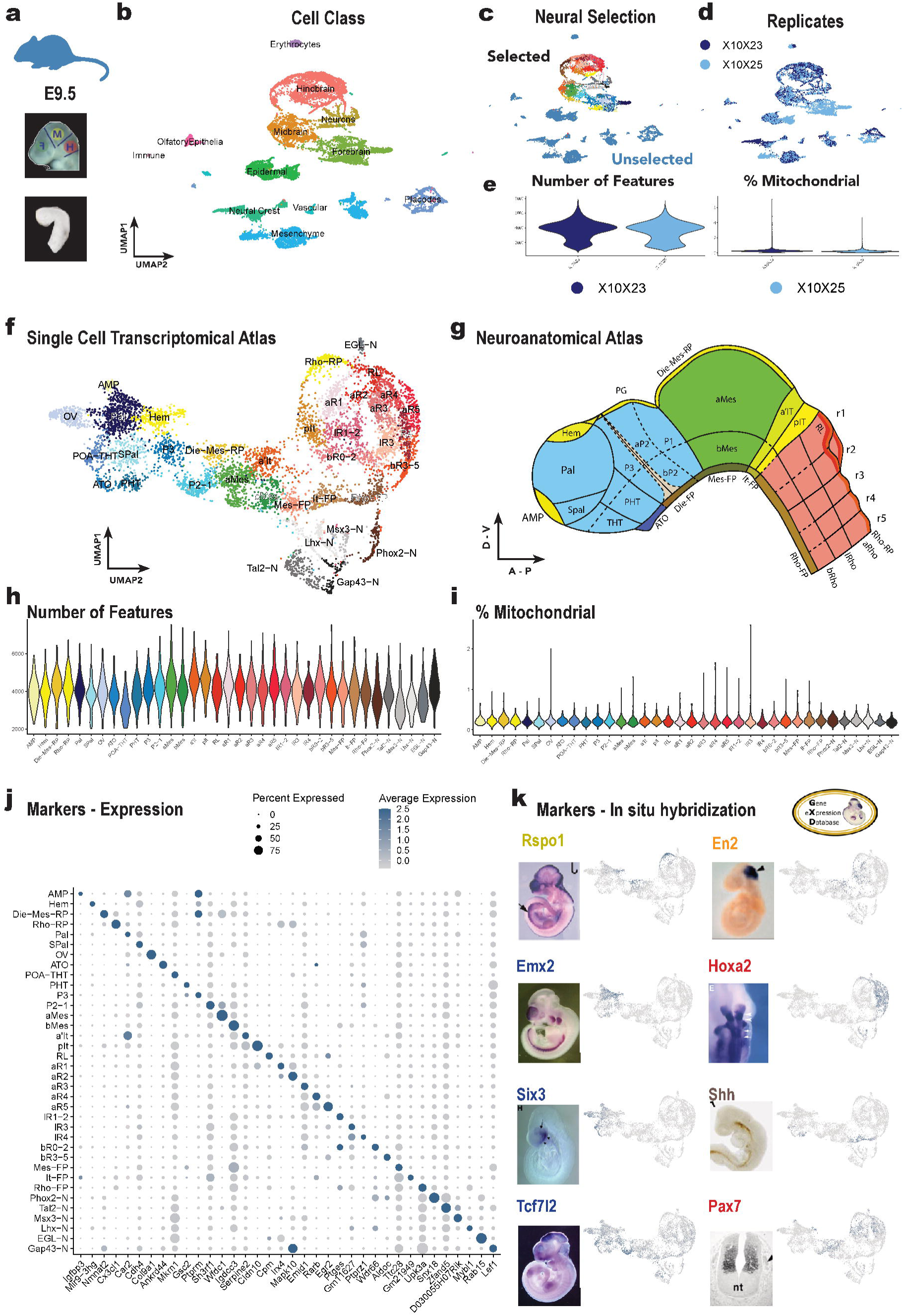
Mouse - Single cell atlas quality control and processing. **a**, Head morphology before^40^ and after brain dissection brain (*in house*) of E9.5 (TS11) mouse. **b**, Unfiltered datasets composed by all neural and non-neural cell grouped by cell classes. **c**, Neural selection coloured by their final cell type annotation, unselected cells in blue. **d**, Technical replicates plotted over the all cells UMAP. **e**, Quality control variables (Feature number and mitochondrial percentage) plotted by replicate. **f-g**, Final single cell transcriptomic atlas and respective neuroanatomical atlas filtered by neural cells. **h-i**, Quality control variables (**h**, feature number; and **i**, mitochondrial percentage) plotted by final cell type. **j**, Marker genes obtained from FindAllMarkers() function (Top1 per cell type). **k**, Examples of *in situ* hybridization assays of several marker genes from mouse GXD database^127^ to annotate each cell type and the corresponding FeaturePlot().

**Figure S04.**
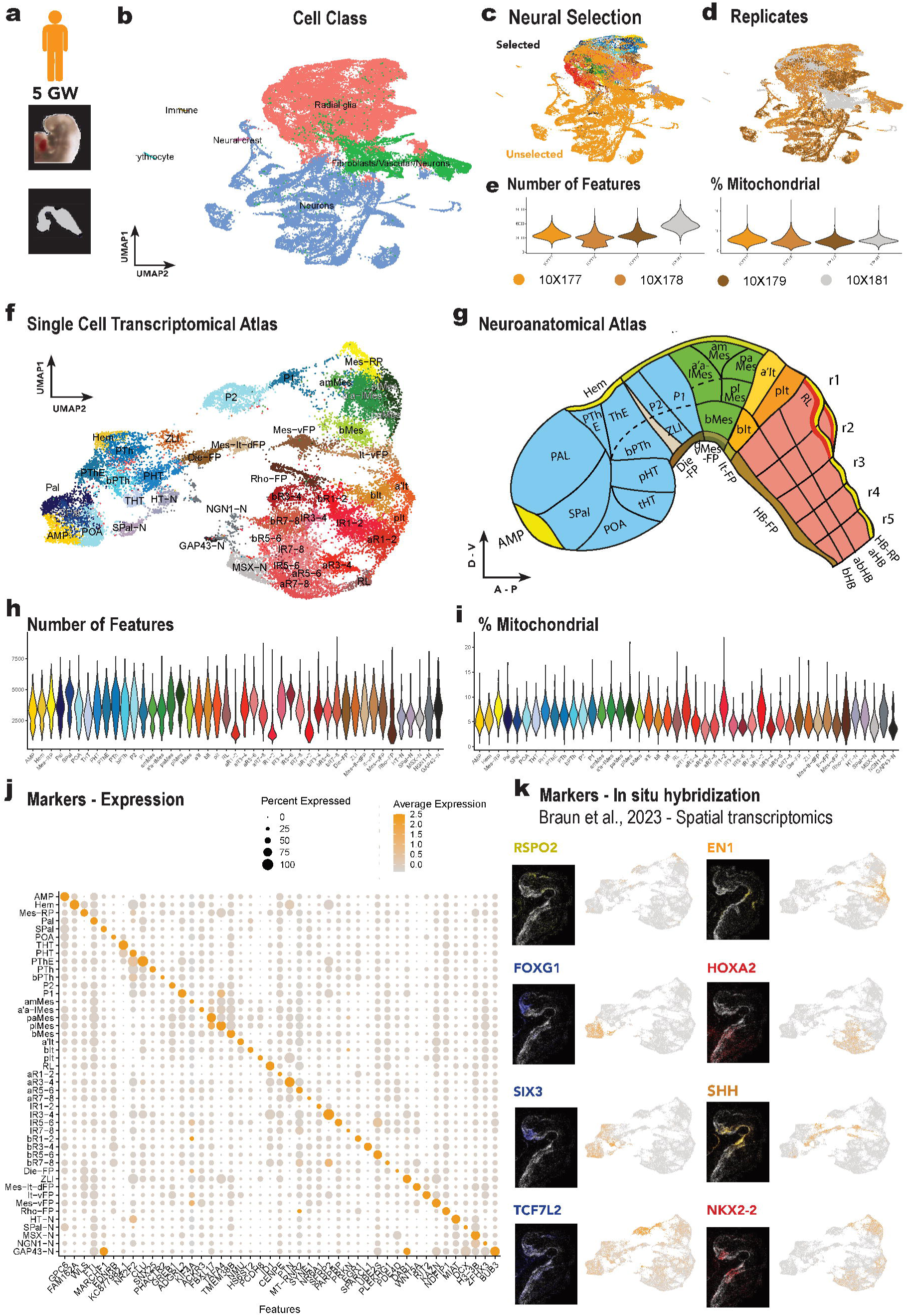
Human - Single cell atlas quality control and processing. **a**, Head morphology before and after brain dissection brain^41^ of 5GW (CS13) human. **b**, Unfiltered datasets composed by all neural and non-neural cell grouped by cell classes. **c**, Neural selection coloured by their final cell type annotation, unselected cells in blue. **d**, Technical replicates plotted over the all cells UMAP. **e**, Quality control variables (Feature number and mitochondrial percentage) plotted by replicate. **f-g**, Final single cell transcriptomic atlas and respective neuroanatomical atlas filtered by neural cells. **h-i**, Quality control variables (**h**, feature number; and **i**, mitochondrial percentage) plotted by final cell type. **j**, Marker genes obtained from FindAllMarkers() function (Top1 per cell type). **k**, Examples of FISH experiments in human CS13 from Braun et al. (2023)^41^ and the corresponding FeaturePlot().

**Figure S05.**
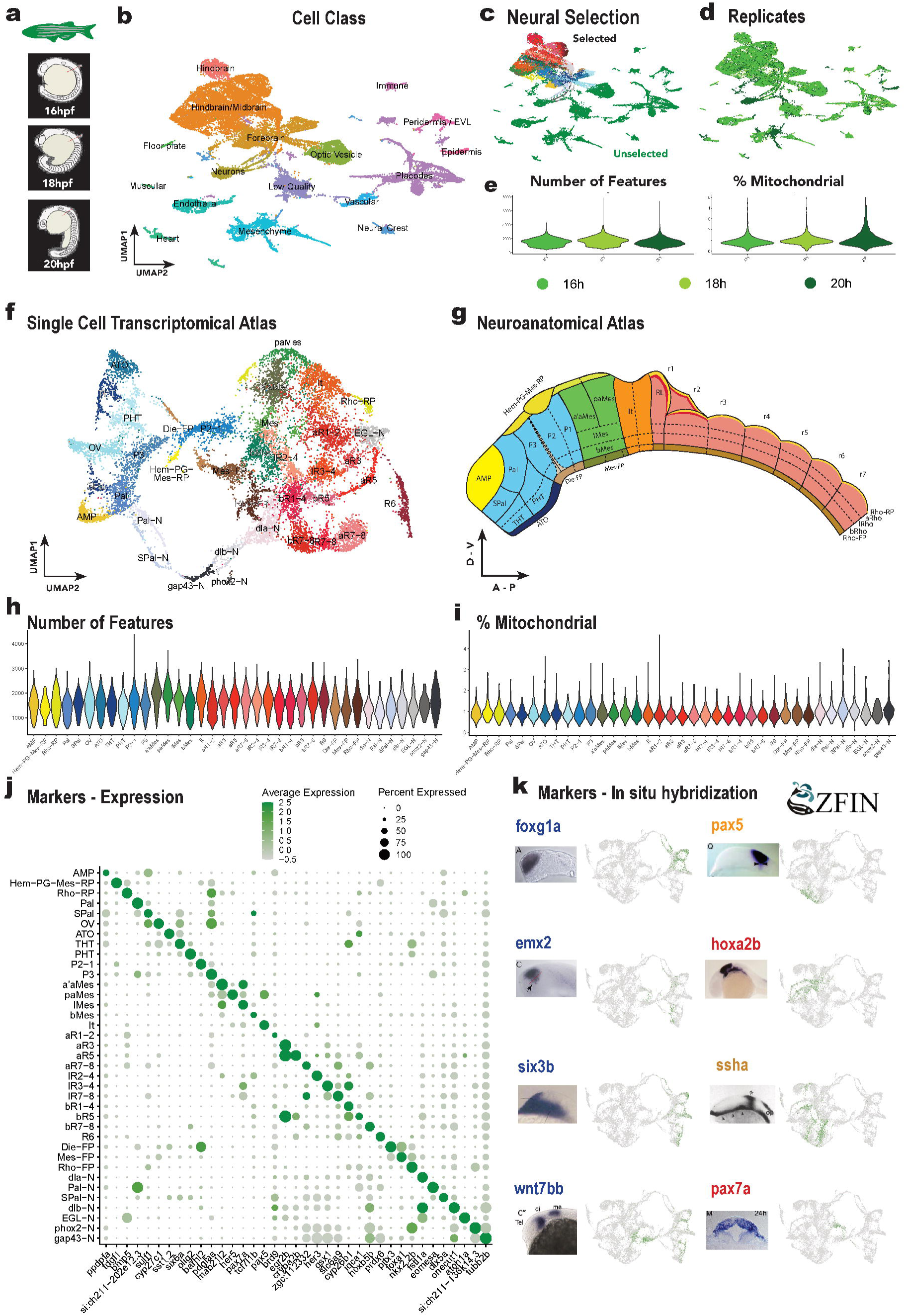
Zebrafish - Single cell atlas quality control and processing. **a**, Embryo morphology at three selected stages of zebrafish development (16, 18 and 20 hpf). **b**, Unfiltered datasets composed by all neural and non-neural cell grouped by cell classes. **c**, Neural selection coloured by their final cell type annotation, unselected cells in blue. **d**, Stage datasets plotted over the all cells UMAP. **e**, Quality control variables (Feature number and mitochondrial percentage) plotted by stage. **f-g**, Final single cell transcriptomic atlas and respective neuroanatomical atlas filtered by neural cells. **h-i**, Quality control variables (**h**, feature number; and **i**, mitochondrial percentage) plotted by final cell type. **j**, Marker genes obtained from FindAllMarkers() function (Top1 per cell type). **k**, Examples of *in situ* hybridization assays of several marker genes from zebrafish ZEFIN database^129^ to annotate each cell type and the corresponding FeaturePlot().

**Figure S06.**
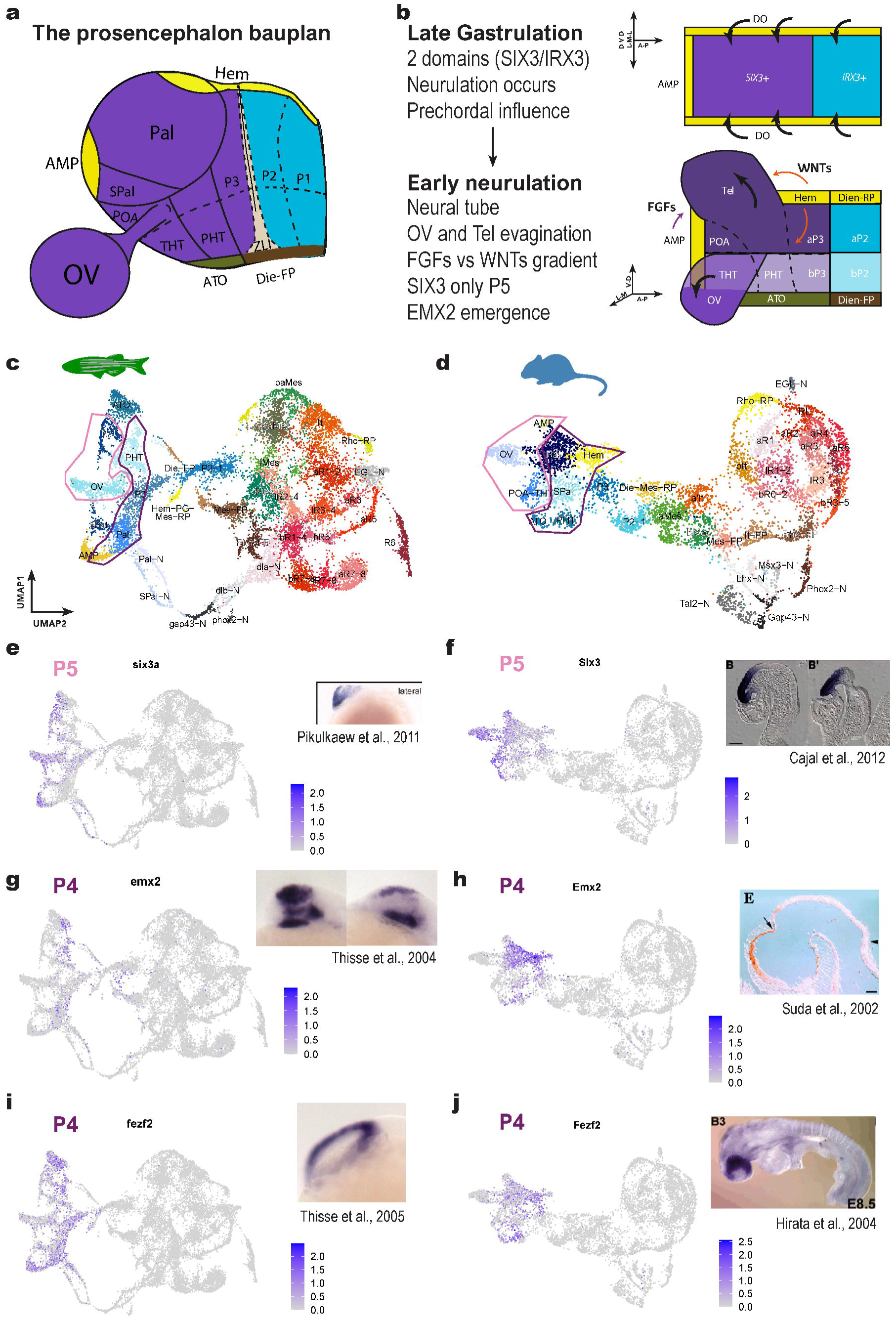
Secondary prosencephalon antero-posterior segmentation: drivers and cell types. a, The prosencephalon bauplan, neuroanatomical scheme of the patterning identities of the phylotypic brain. Secondary prosencephalon in purple, diencephalon in blue, dorsal organizers in yellow, floor plates in brown (except acroterminal organizer in kaki) and zona limithans intrathalamica in grey. **b**, Morphogenetic hallmarks in prosencephalon development. From late gastrulation of the ancestral secondary prosencephalon and diencephalon distinction (SIX3+ purple, IRX3 sea blue); to early neurulation with optic vesicle’s lateral evagination and dorsal telencephalic evagination. Black arrows on top indicate the folding of the neuroectoderm, coloured arrowns on bottom follow morphogenesis and morphogen gradients (evagination in black; Wnt, orange; and FGFs, purple). Medial-Lateral (M-L) is equivalent to the prospective Dorso-Ventral (D-V), Antero-Posterior (A-P) is on the X axis of the scheme. **c-d**, Final UMAP atlases of zebrafish (**c**) and mouse (**d**), where cell types from each secondary prosencphalon segment (prosomere) are circled in their respective color (pink, P5; purple, P4). **e-j**, Gene expression plots for P5 marker six3a/Six3 (**e,f**) and P4 markers emx2/Emx2 (**g,h**) and fezf2/Fezf2 (**i,j**). Each expression plot is accompanied by an *in situ* hybridization or immunohistochemistry of each gene in the correspondent species^130–135^.

**Figure S07.**
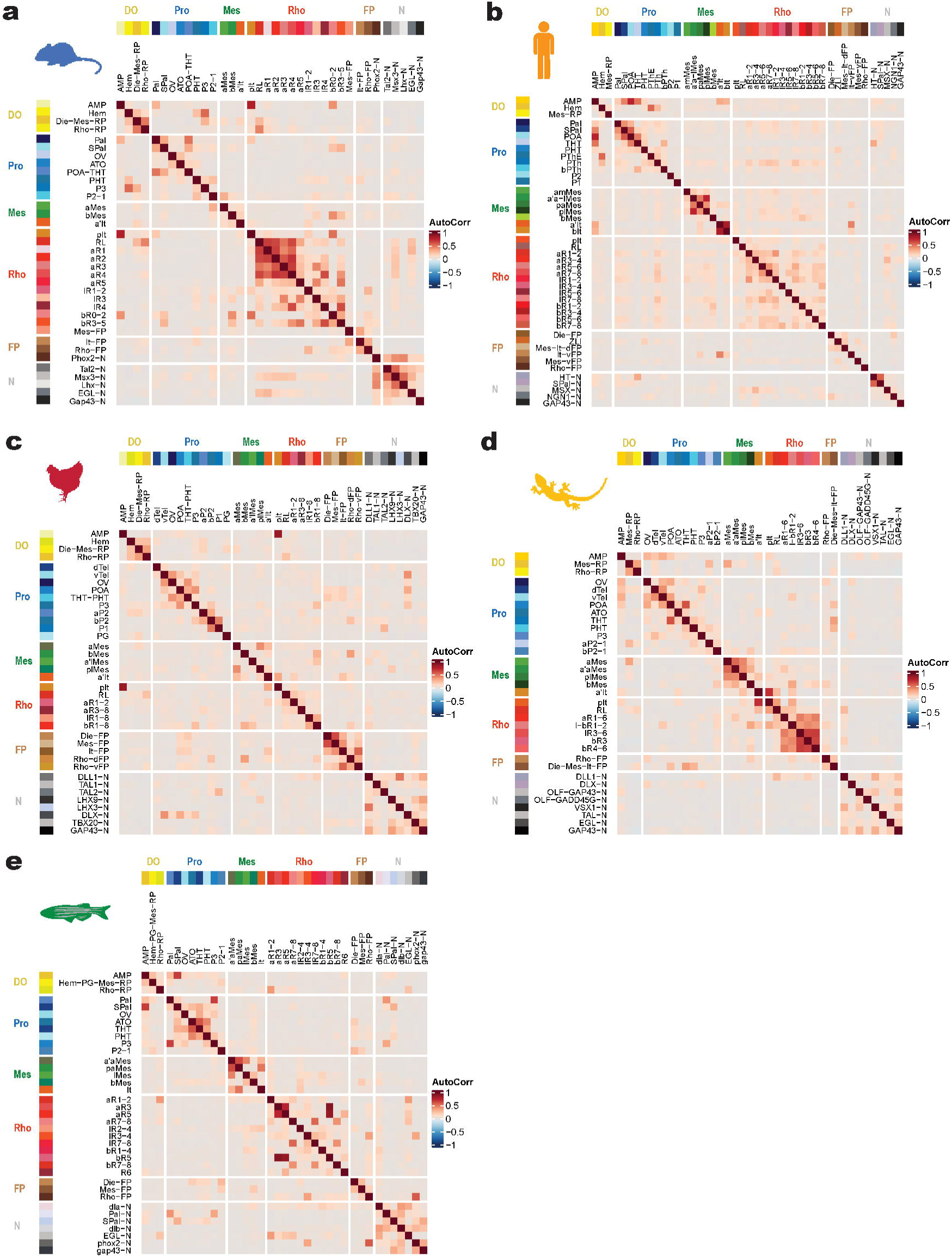
Cell type correlation within every species’ atlas. Mouse (**a**), human (**b**), chick (**c**), gecko (**d**) and zebrafish (**e**). Colors match to correlation values. Annotation color bars in both axes matches cell type regional identity or cell class.

**Figure S08.**
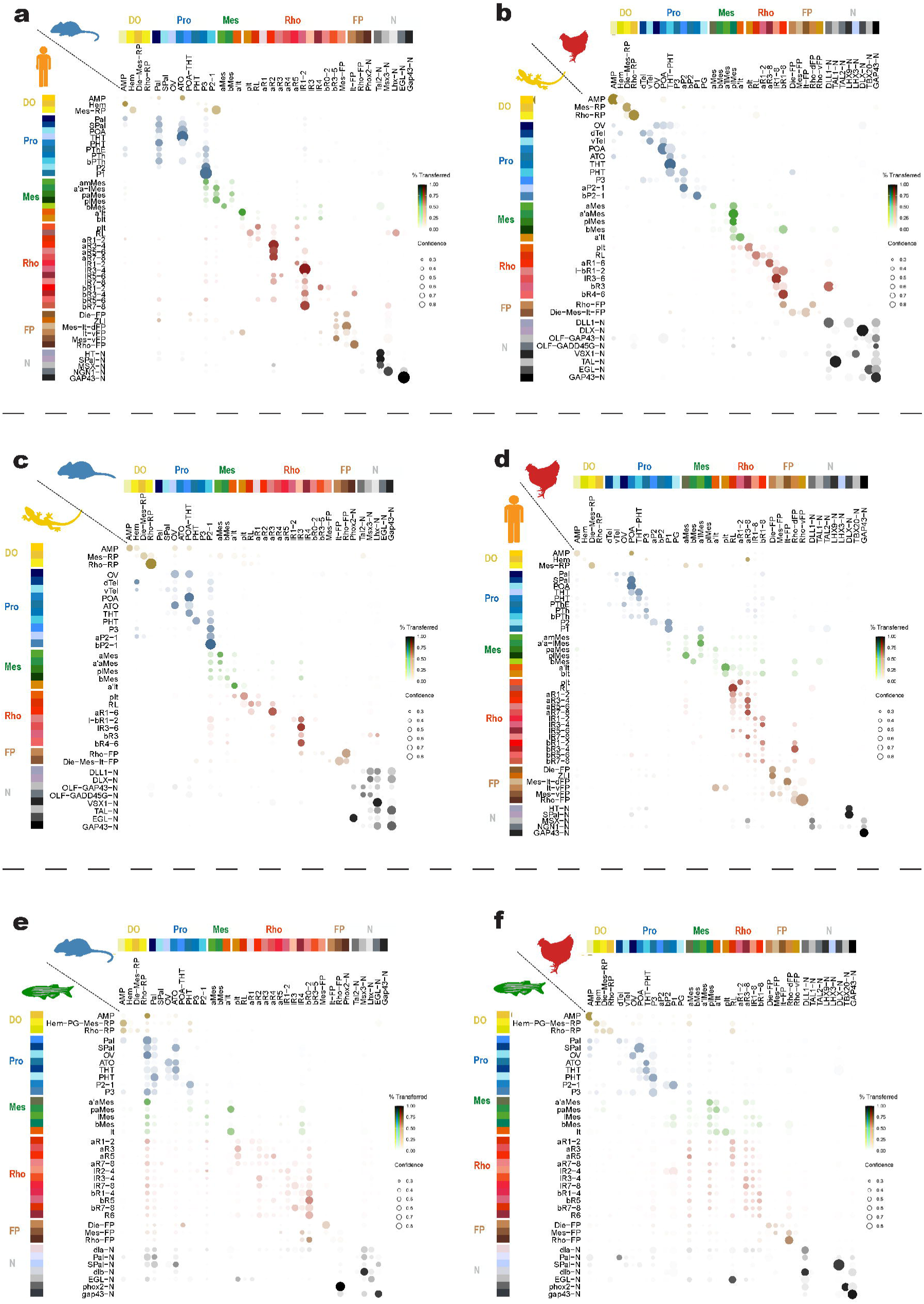
Cell type mapping: Label Transfer across vertebrate cell types. Pairwise comparisons between neural cells: mouse and human (**a**), chick and gecko (**b**); within amniotes: mouse and gecko (**c**), chick and human (**d**); or within vertebrates: mouse and zebrafish (**e**) and chick and zebrafish (**f**). Color intensity represents the percentage of cells transferred from query cell type to reference cell type. Circle size indicates the confidence score.

**Figure S09.**
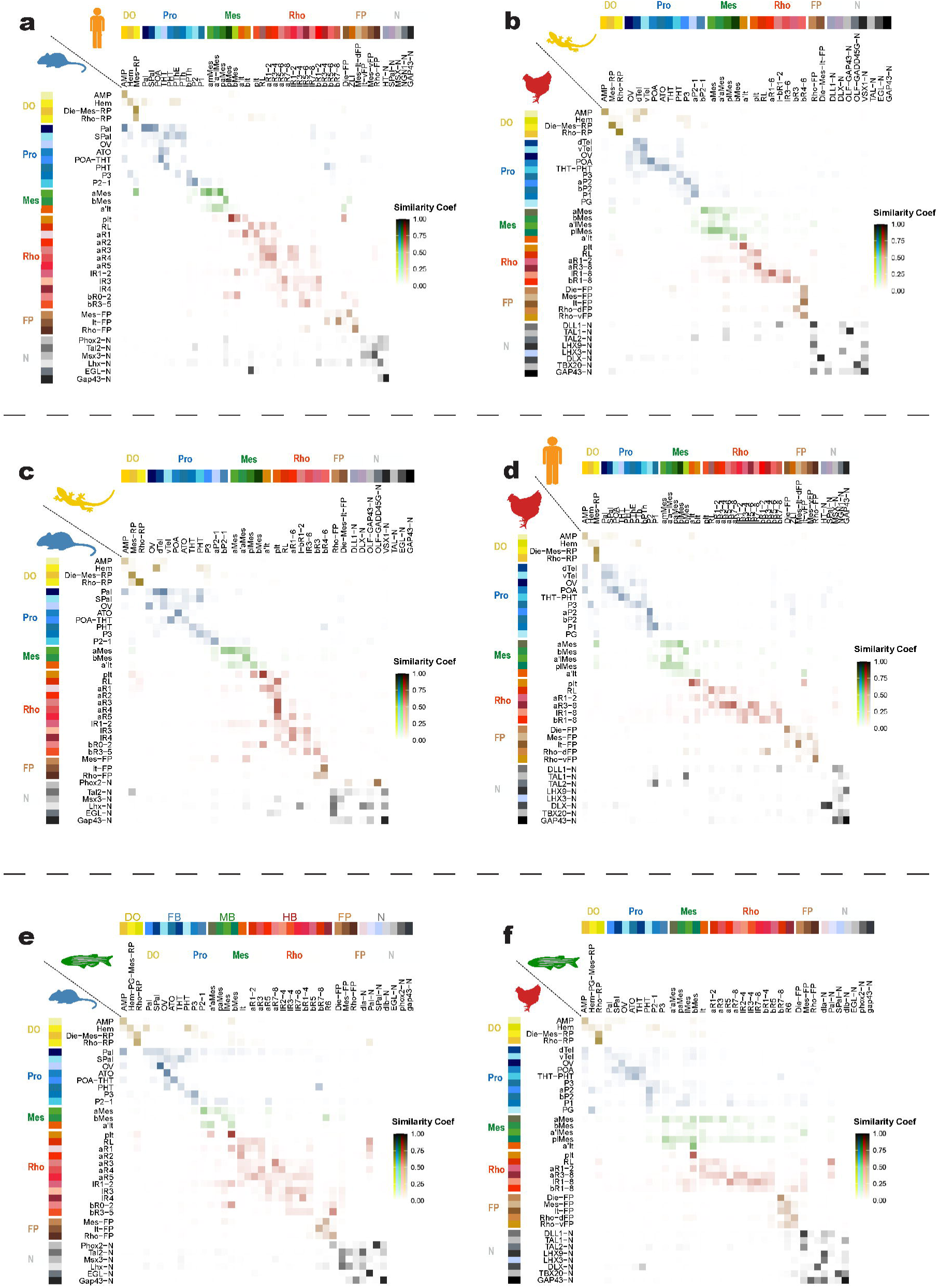
**Cell type mapping: SAMap across vertebrate cell types**. Pairwise comparisons between neural cells: mouse and human (**a**), chick and gecko (**b**); within amniotes: mouse and gecko (**c**), chick and human (**d**); or within vertebrates: mouse and zebrafish (**e**) and chick and zebrafish (**f**). Color intensity represents the similarity coefficient given by SAMap, fixed to 0 to 1 for comparative purposes.

**Figure S10.**
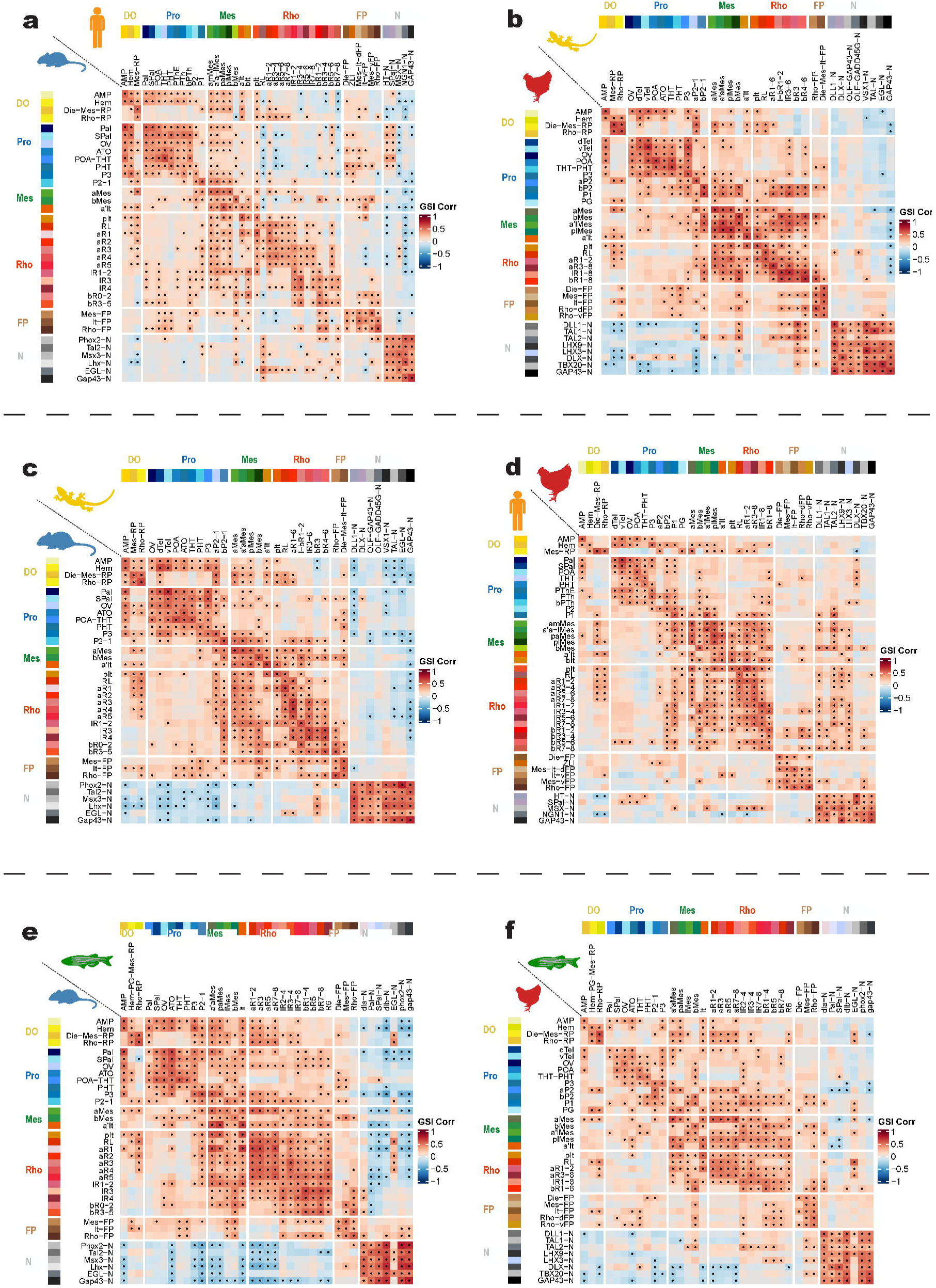
**Cell type similarity: GSI correlation across vertebrate cell types**. Pairwise correlations between neural cells: mouse and human (**a**), chick and gecko (**b**); within amniotes: mouse and gecko (**c**), chick and human (**d**); or within vertebrates: mouse and zebrafish (**e**) and chick and zebrafish (**f**). Colour scale is fixed to -1 to 1 for comparative purposes. Dots indicate statistical significance based on permutation tests.

**Figure S11.**
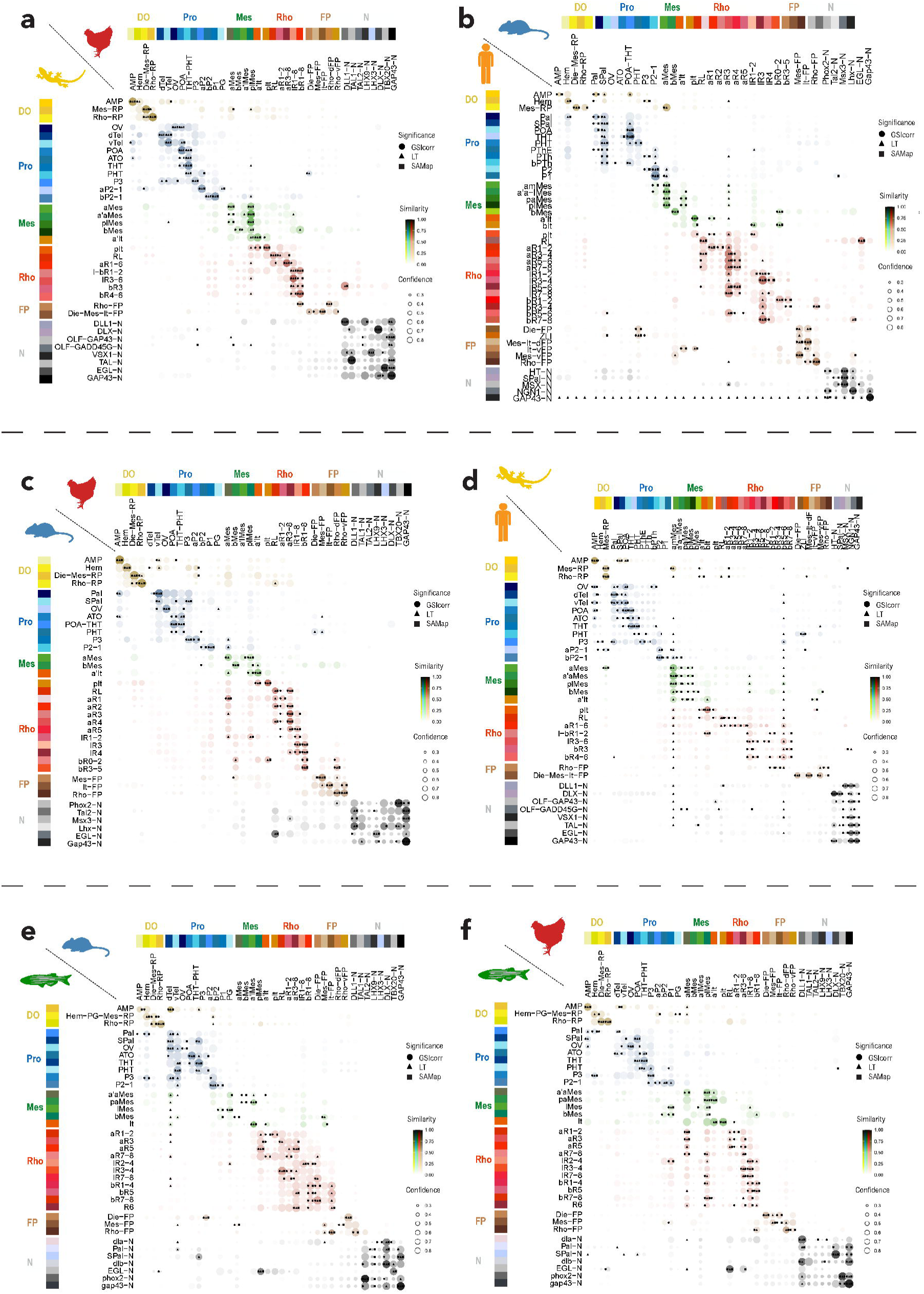
Multi-method similarities: Bassi plots across vertebrate cell types. Pairwise correlations between neural cells: mouse and human (**a**), chick and gecko (**b**); within amniotes: mouse and gecko (**c**), chick and human (**d**); or within vertebrates: mouse and zebrafish (**e**) and chick and zebrafish (**f**). Color intensity is the mean of the label transference percentage, the SAMap similaritiy and the correlation score minus the median overall correlation. Circle size indicates the confidence score based on the sd of the before mentioned three parameters. When the comparison value is on the 95^th^ percentile of its row, significance marks are indicated inside each comparison: circle (GSI Correlation), triangle (Label transference) and square (SAMap).

**Figure S12.**
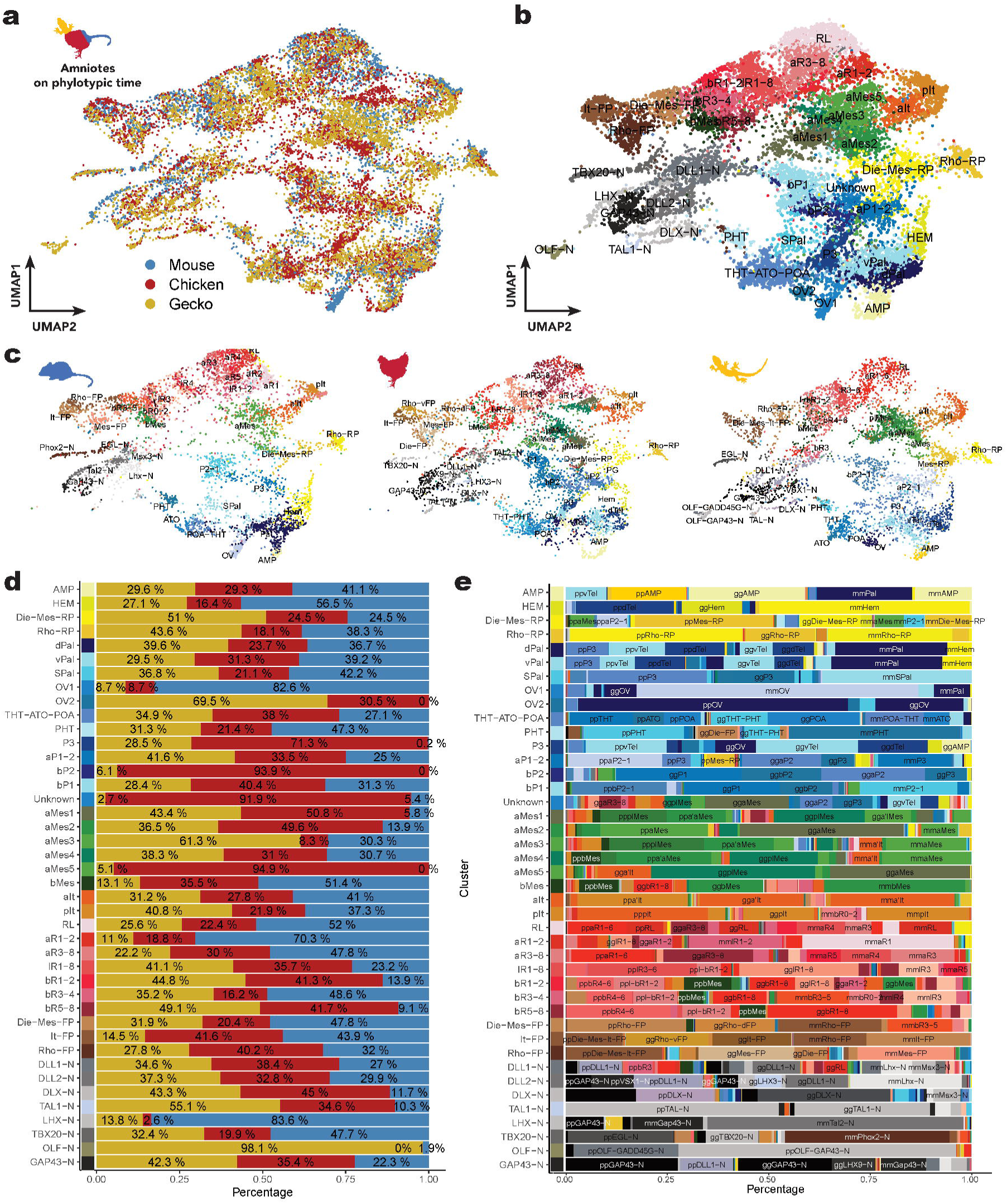
“RPCA” integration across amniote neural cell types: mouse, chick, gecko and zebrafish. a, UMAP of the integrated single cell atlas split by original species (mouse, blue; chick, red; gecko, yellow). **b,** Newly annotated integrated single cell atlas (resolution 2.5) using gene markers and original individual-species annotation. **c,** Subset of individual species cells maintaining original UMAP values. **d,** Proportion of cells from the three species in each newly clustered cell type. **e**, Proportion of cells from the original cell types (preceded by species initials) in each newly clustered cell type and color is the original in each individual species UMAP.

**Figure S13.**
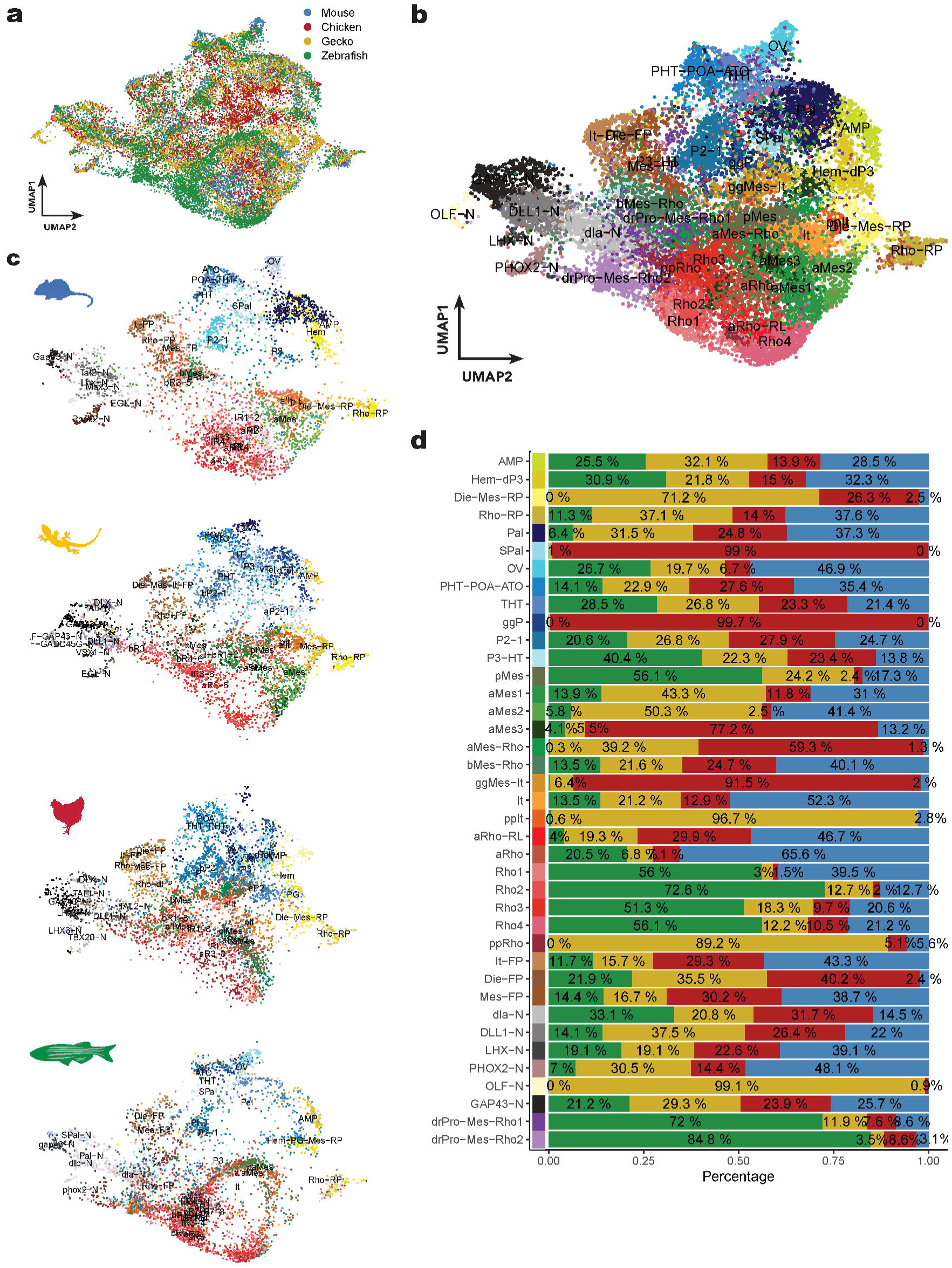
“RPCA” integration across vertebrate neural cell types: mouse, chick, gecko and zebrafish. a, UMAP of the integrated single cell atlas split by original species (mouse, blue; chick, red; gecko, yellow; zebrafish, green). **b,** Newly annotated integrated single cell atlas (resolution 2.5) using gene markers and original individual-species annotation (In purple, zebrafish specific and not neuroanatomical relevant; all following before mentioned color guides). **c,** Subset of individual species cells maintaining original UMAP values. **d,** Proportion of cells from the three species in each newly clustered cell type.

**Figure S14.**
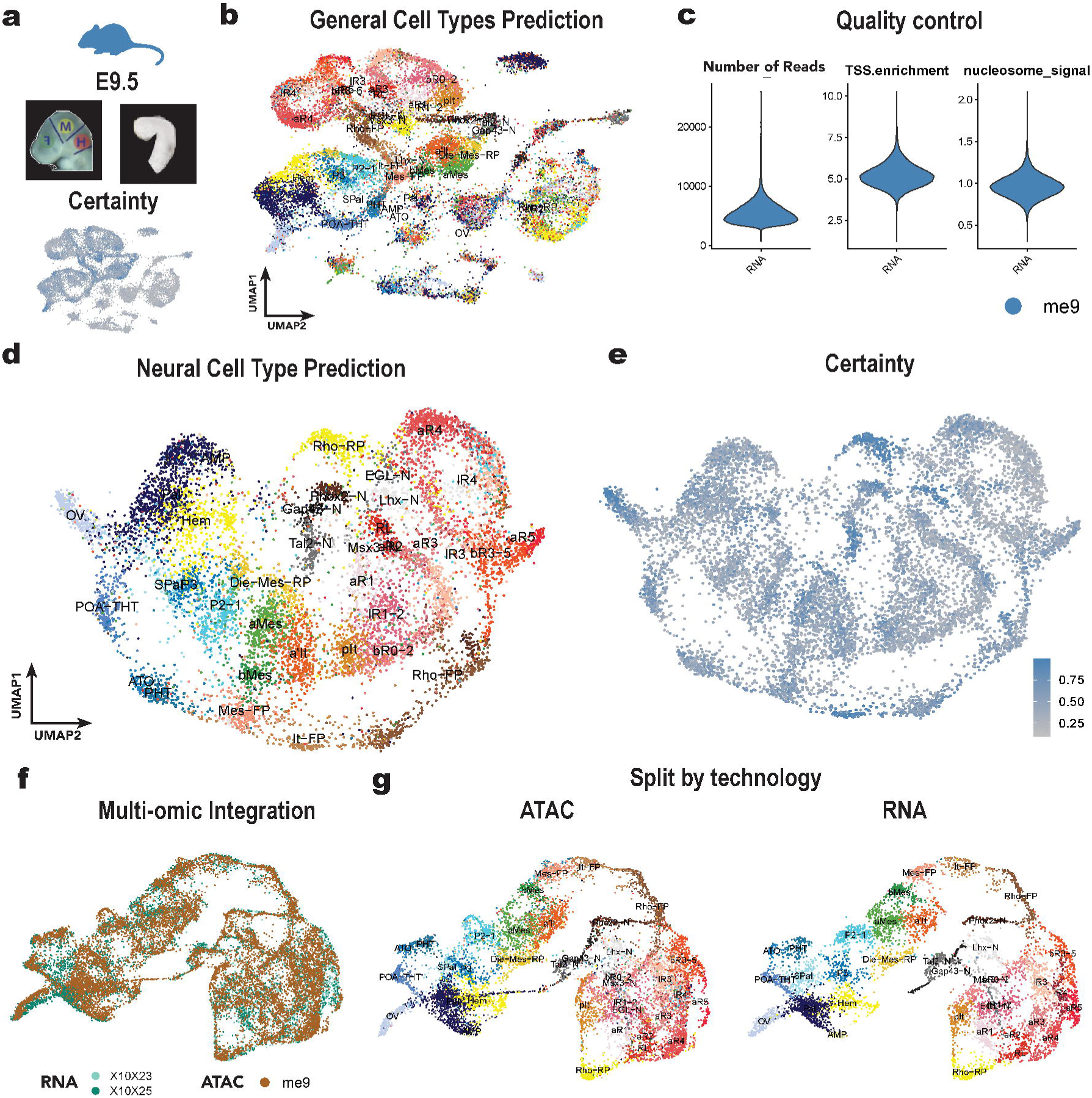
Quality control of single nuclei mouse atlas. **a**, Head morphology before^40^ and after brain dissection brain (*in house*) of E9.5 (TS11) mouse. **b**, UMAP plot of unfiltered predicted cell types. **c**, QC parameters of snATAC-seq replicate: number of reads mean per cell, TSS enrichment and nucleosome signal. **d**, Certainty of cell type prediction of unfiltered atlas. e, Neural Cell Type Prediction, final annotation after non-certain clusters. f, Certainty of final prediction. g, Multi-omic integration of scRNA-seq and snATAC-seq colored per replicate. h, Split of overlapping cell and nuclei from different technologies colored by cell types.

**Figure S15.**
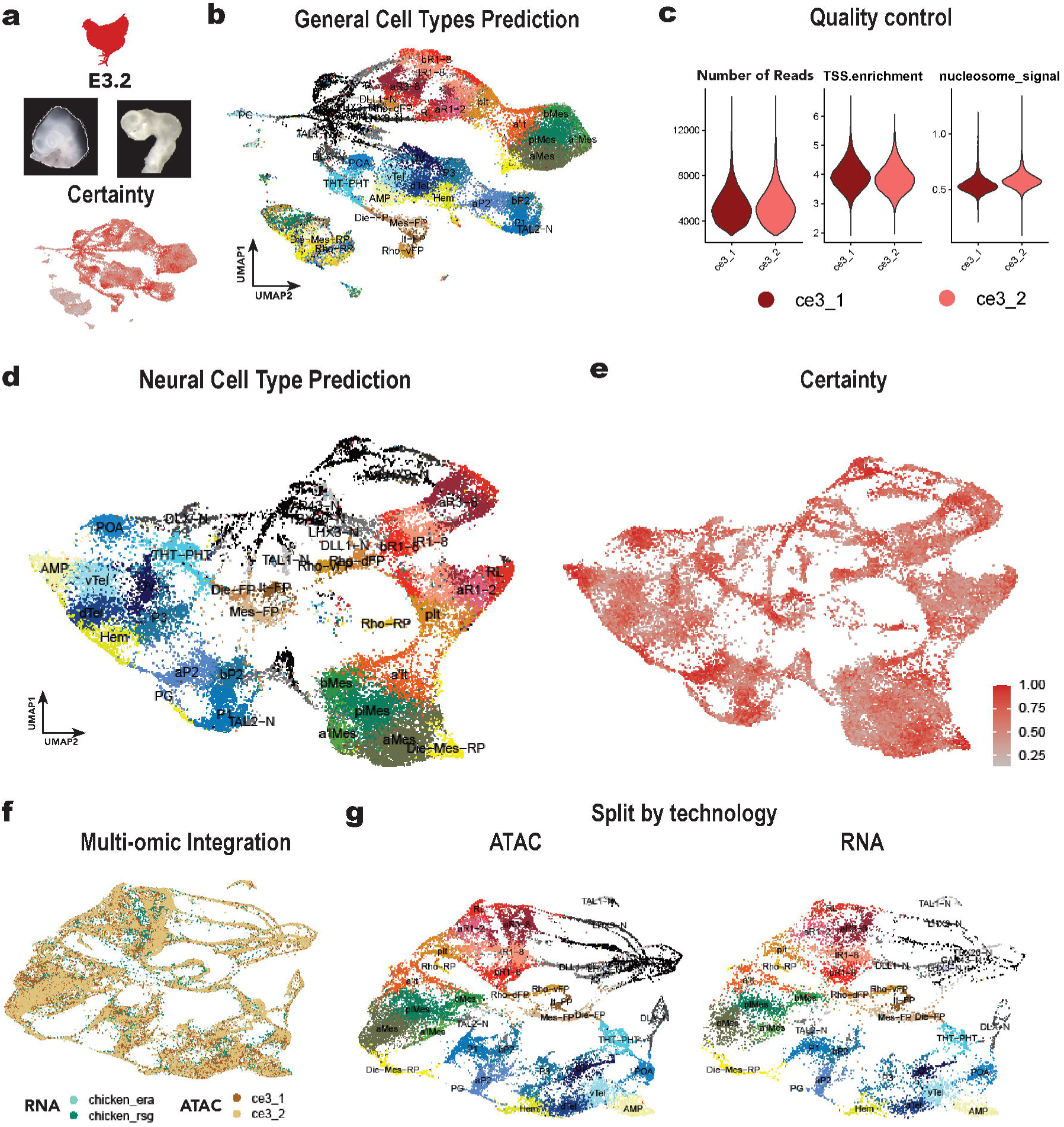
Quality control of single nuclei chick atlas. **a**, Head morphology before and after brain dissection (*in house*) of E3.2 chick. **b**, UMAP plot of unfiltered predicted cell types. **c**, QC parameters of snATAC-seq replicate: number of reads mean per cell, TSS enrichment and nucleosome signal. **d**, Certainty of cell type prediction of unfiltered atlas. e, Neural Cell Type Prediction, final annotation after non-certain clusters. f, Certainty of final prediction. g, Multi-omic integration of scRNA-seq and snATAC-seq colored per replicate. h, Split of overlapping cell and nuclei from different technologies colored by cell types.

**Figure S16.**
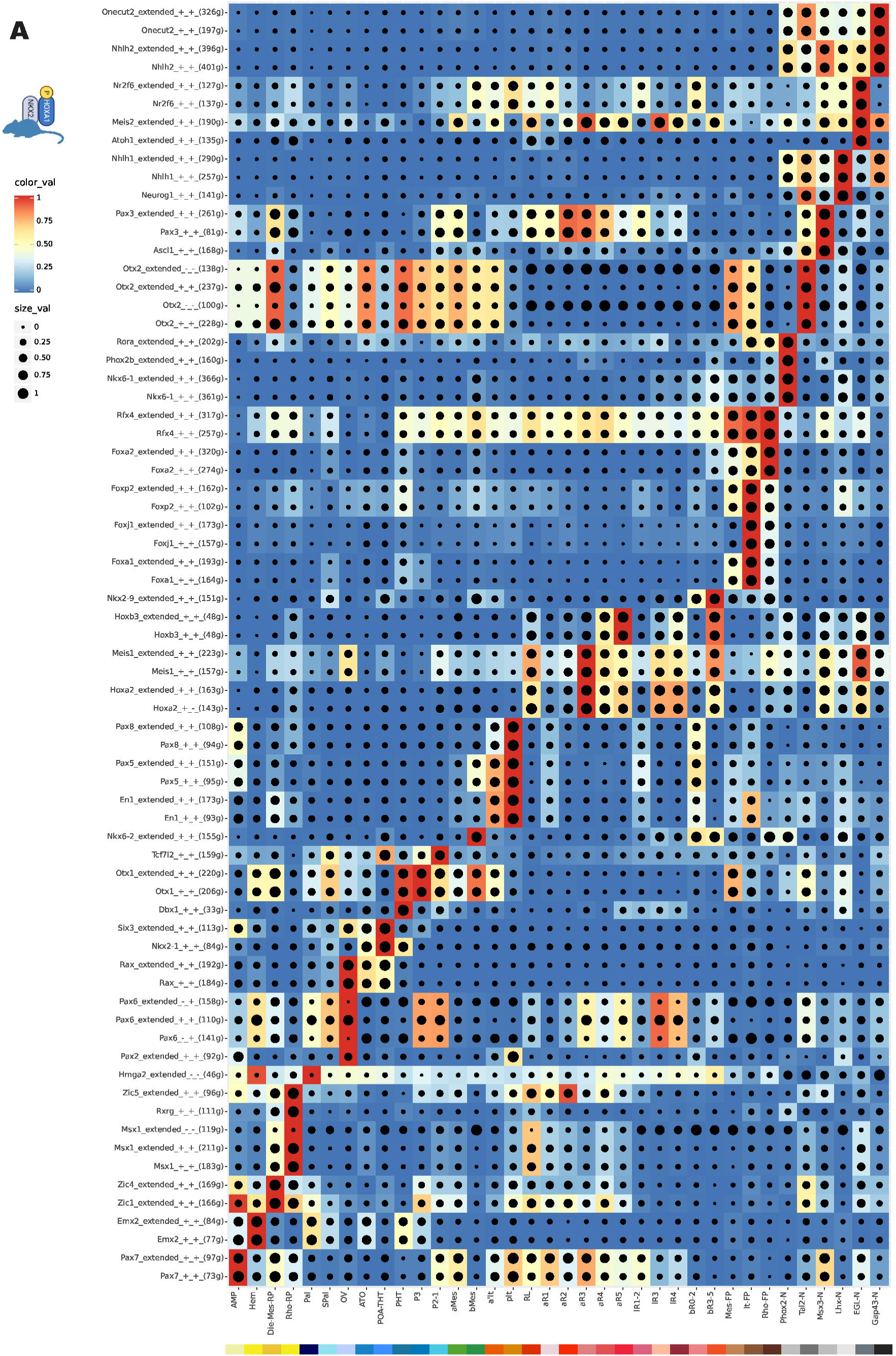
**Mouse early brain drivers: SCENIC+ eRegulons importance per cell type**. eRegulons (Y axis) are ordered by clustering to accommodate the importance for each cell type (X axis), which are ordered and color by region and type. Within the eRegulon lists, negative correlations are indicated by +/-folling computed RNA/ATAC correlations. Color intensity relates to eRegulon importance (AUC). Dot size indicates the size effect, or percentage of cells with the represented eRegulon importance.

**Figure S17.**
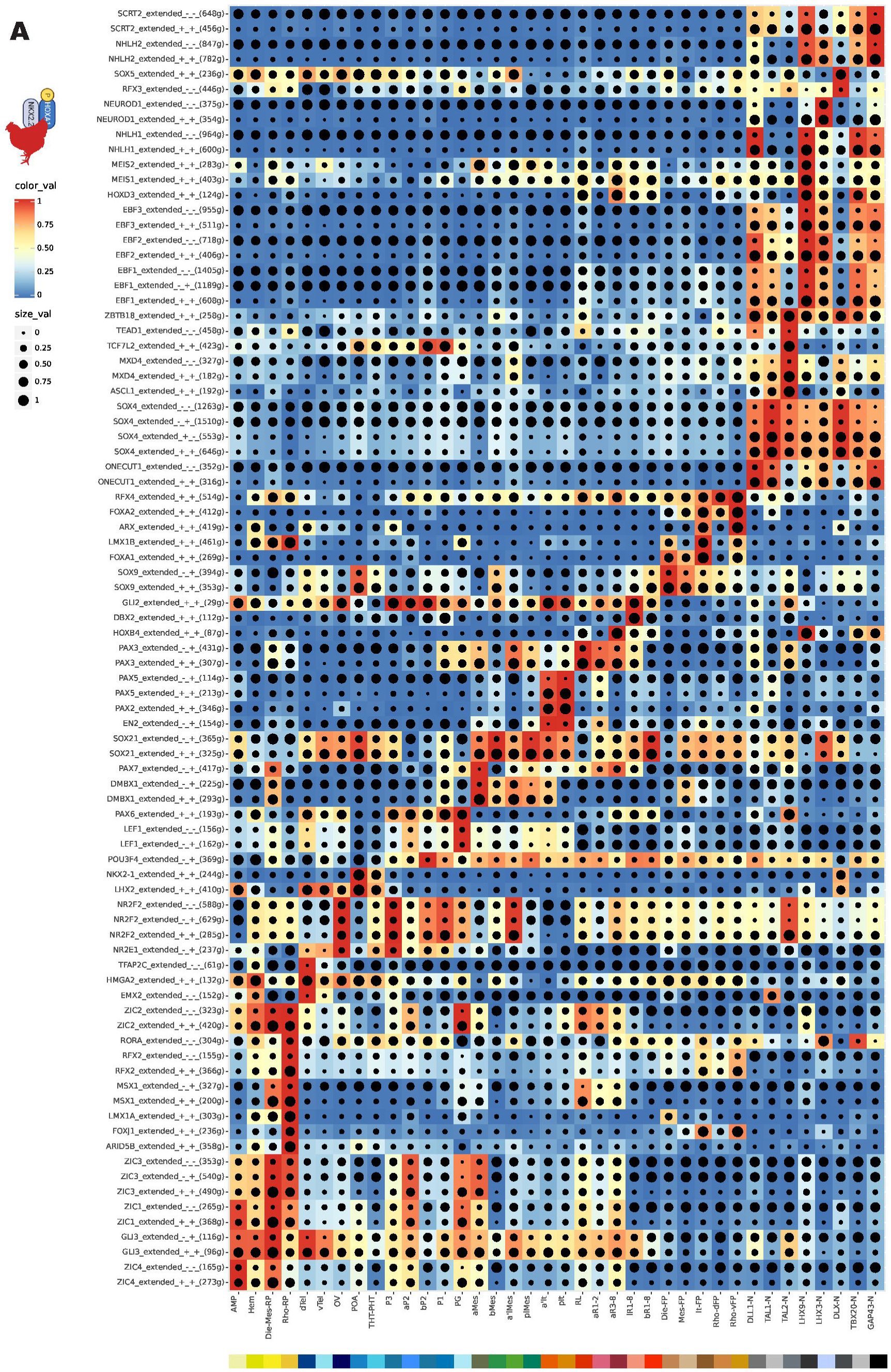
**Chick early brain drivers: SCENIC+ eRegulons importance per cell type**. eRegulons (Y axis) are ordered by clustering to accommodate the importance for each cell type (X axis), which are ordered and color by region and type. Within the eRegulon lists, negative correlations are indicated by +/-folling computed RNA/ATAC correlations. Color intensity relates to eRegulon importance (AUC). Dot size indicates the size effect, or percentage of cells with the represented eRegulon importance.

**Figure S18.**
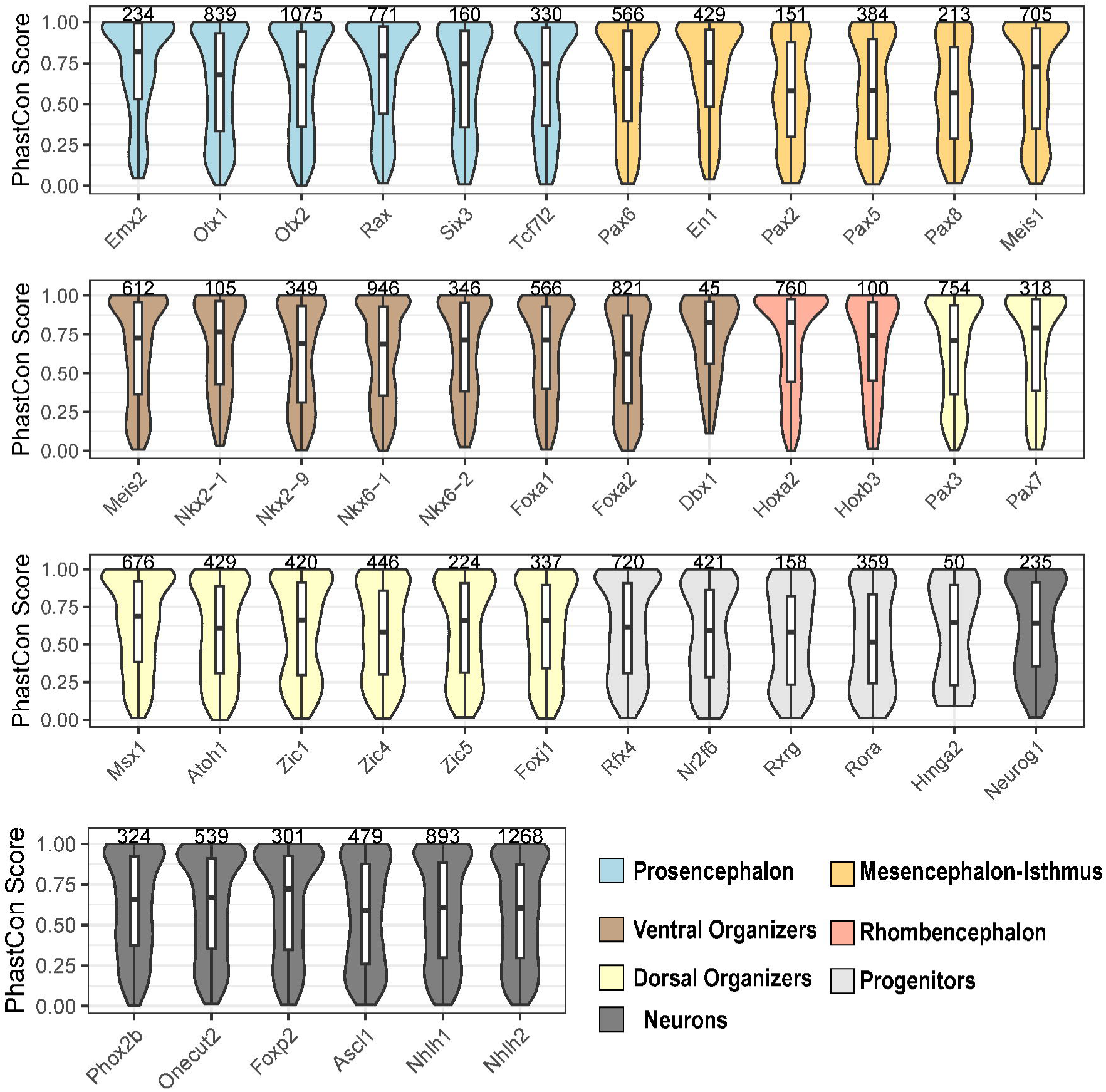
PhastCons on mouse SCENIC+ eRegulons. Mean conservation of region-phastCons scores of 100 bp windows across TFs eRegulons. Color code follows the cell type where TFs are mainly activated: prosencephalon (blue), mesencephalon-isthmus (orange), ventral organizer (brown), rhomencephalon (red), dorsal organizers (yellow), progenitors (light grey), neurons (dark grey).

**Figure S19.**
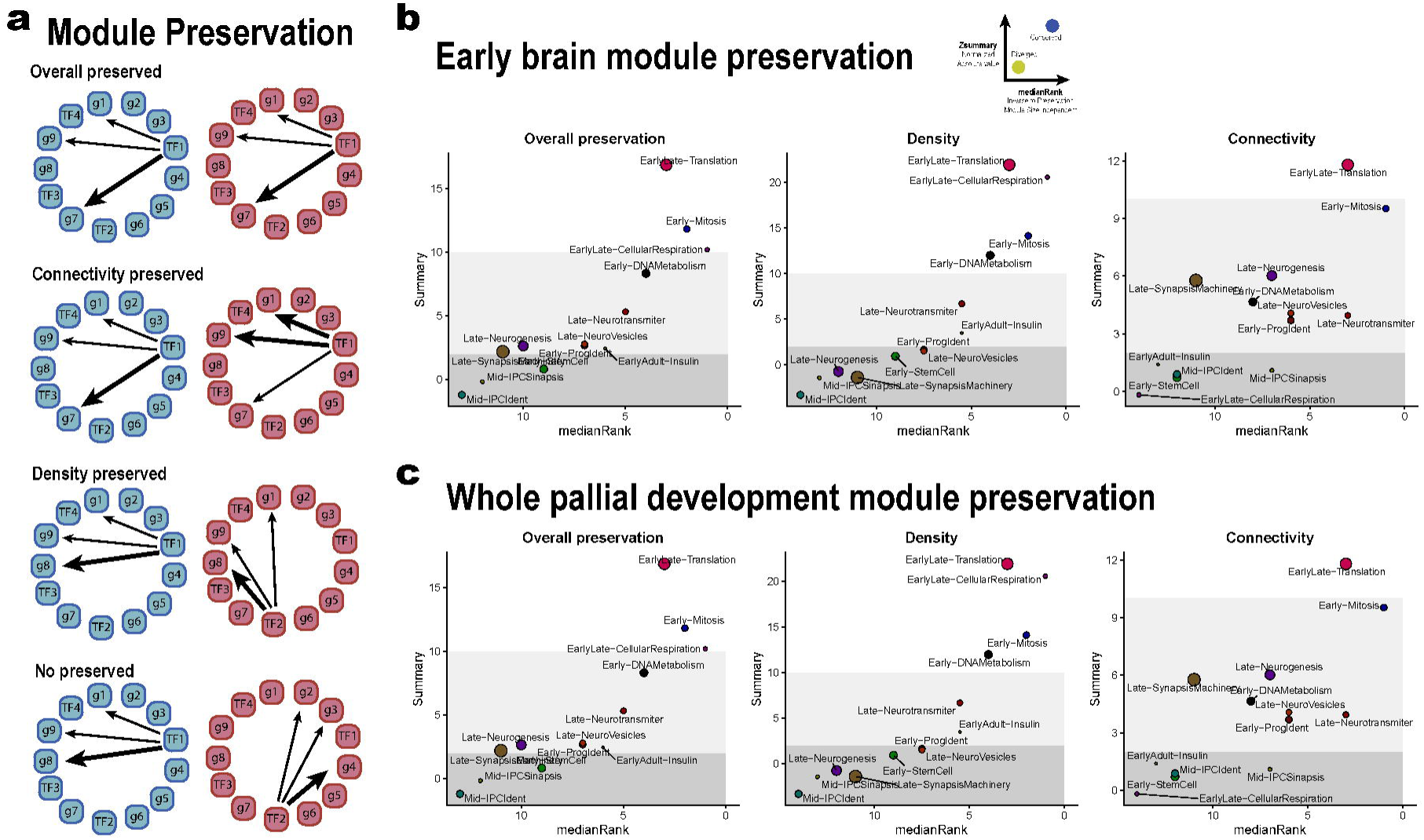
Module preservation by graph-theory statistics. **a**, Explanatory cartoon of the different evolutionary differences between networks (overall, connectivity and density). **b-c**, Early brain (**b**) and whole pallium development (**c**) module preservation at the overall, connectivity and density regarding Zsummary and medainRank (module size dependent and independent).

**Figure S20.**
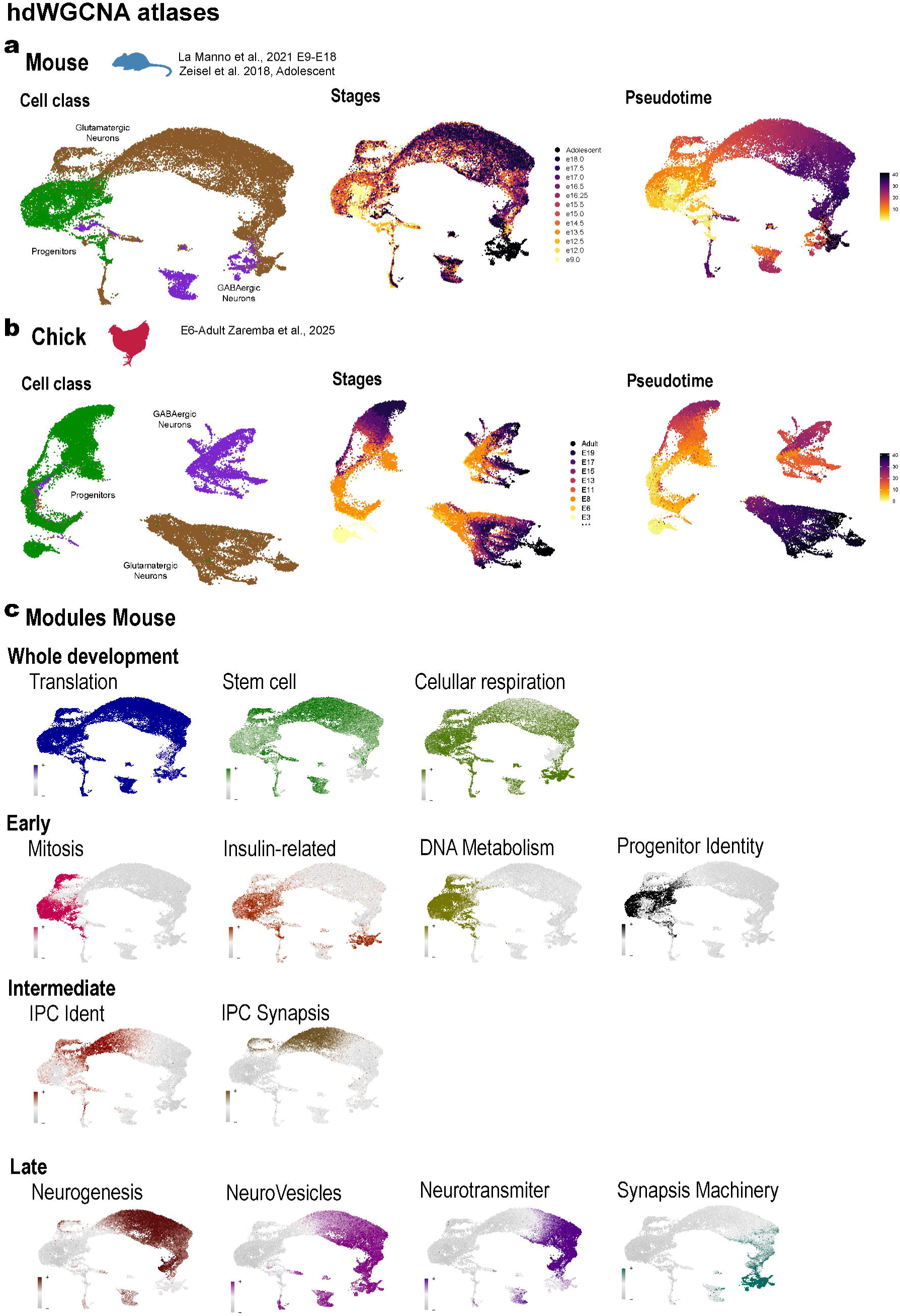
**Developmental atlases and modules of mouse and chick pallium**^40,64,88^. **a**, Single cell integrated atlas of developmental pallium mouse coloured by cell class (green, neural progenitors; brown, glutamatergic lineage; purple, gabaergic lineage), stages (early yellow, intermediate orange-red, later purple-black); pseudotime (monocle3 translation of cell developmental trajectories). **b**, Single cell integrated atlas of developmental pallium mouse coloured by cell class (green, neural progenitors; brown, glutamatergic lineage; purple, gabaergic lineage), stages (early yellow, intermediate orange-red, later purple-black); pseudotime (monocle3 translation of cell developmental trajectories). **c**, Module mean expression across mouse single cell developmental atlas grouped by temporal dynamics (whole-development, early, intermediate and late).

**Figure S21.**
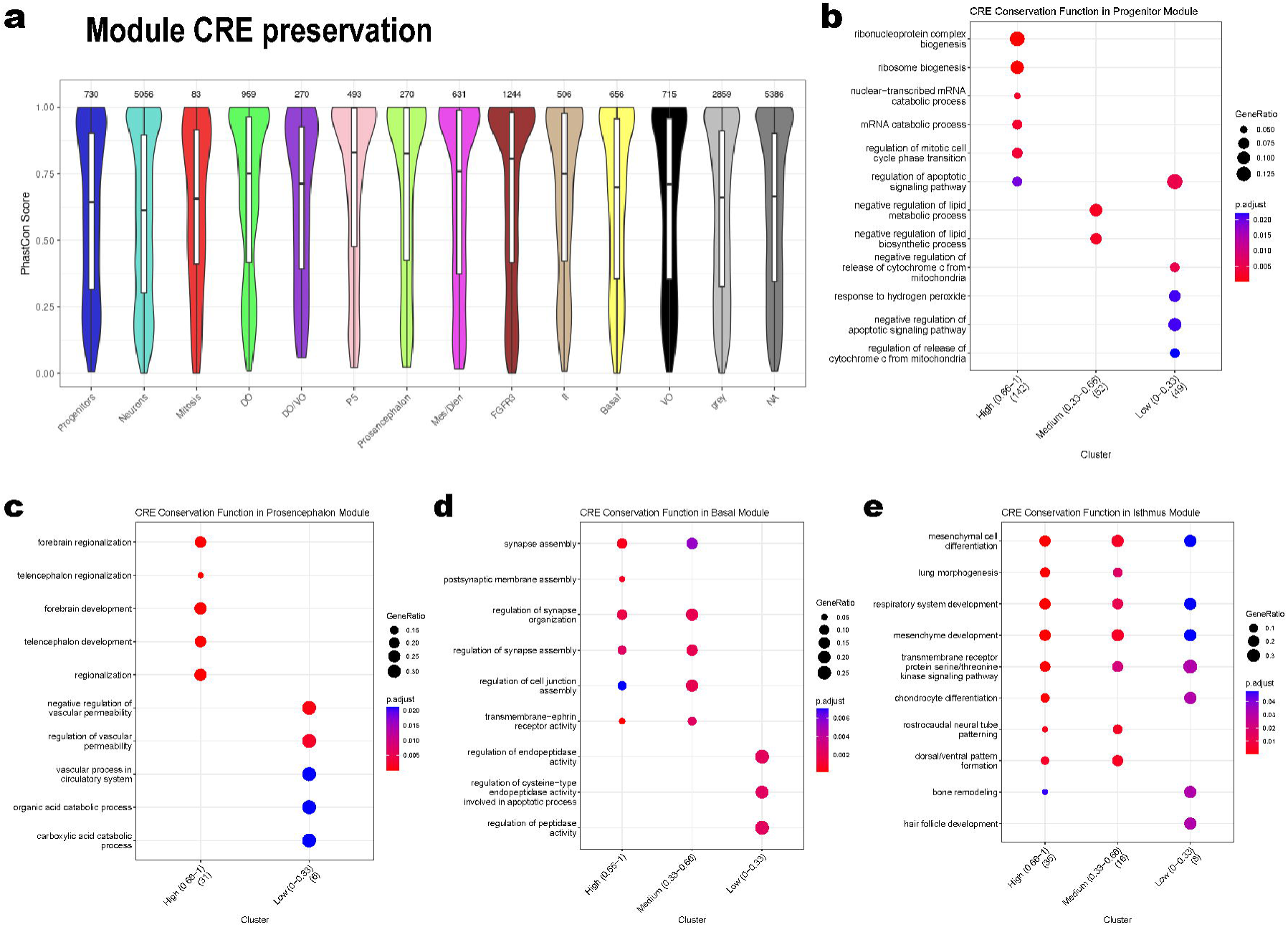
Module CREs preservation and functional enrichment. **a**, Mean conservation of region-phastCons scores of 100 bp windows across associated-gene-modules. Color code follows module colors. **b-e**, Functional enrichment of high (0.66-1), medium (0.33-0.66) and low (0-0.33) phastCons gene-associated score for several modules: Progenitors (**b**), Prosencephalon (**c**), Basal (**d**) and Isthmus (**e**).

**Figure S22.**
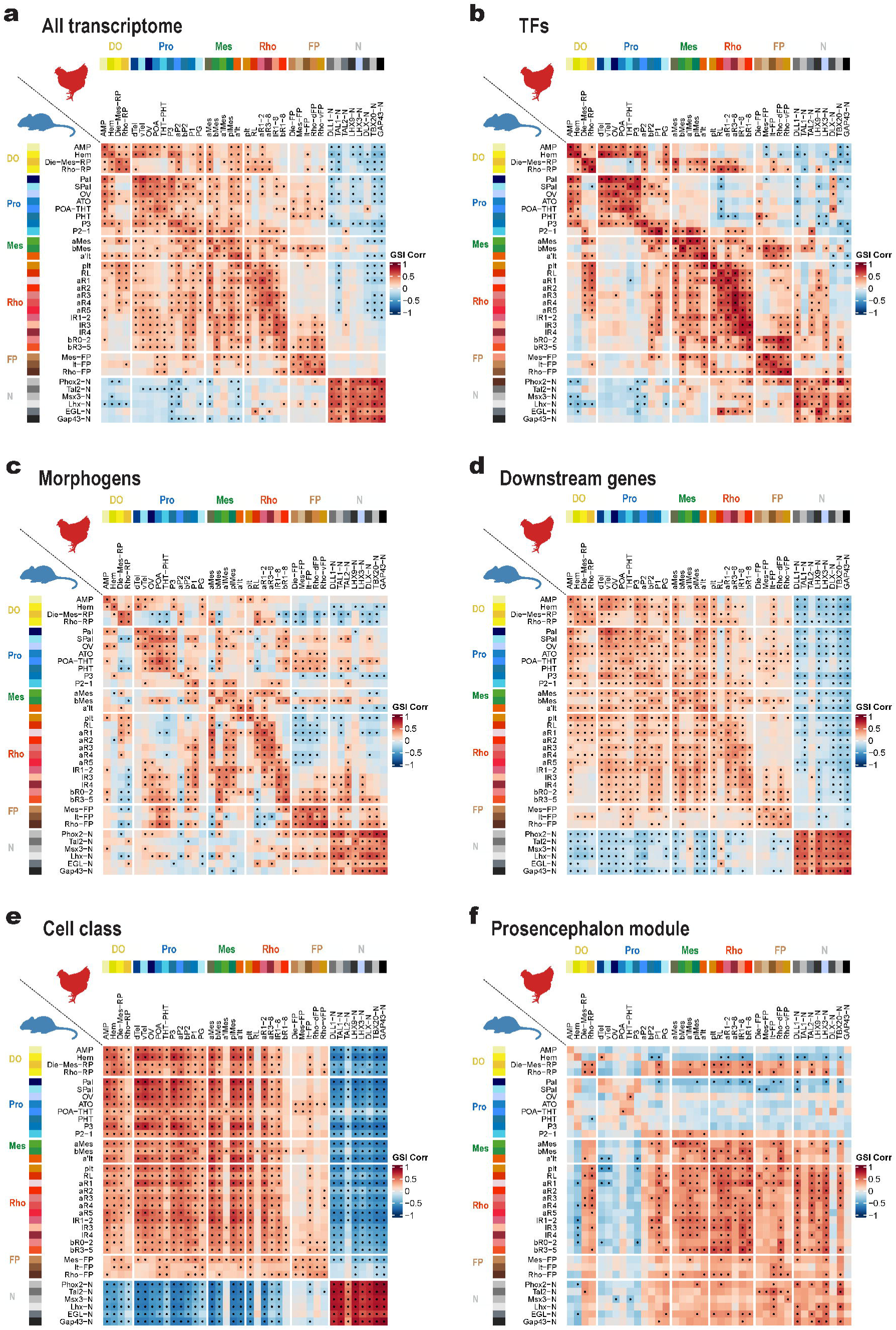
Filtered GSI Correlation per gene sets between mouse and chick. Filtered by none (**a**), TFs (**b**), morphogens (**c**), downstream genes (**d**), differentially expressed per cell class (**e**, progenitors vs neurons) and prosencephalon module (**f**).

**Figures S23.**
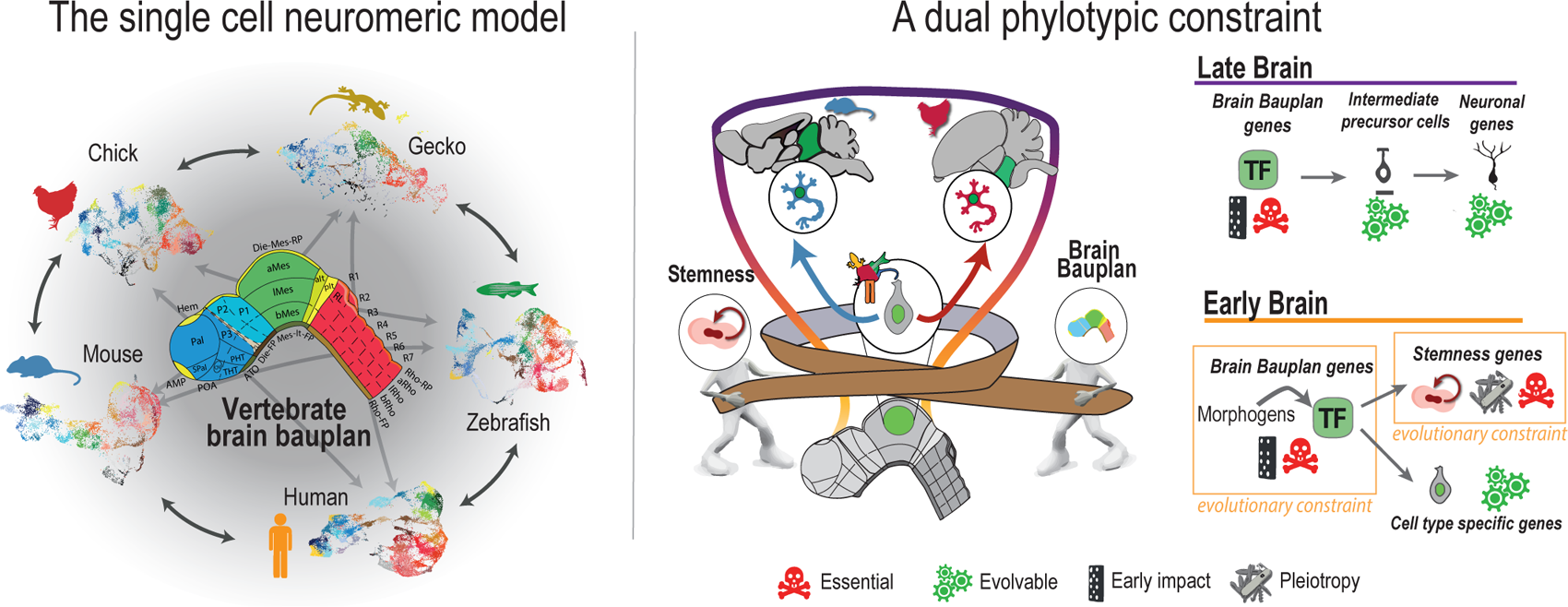
**– Graphical summary.**

## Supplementary Table

External database S1. Single cell dictionary: cell types and marker genes. External database S2. eRegulon metadata for mouse and chick early brain.

## Notes

https://github.com/rodrisenovilla/Senovilla-Ganzo2025

### Competing Interest Statement

The authors have declared no competing interest.

## References

1. Woodger JH. On biological transformations in Growth and form: essays presented to D’Arcy Thompson. (ed. Le Gros Clark WE, M. PB.) (Oxford : Clarendon Press., 1945).

2. Niklas, K. J., Cobb, E. D. & Kutschera, U. Haeckel’s Biogenetic Law and the Land Plant Phylotypic Stage. Bioscience 66, 510–519 (2016).

3. Haeckel, E. H. P. A. Generelle morphologie der organismen. Allgemeine grundzüge der organischen formen-wissenschaft, mechanisch begründet durch die von Charles Darwin reformirte descendenztheorie . Berlin, G. Reimer https://archive.org/details/generellemorphol01haec/mode/2up (1866).

4. Abzhanov, A. von Baer’s law for the ages: lost and found principles of developmental evolution. Trends Genet 29, 712–722 (2013).

5. Baer, K. E. von. *Über Entwickelungsgeschichte Der Thiere. Beobachtung Und Reflexion*. *Über Entwickelungsgeschichte der Thiere*. *Beobachtung und Reflexion* (Bei den Gebrüdern Bornträger, Königsberg, 1828). doi:10.5962/bhl.title.6303.

6. Puelles, L., Harrison, M., Paxinos, G. & Watson, C. A developmental ontology for the mammalian brain based on the prosomeric model. Trends Neurosci 36, 570–578 (2013).

7. Puelles, L. & Ferran, J. L. Concept of neural genoarchitecture and its genomic fundament. Front Neuroanat 6, 1–8 (2012).

8. His, W. Unsere Körperform Und Das Physiologische Problem Ihrer Entstehung: Briefe an Einen Befreundeten Naturforscher. (FCW Vogel, 1874).

9. Bergquist, H. STUDIES ON THE CEREBRAL TUBE IN VERTEBRATES. THE NEUROMERES. Acta Zoologica 33, 117–187 (1952).

10. García-Moreno, F. & Molnár, Z. Variations of telencephalic development that paved the way for neocortical evolution. Prog Neurobiol 101865 (2020) doi:10.1016/j.pneurobio.2020.101865.

11. Rueda-Alaña, E. et al. Evolutionary convergence of sensory circuits in the pallium of amniotes. Science (1979) 387, (2025).

12. Kappers, C. U. A. & Huber, G. C. The Comparative Anatomy of the Nervous System of Vertebrates, Including Man . (Macmillan, 1936).

13. Puelles, L., Martinez-de-la-Torre, M., Bardet, S. & Rubenstein, J. L. R. Hypothalamus. The Mouse Nervous System 221–312 (2012) doi:10.1016/B978-0-12-369497-3.10008-1.

14. Graham, A., Papalopulu, N. & Krumlauf, R. The murine and Drosophila homeobox gene complexes have common features of organization and expression. Cell 57, 367–378 (1989).

15. Duboule, D. & Dolle, P. The structural and functional organization of the murine HOX gene family resembles that of Drosophila homeotic genes. EMBO J 8, 1497 (1989).

16. Raff, R. A. The shape of life : genes, development, and the evolution of animal form. 520 (1996).

17. Duboule, D. Temporal colinearity and the phylotypic progression: A basis for the stability of a vertebrate Bauplan and the evolution of morphologies through heterochrony. Development 120, 135–142 (1994).

18. Slack, J. M., Holland, P. W. & Graham, C. F. The zootype and the phylotypic stage. Nature 361, 490–492 (1993).

19. Slack, J. Developmental biology: A Rosetta stone for pattern formation in animals? Nature 1984 310:5976 310, 364–365 (1984).

20. Irie, N. & Kuratani, S. Comparative transcriptome analysis reveals vertebrate phylotypic period during organogenesis. Nat Commun 2, (2011).

21. Cardoso-Moreira, M. et al. Gene expression across mammalian organ development. Nature 571, 505–509 (2019).

22. 22. Hu, H., et al. Constrained vertebrate evolution by pleiotropic genes. *Nat Ecol Evol* 1, 1722– 1730 (2017).

23. Zhang, J. & Wagner, G. P. On the definition and measurement of pleiotropy. Trends in Genetics 29, 383–384 (2013).

24. Hansen, T. F. Is modularity necessary for evolvability? Remarks on the relationship between pleiotropy and evolvability. BioSystems 69, 83–94 (2003).

25. Arendt, D. et al. The origin and evolution of cell types. Nature Reviews Genetics vol. 17 744–757 Preprint at 10.1038/nrg.2016.127 (2016).

26. Parker, J. & Pennell, M. The cellular substrate of evolutionary novelty. Current Biology 35, R626–R637 (2025).

27. Piasecka, B., Lichocki, P., Moretti, S., Bergmann, S. & Robinson-Rechavi, M. The Hourglass and the Early Conservation Models-Co-Existing Patterns of Developmental Constraints in Vertebrates. PLoS Genet 9, (2013).

28. Richardson, M. K. Heterochrony and the phylotypic period. Dev Biol **172**, 412–421 (1995).

29. Sarropoulos, I. et al. Developmental and evolutionary dynamics of cis-regulatory elements in mouse cerebellar cells. Science (1979) eabg4696 (2021) doi:10.1126/SCIENCE.ABG4696.

30. Owen, R. On the Archetype and Homologies of the Vertebrate Skeleton. (author, 1848).

31. Wagner, G. P. The biological homology concept. Annual review of ecology and systematics*. Vol.* 20 **20**, 51–69 (1989).

32. Wagner, G. P. The developmental genetics of homology. Nat Rev Genet 8, 473–479 (2007).

33. Schindler, M. et al. Comparative single-cell analyses reveal evolutionary repurposing of a conserved gene programme in bat wing development. Nature Ecology & Evolution 2025 1– 17 (2025) doi:10.1038/s41559-025-02780-x.

34. Shubin, N., Tabin, C. & Carroll, S. Fossils, genes and the evolution of animal limbs. Nature 1997 388:6643 388, 639–648 (1997).

35. Briscoe, S. D. Field Homology: Still a Meaningless Concept. Brain Behav Evol 93, 1–3 (2019).

36. Briscoe, S. D. & Ragsdale, C. W. Homology, neocortex, and the evolution of developmental mechanisms. Science (1979) 362, 190–193 (2018).

37. Northcutt, R. G. Field homology: a meaningless concept. Eur J Morphol 37, 95–99 (1999).

38. Shubin, N., Tabin, C. & Carroll, S. Deep homology and the origins of evolutionary novelty. Nature 2009 457:7231 457, 818–823 (2009).

39. Love, A. C. Functional homology and homology of function: Biological concepts and philosophical consequences. Biol Philos 22, 691–708 (2007).

40. La Manno, G. et al. Molecular architecture of the developing mouse brain. Nature 2021 596:7870 596, 92–96 (2021).

41. Braun, E. et al. Comprehensive cell atlas of the first-trimester developing human brain. Science 382, eadf1226 (2023).

42. Raj, B. et al. Emergence of Neuronal Diversity during Vertebrate Brain Development. Neuron 108, 1058–1074.e6 (2020).

43. Rueda-Alaña, E. & García-Moreno, F. Time in Neurogenesis: Conservation of the Developmental Formation of the Cerebellar Circuitry. Brain Behav Evol (2022) doi:10.1159/000519068.

44. Guirado, S. & Carlos Dávila, J. Encyclopedia of Neuroscience Evolution of the Optic Tectum in Amniotes Structure of the Optic Tectum in Reptiles. Encyclopedia of Neuroscience 1–15 (2009).

45. Zhang, S., Xiangtao, L. I., Jiecong, L. I. N., Qiuzhen, L. I. N. & Wong, K. C. Review of single-cell RNA-seq data clustering for cell-type identification and characterization. RNA 29, 517 (2023).

46. Lähnemann, D. et al. Eleven grand challenges in single-cell data science. Genome Biology 2020 21:1 21, 1–35 (2020).

47. Krumlauf, R. & Wilkinson, D. G. Segmentation and patterning of the vertebrate hindbrain. Development 148, (2021).

48. Nakamura, H. Midbrain patterning: polarity formation of the tectum, midbrain regionalization, and isthmus organizer. *Patterning and Cell Type Specification in the Developing CNS and PNS: Comprehensive Developmental Neuroscience*, Second Edition 87–106 (2020) doi:10.1016/B978-0-12-814405-3.00005-9.

49. Herrick, C. J. The morphology of the forebrain in amphibia and reptilia. Journal of comparative Neurology and Psychology 20, 413–547 (1910).

50. Bedont, J. L., Newman, E. A. & Blackshaw, S. Patterning, specification, and differentiation in the developing hypothalamus. Wiley Interdiscip Rev Dev Biol 4, 445–468 (2015).

51. Lemaire, L. A., Cao, C., Yoon, P. H., Long, J. & Levine, M. The hypothalamus predates the origin of vertebrates. Sci Adv 7, (2021).

52. Senovilla-Ganzo, R. & García-Moreno, F. The Phylotypic Brain of Vertebrates, from Neural Tube Closure to Brain Diversification. Brain Behav Evol 99, 45–68 (2024).

53. Pani, A. M. et al. Ancient deuterostome origins of vertebrate brain signalling centres. Nature 483, 289–294 (2012).

54. Lagutin, O. V. et al. Six3 repression of Wnt signaling in the anterior neuroectoderm is essential for vertebrate forebrain development. Genes Dev 17, 368–379 (2003).

55. Tambalo, M., Mitter, R. & Wilkinson, D. G. A single cell transcriptome atlas of the developing zebrafish hindbrain. Development (Cambridge) 147, (2020).

56. Delile, J. et al. Single cell transcriptomics reveals spatial and temporal dynamics of gene expression in the developing mouse spinal cord. Development (Cambridge) 146, (2019).

57. Tosches, M. A. et al. Evolution of pallium, hippocampus, and cortical cell types revealed by single-cell transcriptomics in reptiles. Science (1979) 360, 881–888 (2018).

58. Colquitt, B. M., Merullo, D. P., Konopka, G., Roberts, T. F. & Brainard, M. S. Cellular transcriptomics reveals evolutionary identities of songbird vocal circuits. Science (1979) 371, (2021).

59. Stuart, T. et al. Comprehensive integration of single cell data. Cell 1888–1902 (2019) doi:10.1101/460147.

60. Tarashansky, A. J. et al. Mapping single-cell atlases throughout metazoa unravels cell type evolution. Elife 10, (2021).

61. Zaremba, B. et al. Developmental origins and evolution of pallial cell types and structures in birds. Science (1979) 387, (2025).

62. Hain, D. et al. Molecular diversity and evolution of neuron types in the amniote brain. Science 377, (2022).

63. Shafer, M. E. R. Cross-Species Analysis of Single-Cell Transcriptomic Data. Frontiers in Cell and Developmental Biology vol. 7 175 Preprint at 10.3389/fcell.2019.00175 (2019).

64. Song, Y., Miao, Z., Brazma, A. & Papatheodorou, I. Benchmarking strategies for cross-species integration of single-cell RNA sequencing data. Nature Communications 2023 14:1 14, 1–17 (2023).

65. Stuart, T., Srivastava, A., Madad, S., Lareau, C. A. & Satija, R. Single-cell chromatin state analysis with Signac. Nature Methods 2021 18:11 18, 1333–1341 (2021).

66. Camacho, C. et al. BLAST+: Architecture and applications. BMC Bioinformatics **10**, 1–9 (2009).

67. Shafer, M. E. R., Sawh, A. N. & Schier, A. F. Gene family evolution underlies cell-type diversification in the hypothalamus of teleosts. Nat Ecol Evol 6, 63–76 (2022).

68. Glasauer, S. M. K. & Neuhauss, S. C. F. Whole-genome duplication in teleost fishes and its evolutionary consequences. Mol Genet Genomics 289, 1045–1060 (2014).

69. Achim, K. et al. The role of Tal2 and Tal1 in the differentiation of midbrain GABAergic neuron precursors. Biol Open 2, 990–997 (2013).

70. Korsunsky, I. et al. Fast, sensitive and accurate integration of single-cell data with Harmony. Nat Methods 16, 1289–1296 (2019).

71. Kuemmerle, L. B. et al. Probe set selection for targeted spatial transcriptomics. Nature Methods 2024 21:12 21, 2260–2270 (2024).

72. Ke, R. et al. In situ sequencing for RNA analysis in preserved tissue and cells. Nature Methods 2013 10:9 10, 857–860 (2013).

73. Puelles, L. & Medina, L. Field homology as a way to reconcile genetic and developmental variability with adult homology. Brain Res Bull 57, 243–255 (2002).

74. Trevers, K. E. et al. A gene regulatory network for neural induction. Elife 12, (2023).

75. Levine, M. & Davidson, E. H. Gene regulatory networks for development. Proc Natl Acad Sci U S A 102, 4936–4942 (2005).

76. Bravo González-Blas, C., et al. SCENIC+: single-cell multiomic inference of enhancers and gene regulatory networks. Nature Methods 2023 20:9 20, 1355–1367 (2023).

77. Siepel, A. et al. Evolutionarily conserved elements in vertebrate, insect, worm, and yeast genomes. Genome Res 15, 1034–1050 (2005).

78. Phan, M. H. Q. et al. Conservation of regulatory elements with highly diverged sequences across large evolutionary distances. Nat Genet 57, 1524–1534 (2025).

79. Hecker, N. et al. Enhancer-driven cell type comparison reveals similarities between the mammalian and bird pallium. Science (1979) 387, (2025).

80. Langfelder, P., Luo, R., Oldham, M. C. & Horvath, S. Is My Network Module Preserved and Reproducible? PLoS Comput Biol 7, e1001057 (2011).

81. Zeisel, A. et al. Molecular Architecture of the Mouse Nervous System. Cell 174, 999–1014.e22 (2018).

82. Irie, N. & Kuratani, S. The developmental hourglass model: a predictor of the basic body plan? Development 141, 4649–4655 (2014).

83. Groza, T. et al. The International Mouse Phenotyping Consortium: comprehensive knockout phenotyping underpinning the study of human disease. Nucleic Acids Res 51, D1038–D1045 (2023).

84. Puelles, L. Survey of midbrain, diencephalon, and hypothalamus neuroanatomic terms whose prosomeric definition conflicts with columnar tradition. Front Neuroanat 13, 1–33 (2019).

85. Albuixech-Crespo, B. et al. Molecular regionalization of the developing amphioxus neural tube challenges major partitions of the vertebrate brain. PLoS Biol 15, (2017).

86. Duboule, D. & Wilkins, A. S. The evolution of ‘bricolage’. Trends in Genetics 14, 54–59 (1998).

87. Koonin, E. V., Krupovic, M., Ishino, S. & Ishino, Y. The replication machinery of LUCA: common origin of DNA replication and transcription. BMC Biol 18, 61 (2020).

88. Lin, H. C. et al. Human neuron subtype programming via single-cell transcriptome-coupled patterning screens. Science (1979) 389, (2025).

89. Galis, F. & Metz, J. A. J. Testing the vulnerability of the phylotypic stage: On modularity and evolutionary conservation. Journal of Experimental Zoology 291, 195–204 (2001).

90. Rozenblatt-Rosen, O., Stubbington, M. J. T., Regev, A. & Teichmann, S. A. The Human Cell Atlas: from vision to reality. Nature 2017 550:7677 550, 451–453 (2017).

91. Zeng, H. What is a cell type and how to define it? Cell 185, 2739 (2022).

92. Puelles, L. et al. The Pallium in Reptiles and Birds in the Light of the Updated Tetrapartite Pallium Model. Evolution of Nervous Systems 1–4, 519–555 (2017).

93. Medina, L. & Abellán, A. Development and evolution of the pallium. Semin Cell Dev Biol 20, 698–711 (2009).

94. Richardson, M. K. Vertebrate evolution: the developmental origins of adult variation. BioEssays 21, 604–613 (1999).

95. Gould, S. J. Heterochrony and the Parallel of Ontogeny and Phylogeny. Ontogeny and Phylogeny %7 7 (1977).

96. Paolino, A. et al. Non-uniform temporal scaling of developmental processes in the mammalian cortex. Nature Communications 2023 14:*1* 14, 1–17 (2023).

97. Cárdenas, A. et al. Evolution of Cortical Neurogenesis in Amniotes Controlled by Robo Signaling Levels. Cell 174, 590–606.e21 (2018).

98. Hodge, R. D. et al. Conserved cell types with divergent features in human versus mouse cortex. Nature 573, 61–68 (2019).

99. Antin, P. B., Yatskievych, T. A., Davey, S. & Darnell, D. K. GEISHA: an evolving gene expression resource for the chicken embryo. Nucleic Acids Res 42, D933 (2013).

100. Blaess, S., Szabö, N., Haddad-Tövolli, R., Zhou, X. & Alvarez-Bolado, G. Sonic hedgehog signaling in the development of the mouse hypothalamus. Front Neuroanat 8, (2015).

101. Cheng, C. W. et al. Zebrafish homologue irx1a is required for the differentiation of serotonergic neurons. Developmental Dynamics 236, 2661–2667 (2007).

102. Baldarelli, R. M. et al. The mouse Gene Expression Database (GXD): 2021 update. Nucleic Acids Res 49, D924–D931 (2021).

103. Cao, J. et al. The single-cell transcriptional landscape of mammalian organogenesis. Nature 566, 496–502 (2019).

104. Bradford, Y. M. et al. Zebrafish information network, the knowledgebase for Danio rerio research. Genetics 220, (2022).

105. Thisse, C. & Thisse, B. High Throughput Expression Analysis of ZF-Models Consortium Clones. ZFIN Publication (2005).

106. Thisse, B. & Thisse, C. Fast Release Clones: A High Throughput Expression Analysis. ZFIN Publication (2004).

107. Hirata, T. et al. Zinc finger gene fez-like functions in the formation of subplate neurons and thalamocortical axons. Dev Dyn 230, 546–556 (2004).

108. Suda, Y. et al. Emx2 directs the development of diencephalon in cooperation with Otx2. Development 128, 2433–2450 (2001).

109. Pikulkaew, S. et al. The knockdown of maternal glucocorticoid receptor mRNA alters embryo development in zebrafish. Dev Dyn 240, 874–889 (2011).

110. Cajal, M. et al. Clonal and molecular analysis of the prospective anterior neural boundary in the mouse embryo. Development 139, 423–436 (2012).

